# Protein length distribution is remarkably consistent across Life

**DOI:** 10.1101/2021.12.03.470944

**Authors:** Yannis Nevers, Natasha Glover, Christophe Dessimoz, Odile Lecompte

**Author notes:** Co-last authors (Christophe Dessimoz, Odile Lecompte).

## Abstract

In every living species, the function of a protein depends on its organisation of structural domains, and the length of a protein is a direct reflection of this. Because every species evolved under different evolutionary pressures, the protein length distribution, much like other genomic features, is expected to vary across species. Here we evaluated this diversity by comparing protein length distribution across 2,326 species (1,688 bacteria, 153 archaea and 485 eukaryotes). We found that proteins tend to be on average slightly longer in eukaryotes than in bacteria or archaea, but that the variation of length distribution across species is low, especially compared to the variation of other genomic features (genome size, number of proteins, gene length, GC content, isoelectric points of proteins). Moreover, most cases of atypical protein length distribution appear to be due to artifactual gene annotation, suggesting the actual variation of protein length distribution across species is even smaller. These results open the way for developing a genome annotation quality metric based on protein length distribution to complement conventional quality measures. Overall, our findings show that protein length distribution between living species is more consistent than previously thought, and provide evidence for a universal purifying selection on protein length, whose mechanism and fitness effect remain intriguing open questions.

## Introduction

The relentless sequencing of whole genomes across the tree of life has revealed an enormous diversity in how evolution has shaped them—be it in terms of resulting genome size [1,2], gene and protein sequence content (e.g. GC content) [3–5], gene length and structure [6,7], and number of protein-coding genes. Indeed, both the size of genomes and coding genes repertoire vary greatly between species. This is especially true among eukaryotes, which tend to have the highest number of genes and the biggest genomes—differences in genome size can reach up to 60,000-fold [2]. In archaea and bacteria the genome size and number of genes are generally correlated, however it is more complicated in eukaryotes, as genome size is mostly impacted by non-coding elements [2]. Other features such as GC content or isoelectric point vary on a gene per gene basis, but studying their distribution at the genome scale has shown that GC content distribution varies considerably across species. This has been associated with adaptation to high temperatures in bacteria, bias in codon usage, and is affected by mutation bias toward GC bases in vertebrates [3]. The distribution of isoelectric points of proteins is more consistent among species [4], with the exception of species living in extreme saline environments [8,9]. While the precise mechanism between the interspecies variation of these variables are not well known, they are probably multifactorial, for example, both GC content [10] of protein-coding genes and isoelectric point of proteins[4] have been shown to be to some extent linked to protein length.

Similarly, the global distribution of protein length can vary between species which evolved under different constraints. A protein’s function is directly dependent on its 3D structure, which ultimately depends on its primary amino acid (aa) sequence and organization into structural domains. A functional protein needs to be long enough to shape itself into structural folds, accommodating one or more functional domains [11] but longer proteins likely have a higher energetic cost (see discussion in [12]). However, the global distribution of protein length within genomes has been scarcely studied, unlike other genome features. The few studies on the subject trace back to the early 2000: an early study [13] reported that protein length follows a similarly shaped distribution in the species sequenced at the time — described as either a gamma or log-normal distribution — and that proteins were smaller on average in prokaryotes than in eukaryotes. A second study confirmed the divergence between eukaryotic and prokaryotic protein length, and noted that protein length distribution was generally consistent within Domains of Life [14]. Another study reported similar results when comparing eukaryotic and prokaryotic orthologous proteins [15]. A more recent study [16] aimed to revisit these analyses by including more species with a higher taxonomic diversity (1,442 species). The authors confirmed the previous observations but reported that the shape parameter of the distribution was not uniform across species, with up to a two-fold difference. Within eukaryotes, they reported smaller proteins in plants and longer proteins in unicellular eukaryotic species.

All of these previous studies focused on the relative differences of protein length between clades. They relied on summary statistics, and did not attempt to explore the causes underlying differences or similarities in protein length distributions. Thus, several fundamental questions remain unanswered: how different are empirical protein length distributions across the tree of life? Is the difference between domains due to a complete shift of the distribution, or merely due to an excess of long proteins? How does the variation in protein length distribution compare with the variation in other aspects of genome architecture?

Here, we address these questions by analysing protein length distributions across 2,326 species spanning the three domains of life. We observed a remarkable consistency in the empirical protein length distribution, especially within each domain of life. The near-universality of protein length distribution is particularly striking in comparison with other genomic features, which tend to be much more variable across different species. Additionally, we show that the most divergent exceptions to this observation are likely due to lower genome annotation quality, with annotation errors that escape standard quality assessment methods—which suggests that the true variation in protein length distribution may be even smaller than what we report here.

## Results

### Protein length distribution in the three domains

We used a dataset of 2,326 species, extracted from the Orthologous Matrix (OMA) database [17]. The dataset comprises species from the three domains of Life: 485 eukaryotes, 153 archaea and 1,688 bacteria (full list in Supplementary Table 1). First, we compared summary statistics of protein length to evaluate how it varies between species and clades (Supplementary Table 2). Considering median protein size, proteins are on average smaller in bacteria (270 aa) and archaea (242 aa), compared to eukaryotic proteins (353 aa). Variation in protein length is lower among bacterial and archaeal species (standard deviation 23.3 and 21.3, respectively) than among eukaryotes (standard deviation: 62.5). The higher dispersion in eukaryotes in regard to the median protein length is observed when considering mean and quartiles (Supplementary Table 2). However, the variation is smaller for the first quartile protein length than for the median in both eukaryotes (standard deviation: 44.7) and prokaryotes (22.6 in bacteria and 15.0 in archaea) meaning that most of the variance is due to variation in the distributions of larger proteins, which is consistent with previously observed gamma distributions[13,16].

However, summary statistics are an incomplete reflection of actual protein size distribution. We plotted empirical protein length distributions for a small subset of diverse, well-annotated model species (Figure 1).

**Figure 1.**
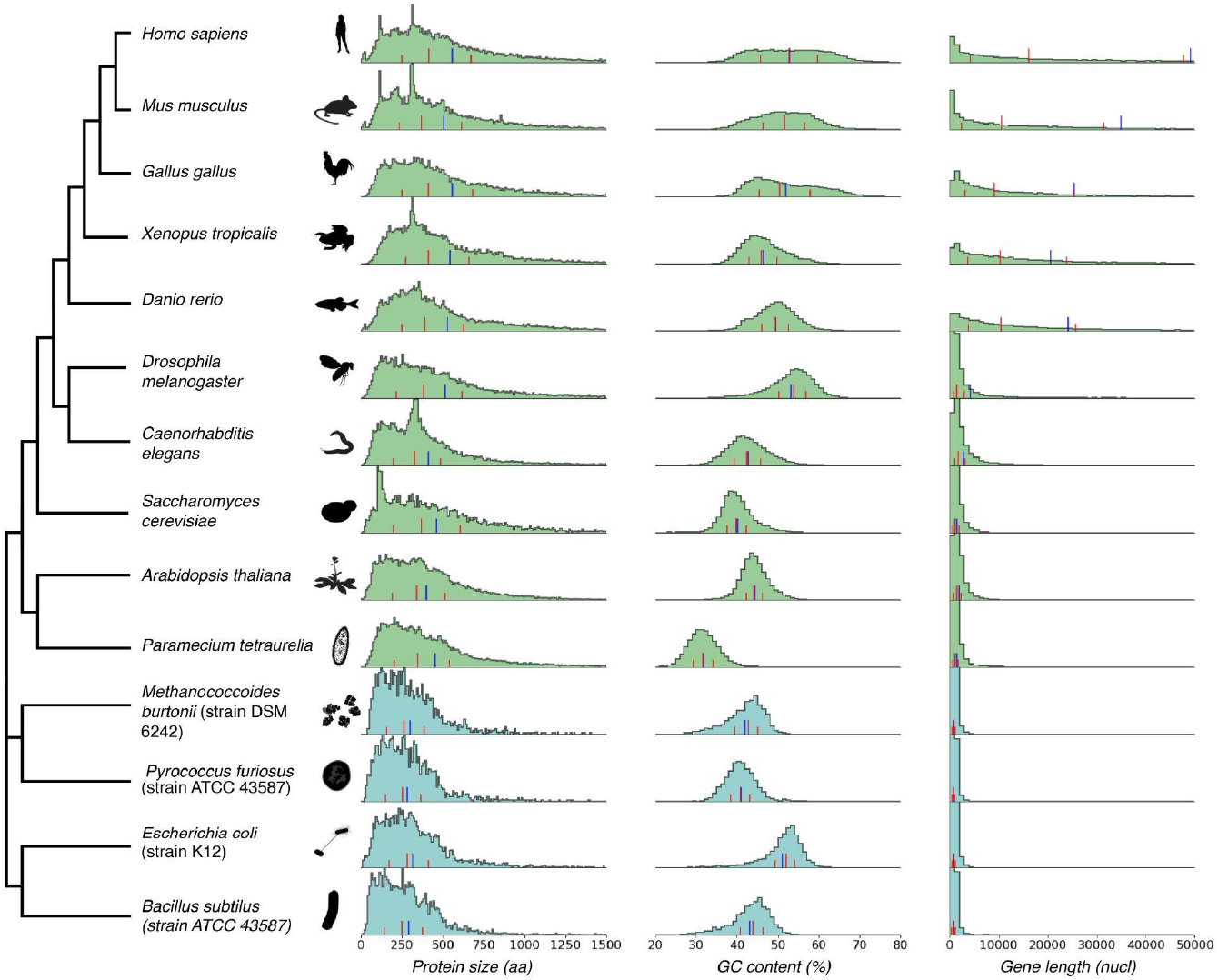
Distributions of protein length, GC content and gene length, for selected model eukaryotic species (light green), bacterial and archaeal species (blue). Summary statistics are shown as lines at the bottom of distribution: red lines indicate first quartile, median and third quartile, and the blue line indicates the mean.

When considering the length distributions of these representative proteomes, a more homogenous picture emerges. Protein size distributions across species greatly overlap, particularly at the left tail and center of the distribution. The right tail — corresponding to larger proteins — shows more variation. Eukaryotes tend to have a higher number of long proteins (>500 aa) than bacteria and archaea. Individual proteome length distributions display peaks at certain protein lengths that appear to correspond to highly duplicated gene families in their respective lineages. For example, the peak of proteins around 320 aa in humans and mice (*Mus musculus*) corresponds to olfactory receptors, a large protein family expanded by gene duplications in mammals [18].

### Protein length is more uniform across species than other genomic features

The similarity of protein length distributions is even more remarkable in comparison with other genomic features, such as the number of protein-coding genes, the number of proteins, the genome length (i.e the size of the genome in base pairs including coding and noncoding sequence), the GC content distribution, and the gene length (including non-coding elements) distribution (Figure 2). We quantified this observation by comparing the variability of protein length distribution with other genomic features across all species in our dataset. To compare scalar features (i.e. features with one global number per genome, such as total genome length) between two species, we used the “Inverted Ratio” (IR; see Methods). An IR close to 0 (in blue) means that the two species have very similar values, whilst an IR higher than 0.5 (in red) represents a more than 2-fold change between species. To compare distributions between two species, we used the Kolmogorov-Smirnov statistic (KS). Briefly, KS is the maximum difference between two cumulative density distributions: it ranges from 0, when the distributions are identical, to 1, when the distributions are so different that they do not overlap.

**Figure 2.**
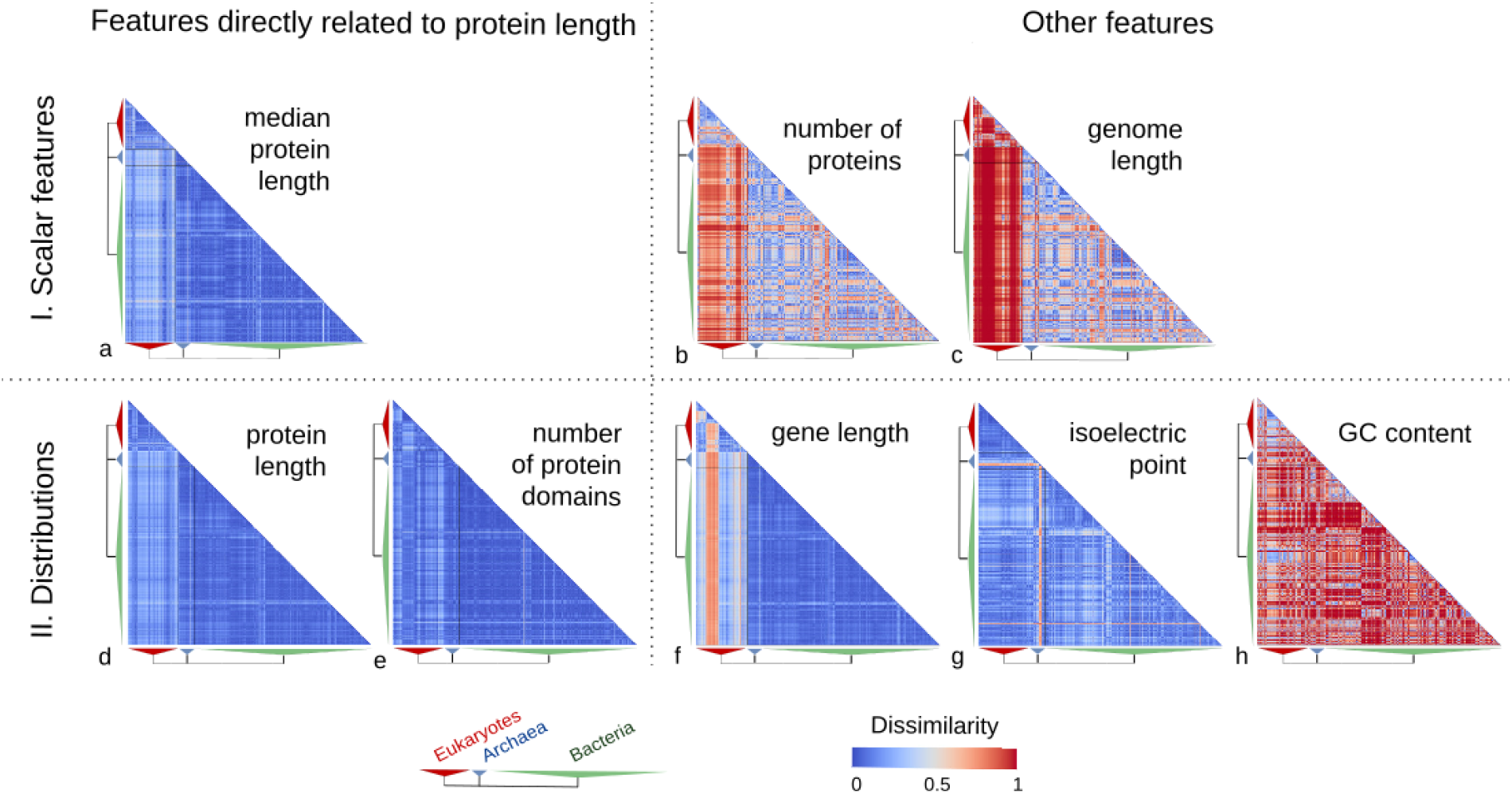
Heatmaps of pairwise species comparison of genomic features. Row and columns are species, ordered by taxonomy. **I**. Heatmaps of dissimilarity of three genomic features for every comparison of species. The dissimilarity measure used is an inverted ratio of the pair. An inverted ratio close to 0, in cool colors, means the compared values are identical or very similar. An inverted ratio higher than 0.5, in warm colors, represents a more than 2-fold difference between the highest and lowest values in the pair. Features compared are: **a**. median protein length, **b**. protein number, **c**. genome length. **II**. Heatmaps of dissimilarity between distributions of gene-centric features for every comparison of species. The dissimilarity measure used is the Kolmogorov-Smirnov statistics. A statistic of 0 (in blue) means complete overlap between distribution and a statistic to 1 (red) no overlap at all, with intermediate ranges between the two extremes. Compared features are: **d**. protein length distribution, **e**. protein domain number distribution, f. gene length distribution, **g**. isoelectric point distribution, **h**. GC content distribution. The heatmaps on the left section correspond to variables directly associated with protein length.

These quantitative measures confirmed the small observed difference in protein length across all species pairs, be it in terms of the median (mean IR: 0.15; Figure 2a, Supp. Figure 1) or in terms of the entire distributions (mean KS: 0.13; Figure 2d, Supp. Fig. 5). Protein lengths are particularly similar among archaea and bacteria (mean IR: 0.09; mean KS: 0.08). As noted above, the protein length variation is higher between eukaryotes and the other two domains (mean IR:0.20; mean KS:0.13) or within eukaryotes (mean IR: 0.17; mean KS: 0.13). Comparing the distributions of the number of structural protein domains per protein-coding gene yielded similar results (mean KS: 0.10, Figure 2e, Supplementary Figure 6) but with higher similarity between eukaryotes and the other domains (mean KS: 0.14).

In comparison, other features vary considerably more. The number of proteins (Figure 2b, Supplementary Figure 2) can change by several order of magnitudes within the same domain, with similar intra-domain variation in eukaryotes and bacteria (mean IR: 0.42, 0.43 respectively), and slightly lower variation in archaea (mean IR: 0.29). As expected, huge interdomain variations of protein number are observed between eukaryotes and prokaryotes (mean IR of 0.8 and 0.73 between eukaryotes versus archaea and bacteria respectively), compared to smaller variations between bacteria and archaea (mean IR of 0.40).

Genome length (Figure 2c, Supplementary Figure 3), varies also considerably in archaea and bacteria (mean IR: 0.35, 0.43 respectively). In eukaryotes, the inter- and within-domain magnitude of differences is even higher (mean IR: 0.92 and 0.74, respectively).

Gene length includes the untranslated regions (UTR), introns, and exons. As such, it is related to, but not equivalent to, protein length. In archaea and bacteria where UTR are short and there are no introns, the distribution of gene length is as consistent across species as that of protein length (mean KS: 0.07 and 0.08). By contrast, it is highly divergent in eukaryotes (Figure 2f, Supplementary Figure 7). Specifically, gene length distribution varies more (mean KS: 0.35), and with higher intensity (KS from 0 to 1 in extreme cases) within eukaryotes and between eukaryotes and the other domains. Even among eukaryotes, deuterostomes (red line in Figure 2f) diverge highly from all other species. These divergences can be attributed to the intron-exon structure that leads to longer genes in eukaryotes, and particularly in deuterostomes [19].

Likewise GC content (the percentage of guanines and cytosines in the coding gene sequence) distribution is not consistent across domains of Life. GC distributions are similar only within some smaller clades. In each domain, the variation (Figure 2h, Supplementary Figure 8) ranges from a KS of 0 to 1 (i.e. no overlap at all). Barring a few exceptions, GC content distributions are relatively more stable within eukaryotic species (mean KS: 0.55) than within bacteria and archaea (mean KS: 0.76 and 0.71, respectively).

Finally, isoelectric point (pH at which a protein is neutrally charged) distributions do not vary as much as GC content across species (Figure 2g, Supplementary Figure 9). But, as with GC content and in contrast with protein length, isoelectric point is more consistent within eukaryotes (mean KS: 0.10) than bacteria (mean KS: 0.18) and archaea (mean KS: 0.32). The exception was a clade of archaea which deviates particularly from other species in terms of isoelectric point: the Haloarchaea. This is not unexpected as these species are known to reside in extreme pH conditions [8,20].

### Many protein length distribution outliers are explained by quality issues

Despite the high overall similarity in protein length distribution, a few species have a protein length distribution that departs from the canonical one, apparent in Figure 2 as white lines crossing their respective domains. These proteomes have a markedly different shape (Examples in Figure 3a, b and c) and are often found in species taxonomically related to species with canonical distribution. For example, Figure 3d-f shows that while *Drosophila melanogaster* (d) and a non-model species of the same genus (e) have a canonical distribution, other close species like *Drosophila simulans* (f) have a comparatively higher abundance of small proteins. Species with an atypical protein length distribution do not have an obvious biological phenotype in common and given the otherwise consistent protein length distribution among most proteomes, even taxonomically distant ones, we hypothesised that these departures could be artefactual. To test this, we sought to assess the genome and proteome quality of outliers from our pairwise comparison of protein length distribution, defining them as being proteomes with a high mean KS dissimilarity (>0.2) with the species of their respective domains.

**Figure 3.**
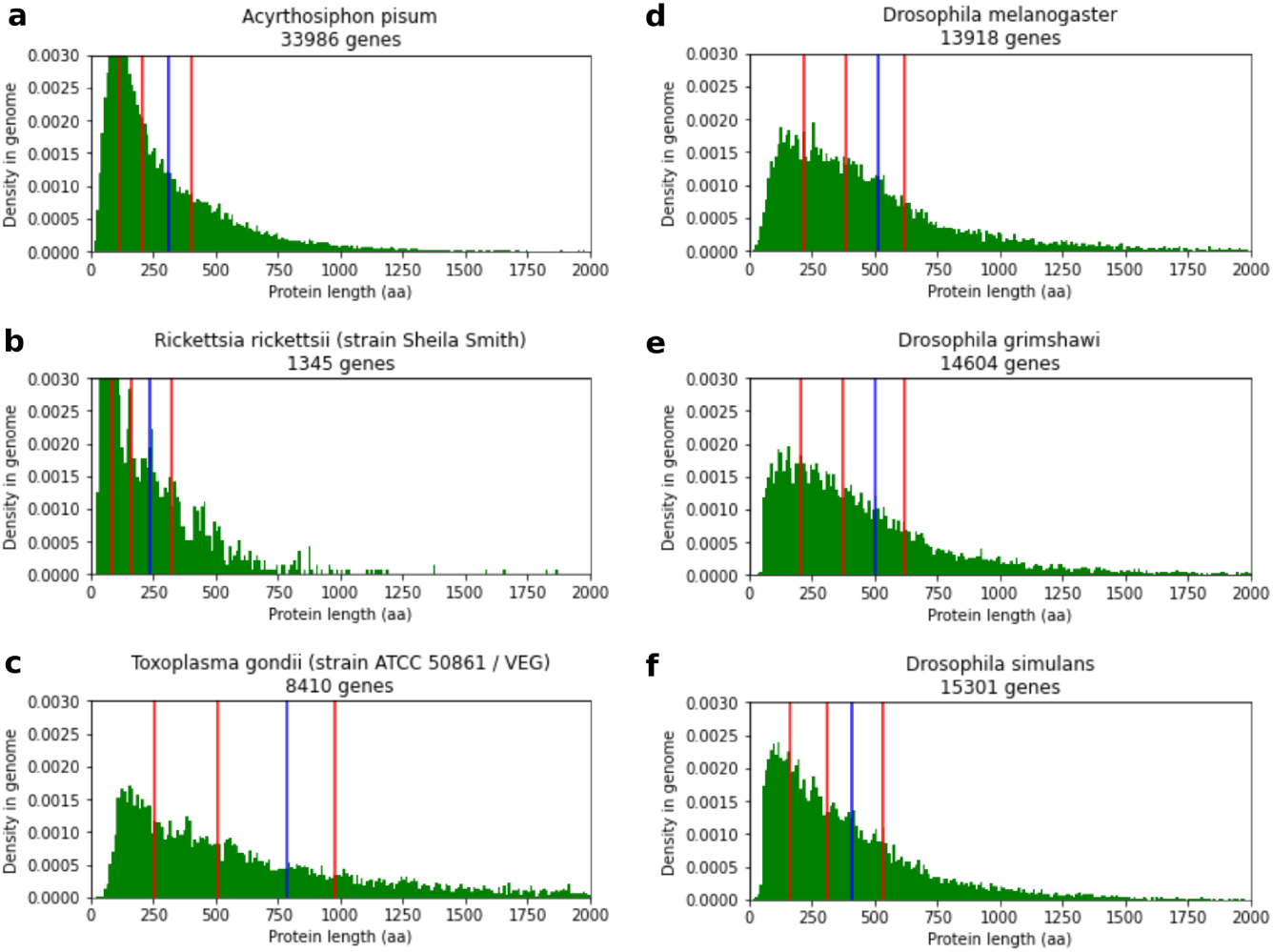
Examples of atypical protein length distributions and distribution heterogeneity between close species. All graphs show the density distribution of protein lengths. The red lines represent the first quartile, median and third quartile of protein lengths, and the blue lines represent the mean. **a-b**. Examples of proteomes with an overabundance of small proteins (eukaryote *Acyrthosiphon pisum* (pea aphid) (a), and bacteria *Rickettsia rickettsii* (b)) **c**. *Toxoplasma gondii*, an example of a proteome with a high proportion of longer proteins. **d-f**. Example of difference in protein length distributions in the *Drosophila* genus. *Drosophila melanogaster* **(d)** has a canonical protein length distribution shape, and similar distributions exist in other *Drosophila* species like *Drosophila grimshawi* **(e)**. *Drosophila simulans*, however, shows a relative abundance of small proteins **(f)**.

Thus we obtained 36 eukaryotes, 3 archaea, and 22 bacteria as outliers. With the exception of 3 eukaryotes with a large tail of long proteins, all the divergent proteomes are characterized by a high peak of proteins in a certain length range, most often small proteins (<100 amino acids). While such distributions were suspected before as being potentially erroneous [16], to our knowledge, this has not yet been demonstrated to be the case. Therefore, we investigated the role of annotation completeness and coding sequence integrity on the occurrence of these outliers.

As smaller proteins are commonly observed in the outlier distributions, it could be an indicator of a high proportion of fragmented protein-coding genes, or incomplete representation of the protein-coding gene repertoire. BUSCO[21] is a commonly used method to assess proteome completeness and fragmentation. We ran BUSCO on our dataset of 2,326 proteomes and flagged all proteomes for which less than 90% of complete BUSCO genes were found (Figure 4): 228 of the 485 (47%) eukaryotic proteomes, 17 of the 153 (11%) archaeal proteomes, and 133 of the 1,688 (8%) bacterial proteomes. In particular, proteomes with an atypical distribution tend to be in the incomplete category: 27 of 36 (75%) eukaryotic proteomes, 2 of the 3 (67%) archaeal proteomes, and 8 of the 22 (36%) bacterial proteomes, meaning that in these cases, low quality genomes may be a cause of the atypical distribution.

**Figure 4.**
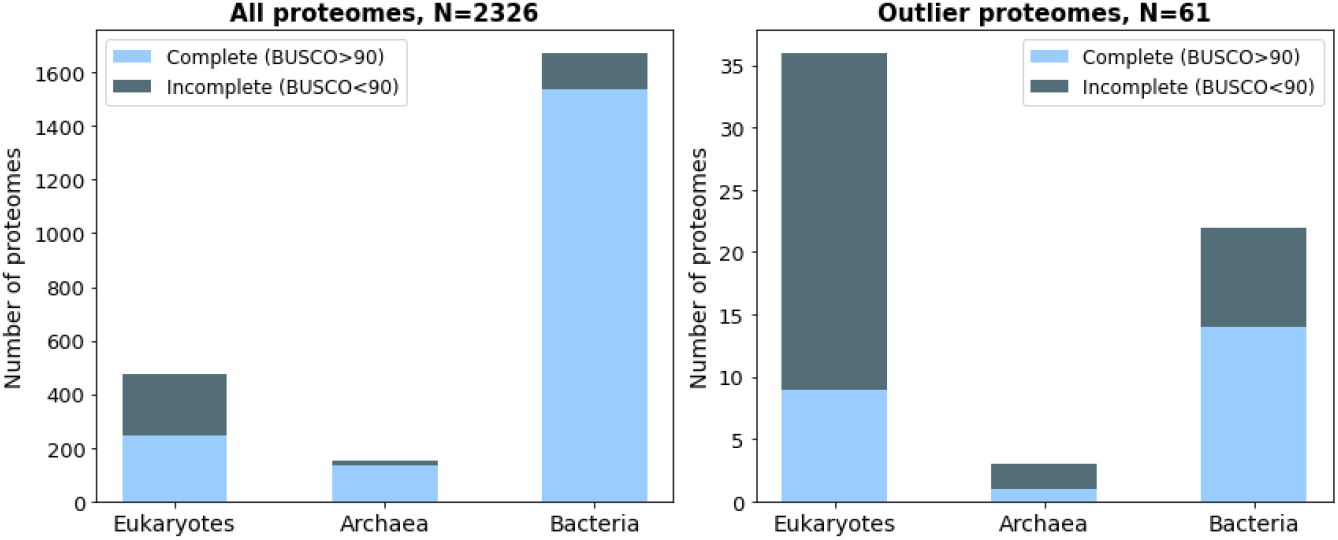
Outlier proteomes in terms of gene length distribution are more likely to be incomplete. Left: Stacked bar of proteomes by domain: mostly complete proteomes in light blue and incomplete proteomes in dark blue. Right: Same representation, with proteomes having the most atypical distribution in regard to their Domain (outlier proteomes).

Still, 24 remaining outliers have mostly complete genomes and few fragments according to BUSCO. We investigated whether these atypical distributions could be due to the biology of these species, or merely annotation artifacts not captured by BUSCO. We checked UniProt [22], RefSeq [23] and the literature for alternative annotation sets (details in *Supplementary Results*), and found seven examples where the annotation set where not consistent and an alternative annotation of the same species had canonical protein length distribution, suggesting these outliers are mainly due to artifactual annotation.

Three eukaryotic species (the fungal plant pathogen *Ustilago maydis*, and the protozoan obligate parasites *Toxoplasma gondii* (strain VEG) and *Hammondia hammondi)* have diverging distributions characterized by a relatively high amount of proteins longer than 500 amino acids and no overrepresentation of small proteins. In-depth analysis of these proteomes (details in *Supplementary Results*) suggested possible but not conclusive link to their lifestyles in the case of apicomplexa.

Combined together, these results suggest that many of the genomes with atypical protein length distribution are characterized by an excess of small proteins, which may be explained in many cases by annotation artifacts — not all of which are captured by conventional quality measures. As for the few which had an excess of larger proteins, we did not find any evidence of artifact, and they all happened to be parasites.

## Discussion

We showed that the distribution of protein size is remarkably consistent within and across the three Domains of Life, particularly in comparison with other genomic features. Moreover, the exceptions appear to be largely caused by genome annotation artefacts.

While staying within tight boundaries, eukaryotic proteins are noticeably longer than both archaeal and bacterial proteins, which is mostly due to a higher proportion of proteins over 500 amino acids. Other studies have shown that this holds true even when comparing orthologous genes [15,24]. It is not clear what evolutionary forces drove eukaryotic proteins to be longer; but it may be associated with the higher modularity of eukaryotic proteins [11], their adoption of alternative splicing [25], and an extension of their chaperone (proteins associated with protein folding) repertoire [26].

The few atypical proteomes characterized by an excess of long proteins are found in the *Ustilago* fungal genus and the Apicomplexa phylum, which are both characterized by a partly intracellular parasitic lifestyle. While this excess could be artifactual — and indeed a previous study has suggested that many gene models in Apicomplexa may be erroneously long [27] — it could also be explained by biological particularities. For instance, proteins directed to the apicoplasts, an organelle specific to the Apicomplexa, typically have signal extensions that make the proteins longer. These proteins are especially long in *T. gondii* [28] — one of the species in our dataset. Second, the process of host-cell invasion in these species rely on proteolytic processing of long protein precursors, located in other organelles specific to the Apicomplexa [29]. This includes proteins involved in host-parasite adhesion [30], which could explain the enrichment of these functions in the longer genes of *Plasmodium falciparum*. While relatively few proteins are well characterized as being part of this process [31], the existence of long protein precursors of smaller functional proteins in the genomes of these species may partly contribute to the observed bias in length distribution

In contrast, most atypical protein length distributions featured an abundance of small proteins (< 100 aa). Small proteins are known to be involved in important biological processes [32] and are generally under-annotated [33], however, our analysis suggests that these outliers stem from annotation artifacts [16]. First, none of the well-annotated model species display an enrichment in small proteins. Second, proteomes with high numbers of small proteins were more likely to be incomplete or fragmented. Third, for a given species, different annotation sets have different proportions of small proteins. For instance, the recent reannotation of the *Daphnia pulex* genome [34] showed that the high number of small proteins in the previous genome is likely spurious. These errors are possibily due to fragmented assembly leading to genome annotation errors [35,36] or by the notorious difficulty to discriminate coding and non-coding ORF [37,38] The overabundance of spurious protein is likely to bias all downstream analyses involving them, leading to an inflated genome size [34,39], an inflated number of orphan genes [40], and errors in orthology inference (See the ‘Addressing Proteome Quality’ section in [41]) but the proportion of small proteins is generally ignored when providing a new annotation set. We propose that the distribution of protein length be used as a new criterion of protein-coding gene quality upon publication, to complement existing quality measures.

The universal character of protein length distribution across the tree of life suggests strong, universal purifying selection that keeps a high proportion of the coding sequence between 50 to 500 amino acids. This force does not act uniformly across all proteins, as the length of known active proteins range from two amino-acids peptides [42] to more than 30,000 aa [43] but our observations support that it does have an effect at the proteome level. The limitation of protein size can be viewed as a simple stochastic process linked to the nature of the genetic code: for any random sequence of codons, the probability of not encountering a stop codon decreases exponentially with length [44]. A similar phenomenon happens due to random mutations — the longer the protein, the higher the chance of accumulating deleterious mutations [45]. Thus, all else being equal, the maintenance of long proteins requires stronger purifying selection. A second factor may be the higher production costs of longer proteins: a longer coding sequence implies increased costs of protein synthesis[12], transcription [46], splicing (in eukaryotes) [47], translation, and chaperone-mediated folding. The costs differential at the level of individual proteins, however, are likely to be negligible and it is unclear what the impact of gene length would be on fitness at the organism level. Finally, recent studies have shown that housekeeping genes tend to be shorter than other transiently expressed genes [45], making them easier to regulate than longer genes, the timing of protein production may then be another parameter affected by selection. Overall, it is unclear which forces contribute to the exponential decrease of proteins toward longer sizes, but it likely results from a mix of several factors.

While evolution might favor shorter proteins overall, the observed gamma distribution (rather than a decreasing exponential shape) implies the existence of other factors favoring protein within the 50-500 length range. The modular organization of proteins into structural domains [48] and the stabilization of proteins by folding may explain this [44], as our data suggests that the distribution of the number of protein domains is even more consistent than protein size across species. Thus, accounting for the average length of a protein domain (100 amino acids [49]), it reflects that most functional proteins are composed of 1 to 5 protein domains [11]. This modular organisation may allow for sufficient diversity to cover the majority of the basic cellular needs, the rest of the functional space could be covered by fewer of smaller and longer proteins able to carry specific function (i.e structural cell organization in the case of the titin), as well as by the modularity of the proteins themselves into protein complexes.

A full exploration of these hypotheses is beyond the scope of this article, but our results invoke intriguing questions about the constraints governing the evolution of gene repertoire. They open the path to future studies to explore the reasons for such an unexpected consistency and provide an operational framework for developing a metric based on protein length distribution to assess the coherence of gene annotations.

## Methods

### Dataset acquisition

Data regarding genomic features of all species were extracted from the August 2020 (All.Aug2020) release of the OMA Database [17]. It consists of 2,326 species: 485 eukaryotes, 153 archaea, and 1,688 bacteria. Genomic and proteomic data available in OMA are from different databases, whose origin can be found on the release page. Genomic features were extracted from OMA as described below:

- **Number of proteins**. We counted the number of protein-coding genes in each species’ proteomes.
- **Genome length**. This data is not available in OMA and not easily obtainable due to the heterogeneity of different data sources. We estimated the genome size by adding for each chromosome or contig, the difference between the 3’-most position (either starting or ending position) of the 3’-most genes and the 5’-most position (either starting or ending position) of the 5’-most gene. This is an estimate that systemically underestimates the real genome length, but is likely to be of a similar order of magnitude.
- **Median protein length**. Median of the protein length of every unique protein in the genome, selecting only one isoform in case of alternative splicing (see below).

#### Isoform and distribution acquisition

All distributions used in this analysis were obtained using one representative protein sequence per protein-coding gene, selecting the main isoform in OMA. These representative isoforms were selected as described in Altenhoff et *al* [17], as the isoform with the highest sequence match compared to orthologous sequences across all species. For each gene, the values for the gene-centric metrics were obtained as follows:

- **Protein length**: The length of the string representing the amino-acid sequence of the protein stored in OMA.
- **Gene length**: The difference between the 3’-most position of the gene and the 5’-most position of the gene, as sorted in OMA. These positions account for untranslated regions.
- **Number of protein domains**: The count of the number of domains as stored in OMA, obtained from the Gene3D [50] database (see Altenhoff et al [51]).
- **GC content**: Proportion of Guanine and Cytosine in the cDNA sequence, as stored in OMA.
- **Isoelectric point**: The isoelectric point is equal to the pH at which a protein is neutrally charged. It was computed from the protein sequence in OMA, using the Biopython package [52].

### Species pairwise comparisons (Heatmaps)

#### Discrete pairwise comparisons (Inverted Ratio)

The pairwise comparisons (IR) between discrete values (protein number, genome length, median protein length) was computed using the formula:

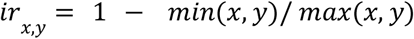

where x is the value in species 1 and y is the value in species 2.

The score is 0 when the values are equal in both species, and goes closer to 1 the more they diverge.

#### Distribution pairwise comparisons

The pairwise comparisons between genewise distributions (protein length, gene length, number of protein domains, GC content, isoelectric comparisons) were done using the two-sample Kolmogorov-Smirnov (KS) statistic. The statistic is computed according to this formula:

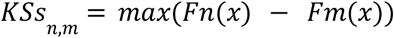

where Fn and Fm are the two compared empirical cumulative distributions. All KS statistics were computed using the SciPy Python package [53].

For the protein length comparison, the mean KS statistic was computed for each species (row), including only comparisons with species of the same domains (columns).

### BUSCO runs

We computed BUSCO [21] statistics on the whole proteome dataset.

First, we generated a FASTA file for each proteome. Then we ran BUSCO (version 4.1.4) on every individual proteome, using the most specific odb10 reference set for this species. This was determined automatically by mapping datasets to the NCBI taxonomic ID in each species lineage. Indication of the BUSCO set used for each species, as well as all statistics are available in Supplementary Table 1.

We divided all proteomes into two sets: complete or incomplete proteomes, based on the number of genes with a complete BUSCO score (not missing nor fragmented). “Complete” proteomes were those with at least 90% of their genes found as ‘Complete’ BUSCO. The complete set was composed of 1,942 species, including 253 eukaryotes, 136 archaea and 1,553 bacteria. The incomplete set (<90% complete BUSCO) was composed of 228 eukaryotic, 17 archaeal, and 133 bacterial proteomes.

### Outlier proteomes definition

We defined proteomes as outliers in regard to their protein length distribution on the basis of the global pairwise comparisons of protein length distribution (see ‘Distribution pairwise comparisons). Specifically, we labeled all proteomes with an average KS with the species of their respective domain higher than 0.2 as outliers.

### Third-party proteomes acquisition

For 24 species, with high distribution divergence (Supplementary Table 3), we manually queried two sequence databases: Uniprot [22] and RefSeq [23], and downloaded the reference proteome for the same species if available. All proteomes were last downloaded in March 2021.

### Summary statistics and analysis

All summary statistics were computed from the data using the Numpy [54] (v. 1.19.0) Python module. Figures were made using the Seaborn [55] (v. 0.11.0) and Matplotlib [56] (v. 3.3.2) Python module.

All code was run with Python v. 3.7.7. A curated version of the code is available as a Jupyter Notebook. (Supplementary Data)

### Gene Ontology enrichment analysis of long genes

We investigated the functional representation of long genes in proteomes with outlier distribution characteristized by an abundance of long proteins. The analysis was carried out for *Ustilago maydis* and all representatives of the Apicomplexa clade in the dataset. Sequence data and GO annotations were extracted from OMA. One species (*Hammondia hammondi*) had no existing GO annotation, thus proteins were automatically annotated using the ‘Gene Ontology Functional Prediction’ of the OMA browser. Briefly, genes from *H. hammondia* were mapped to their closest sequence in the OMA database, and then GO terms were propagated to them from genes in the same Orthologous Group [51,57].

All enrichment analyses were run in Python using the goatools library [58]. For each species, the enrichment procedure was performed using all genes from that species with a protein size greater than different length thresholds (1000, 2000, 3000, 4000, 5000 aa) as study sets. Two background populations were used: either all genes from that species or all genes from the 25 Apicomplexans in the OMA database of the same length requirement as the foreground population. Only GO terms enriched with a Bonferroni-corrected p-value<=0.05 were considered as significant. Results were plotted and visualized using the Go-Figure software [59].

The analysis described here was run with Python v. 3.7.7. A curated version of the code is available as an independent Jupyter Notebook (Supplementary Data).

## Author contributions

YN, CD and OL designed the study. YN carried out data extraction, comparative analyses of genomic features and primary analysis of outlier proteomes. NG carried out in-depth analysis of outlier proteomes with abundance of long proteins. YN and NG drafted the manuscript. NG, CD and OL edited the manuscript. All authors read and approved the final version of the manuscript.

## Acknowledgements

The work was carried out with the support of the IdEx Unistra in the framework of the “Investments for the future” program of the French government and Institute funds from the Centre National de la Recherche Scientifique and the Université de Strasbourg, as well as the Swiss National Science Foundation (Grant No. 183723). The authors thanks Natalia Zajac for providing critical feedback on an early version of this work.

