## Supplemental Information for "Protein length distribution is remarkably consistent across Life"

### Supplementary materials

**Supplementary Data 1** : Archive containing a graphical representation as png file of protein length distribution for each proteome in the dataset. Available on Zenodo at https://doi.org/10.5281/zenodo.5379545.

**Supplementary Data 2 :** Dataset files and associate Jupyter Notebook for the main analyses of the paper (Species comparison of genomic features, outlier protein length investigation…). Available on Zenodo at https://doi.org/10.5281/zenodo.5379545.

**Supplementary Data 3** : Jupyter Notebook for the gene enrichment in species with outlier distribution characterized by long proteins, and result files of the enrichment. Available on Zenodo at https://doi.org/10.5281/zenodo.5379545.

**Supplementary Table 1 :** Data summary and BUSCO sets

**Supplementary Table 2**: Summary statistics of protein length for genomes in the dataset, by domains.

| | **eukaryotes** | | **bacteria** | | **archaea** | |
|---|---|---|---|---|---|---|
| | Average | Std dev | Average | Std dev | Average | Std dev |
| First Quartile | 216.8 | 41.9 | 161.5 | 22.1 | 145.3 | 12.2 |
| Median | 372.3 | 55.8 | 270.8 | 22.5 | 245.5 | 14.5 |
| Mean | 486.3 | 83.2 | 317.3 | 24.9 | 286.3 | 15.6 |
| Third Quartile | 604.1 | 98.1 | 407.4 | 25.7 | 373.0 | 16.0 |

**Supplementary Table 3** : Complete proteomes with atypical distribution and comparisons with other annotation sets.

**Supplementary results**

- Correlation between genomic features
- Comparison of outliers proteomes with other annotation sets
- Functional analysis of proteomes with abundance of long proteins

### Supplementary results

#### Correlation between genomic features

Within each domain, we computed all pairs of correlations between the three scalar genomic features (median protein length, protein number, genome size). Variation of protein length in regard to the other parameters are mainly uncorrelated (pearson correlation < 0.3). The exception, as noted in a previous study [1], was that median protein length in eukaryotes is inversely correlated to protein number. We find a weak but significant Pearson correlation between these variables (correlation: -0.38, p=1.9e-18). A Spearman correlation indicated the same result, but with an even weaker correlation (correlation: -0.29, p=7.4e-11), meaning that this relation is partly driven by extreme values of certain clades. Plants, for example, have the highest number of proteins and relatively few long proteins (see Supplementary Figure 4 for comparison of proteome size and protein length by clades).

From our main analysis (see main text : *Protein length is more uniform across species than other genomic features*) distribution of protein length, isoelectric point, and GC content appear to follow different patterns across species. At the gene level, however, correlation between these three distinct features has been noted before [2,3]. To check if this relation could be retrieved at the species level, we computed the Pearson correlation coefficient between all pairs of species' mean protein length, mean GC content and mean isoelectric point for all domains. Only isoelectric point and GC content in archaea were moderately inversely correlated (correlation=-0.54, p-value=6.0e-13), with weaker but statistically significant association in bacteria (correlation=-0.30, p-value=4.9e-36) and eukaryotes (correlation=-0.18, 5.3e-05). Protein length was only weakly positively correlated to GC content in bacteria (correlation=0.22, 3.9e-21) and eukaryotes (correlation=0.12, p-value=0.006) though not in archaea, but no significant relation was found between mean protein length and isoelectric point in any domain.

#### Comparison of outliers proteomes with other annotation sets

For the 24 outlier proteomes with high BUSCO scores (see main text : *Many protein length distribution outliers are explained by quality issues*), we checked UniProt [4] and RefSeq [5] for alternative annotation sets. We were able to retrieve 23 annotation sets from UniProt and 21 from RefSeq, with one species for which no additional annotation set was available in both databases. (Supplementary Table 3), at least one of the alternative distributions was much closer to the typical distribution of the species' domain than the one available in OMA, with an average KS<0.15, and would not have been considered as an outlier in the present analysis. In each of these cases, the proportion of small proteins differed between sets (Supplementary Figures 10-34), with higher median protein sizes in the alternative protein sets. The differences are clearly visible in the case of four eukaryotic protein sets (*Amborella trichopoda*, *Acyrthosiphon pisum*, *Brugia malayi, Loa loa* - Supplementary Figures 24 and 26-28): the alternative annotation distributions are much closer to a 'canonical' distribution.

No differences between the retrieved annotation sets does not necessarily mean that the annotations are correct. For example, while few differences were found in the available annotation sets we compared for *Daphnia pulex*, a reannotation of the genome [6] showed significant differences between the new and old annotations. However, the annotation sets from UniProt and from our present dataset are based on the older annotations. In the reannotation study, the authors found a total of 18,440 protein-coding genes in their annotations, instead of the 30,097 previously reported, yet both annotations had a similar high BUSCO completeness score, close to 96%. Discrepancies between the two sets were reported to be mostly due to a high number of small protein-coding genes in the original annotation, that were not retrieved in the new one. This example tends to support the hypothesis that an excess of small proteins in genome annotations is due to the methodology of genome assembly or annotation rather than biological particularity.

Finally, even in the cases where few differences are found between the original and alternative annotation sets, other species that are closely related to the "outlier" species have a protein length distribution close to the canonical one. In particular, the atypically-distributed set of bacteria proteomes is composed of 9 representatives of the *Rickettsia* genus. This could hint at a taxonomic biological specificity of the genus, however other species of the same genus display a "canonical" distribution (Supplementary Figure 35). The fact that inconsistency of the protein length distribution is not verified in the whole clade goes against the hypothesis that these uncommon distributions may be explained by the biological specificities.

#### Functional analysis of proteomes with abundance of long proteins

In contrast to genomes with a relatively high amount of short proteins, three eukaryotic species (the fungal plant pathogen *Ustilago maydis*, and the protozoan obligate parasites *Toxoplasma gondii* (strain VEG) and *Hammondia hammondi)* had diverging distributions characterized by a relatively high amount of proteins longer than 500 amino acids and no overrepresentation of small proteins. Few differences in terms of protein number and length distributions were found between our annotation set and those found on UniProt and RefSeq (although none could be found for *Hammondia hammondi*). Other species in Apicomplexa (clade comprising T*oxoplasma gondii* and *Hammondia hammondi*) - in particular from the *Plasmodium* and *Toxoplasma* genus - and in the *Ustilago* genus displayed similar shape of distributions (large tail of long proteins) (Supplementary Figures 36 and 37), though they were not flagged as outliers in this analysis, likely because their divergence were not as extreme as the aforementioned three.

The possibility of taxonomic-specific biological particularity appears likelier for these species, especially when taking into account that these are all parasitic or pathogenic species. In order to check if the longer proteins were associated with a specific function, we checked the longest protein-coding genes in these genomes for enrichment of Gene Ontology (GO) [7,8] terms (Supplementary Figures 37-39, Supplementary Files). For the background populations, we used two sets of genes: either the entire gene repertoire of a given species, or the entirety of similarly long-size genes in 25 Apicomplexan species. These analyses are complementary since the former informs on functional categories overrepresented in the longest proteins of the genomes, while the latter informs on functional terms that are specific to long genes in the target species.

No enrichment was supported for longer genes in *Hammondia hammondi* using both backgrounds, likely because it only had 115 genes (1.4%) of its genes annotated with GO terms. Significant enrichments were found for *Ustilago maydis* (Supplementary Figures 37) and *Toxoplasma gondii* (Supplementary Figure 38) for proteins longer than 1,000 aa, with both backgrounds. However, the enriched GO terms were mostly generic terms. In particular, *Ustilago maydis* genes were highly enriched over many non-specific categories compared to the "all species' long proteins" background, as exemplified by the most enriched terms (GO:0005515 protein binding, GO:0009987 cellular process, etc). This may reflect that the observed abundance of long genes is not tied to a species-specific feature and that genes in these species are longer than their orthologs in other species regardless of function.

In the case of the *Toxoplasma gondii* GO enrichment using all Apicomplexa long proteins as a background, kinase activity seems important, specifically positive regulation of MAP kinase activity (GO:0043406) and myosin light chain kinase activity (GO:0004687). *T. gondii* contains two MAP kinases, one of which has an increased expression under osmotic stress and during parasite life cycle differentiation [9]. This, combined with the finding that MAPK inhibitors can block *T. gondii* replication [10], suggests that MAP kinases are important for parasite proliferation [11]. Furthermore, myosin light chain kinase was shown to play a key role in the apicomplexan 'gliding' motility, a strategy for penetrating host cells [12].

We extended this analysis to other Apicomplexa, that have a similar, yet less marked, abundance of higher-length proteins. Doing this, we found a significant enrichment ($p < 10^{-10}$) on terms pertaining to host-pathogen interactions in *Plasmodium falciparum* (strain D7) for genes longer than 1,000 amino acids, using either background. For example, 'adhesion of symbiont to host' GO:0044406), 'modulation by symbiont of host erythrocyte aggregation' (GO:0020013), or 'rhoptry' (GO:0020008), which is a specialized apicomplexan organelle important for host invasion [13]. Whilst it is unclear whether it applies to other apicomplexans, these results point to the abundance of long protein-coding genes possibly contributing to the parasitic lifestyle of *Toxoplasma gondii* and *Plasmodium falciparum*. Overall, however it is not conclusive enough to exclude the possibility of annotation artifacts.

### Supplementary Figures

**Supplementary Figure 1**: Distribution of the inverse ratio in pairwise comparisons of median protein length (Heatmap in Figure 2a). a. Global. b. By life domain.

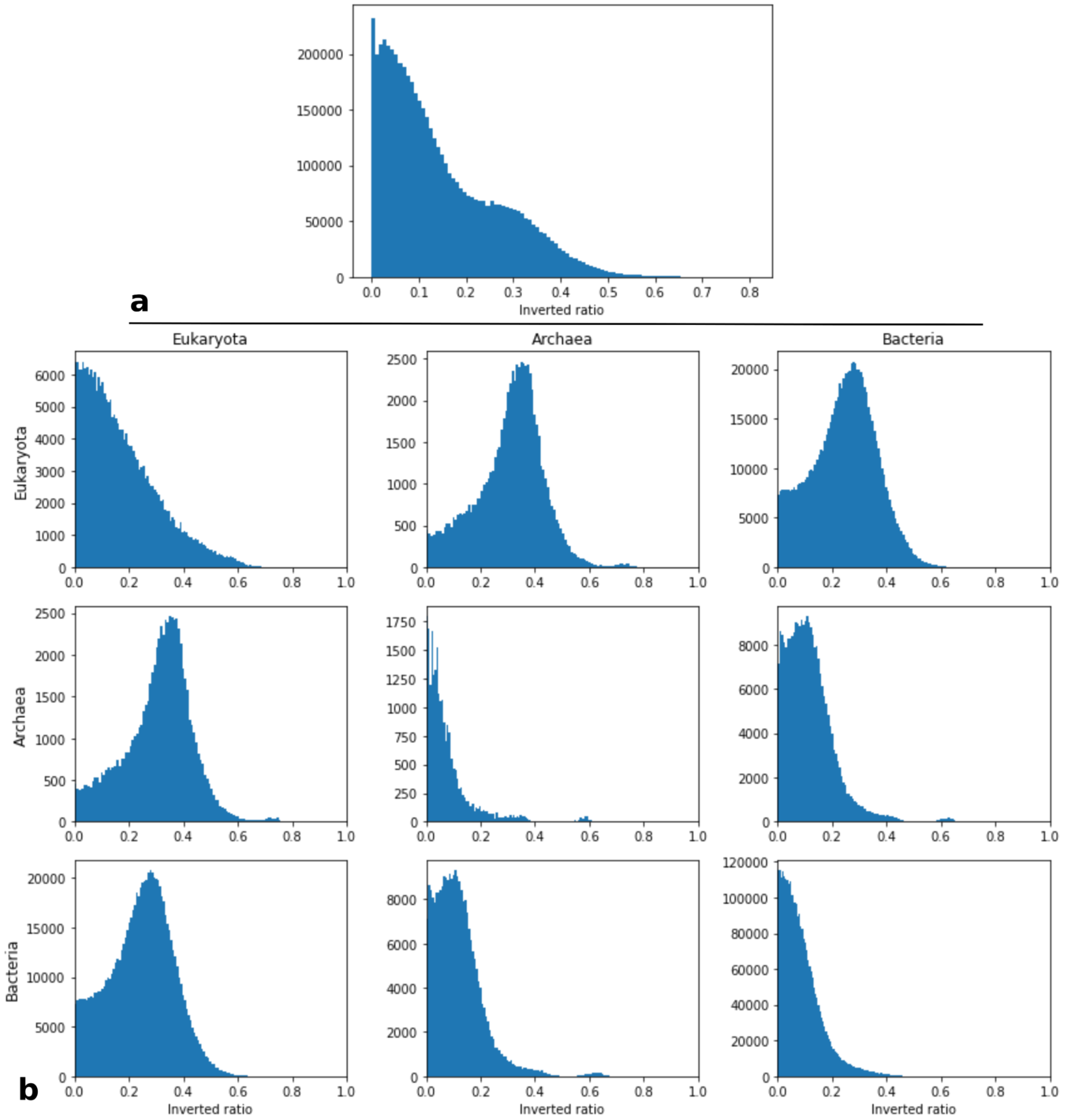

**Supplementary Figure 2**: Distribution of the inverse ratio in pairwise comparisons of number of proteins (Heatmap in Figure 3a). a. Global b. By life domain.

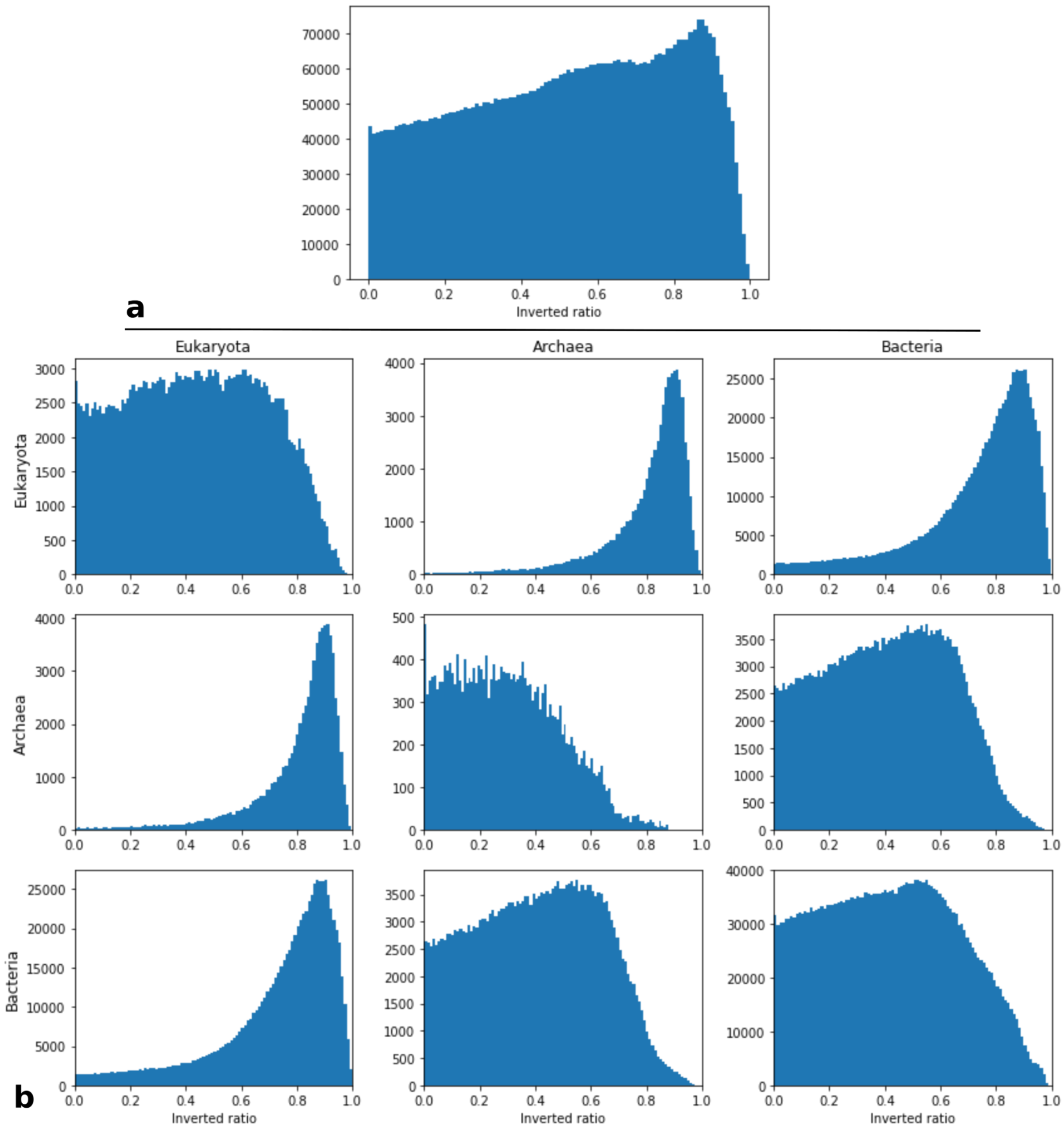

**Supplementary Figure 3**: Distribution of the inverse ratio in pairwise comparisons of genome length (Heatmap in Figure 3a). a. Global b. By life domain

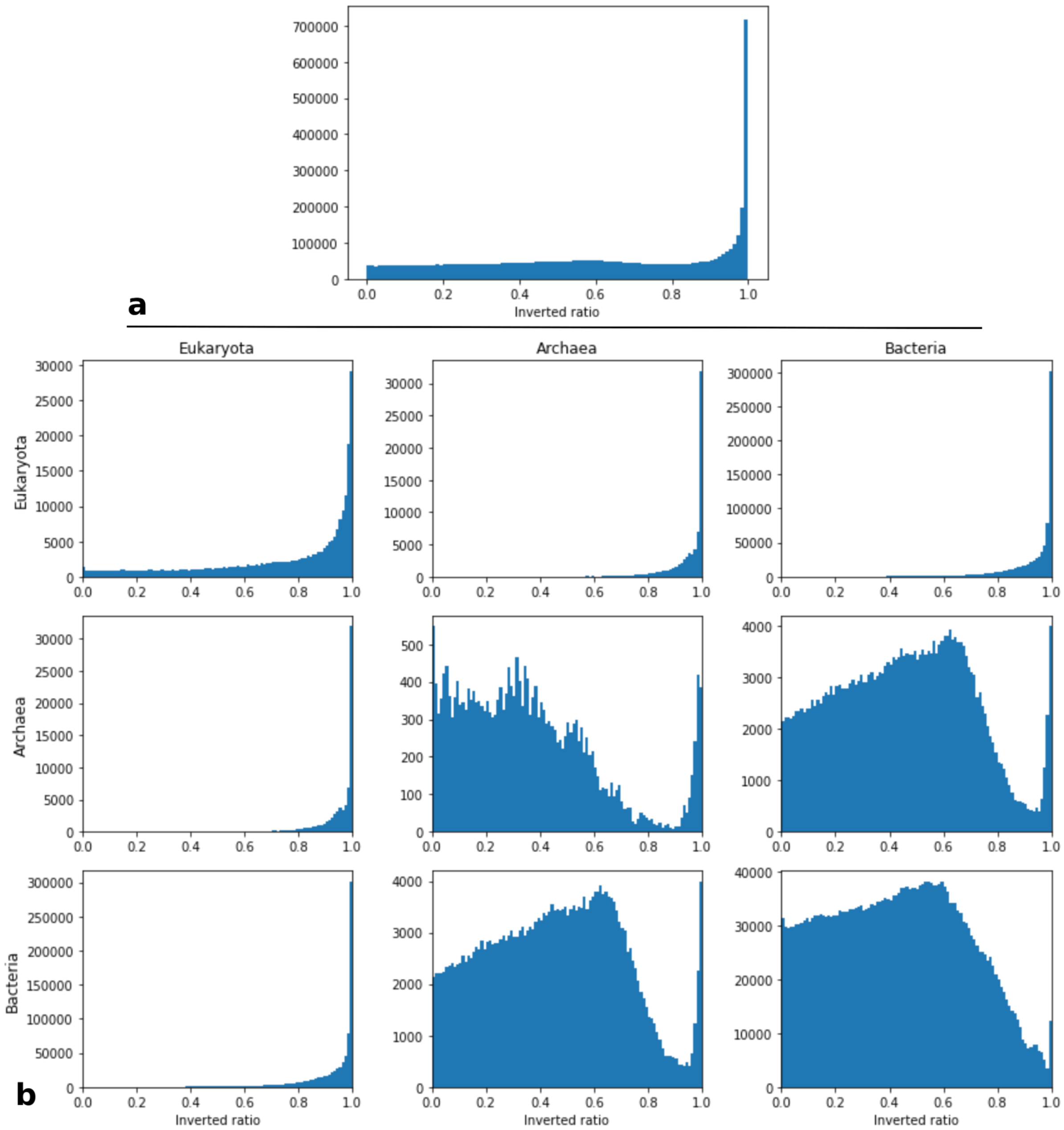

**Supplementary Figure 4 :** Relation between proteome size and protein length. Scatterplot of protein length (x-axis) and number of protein (y-axis). Colors correspond to taxonomic divisions. Archaea and bacteria form a dense cluster with smaller proteins and fewer genes than eukaryotes, which are more diverse regarding both parameters. Plants (in green) tend to have both a higher number of proteins and smaller proteins than other eukaryotes.

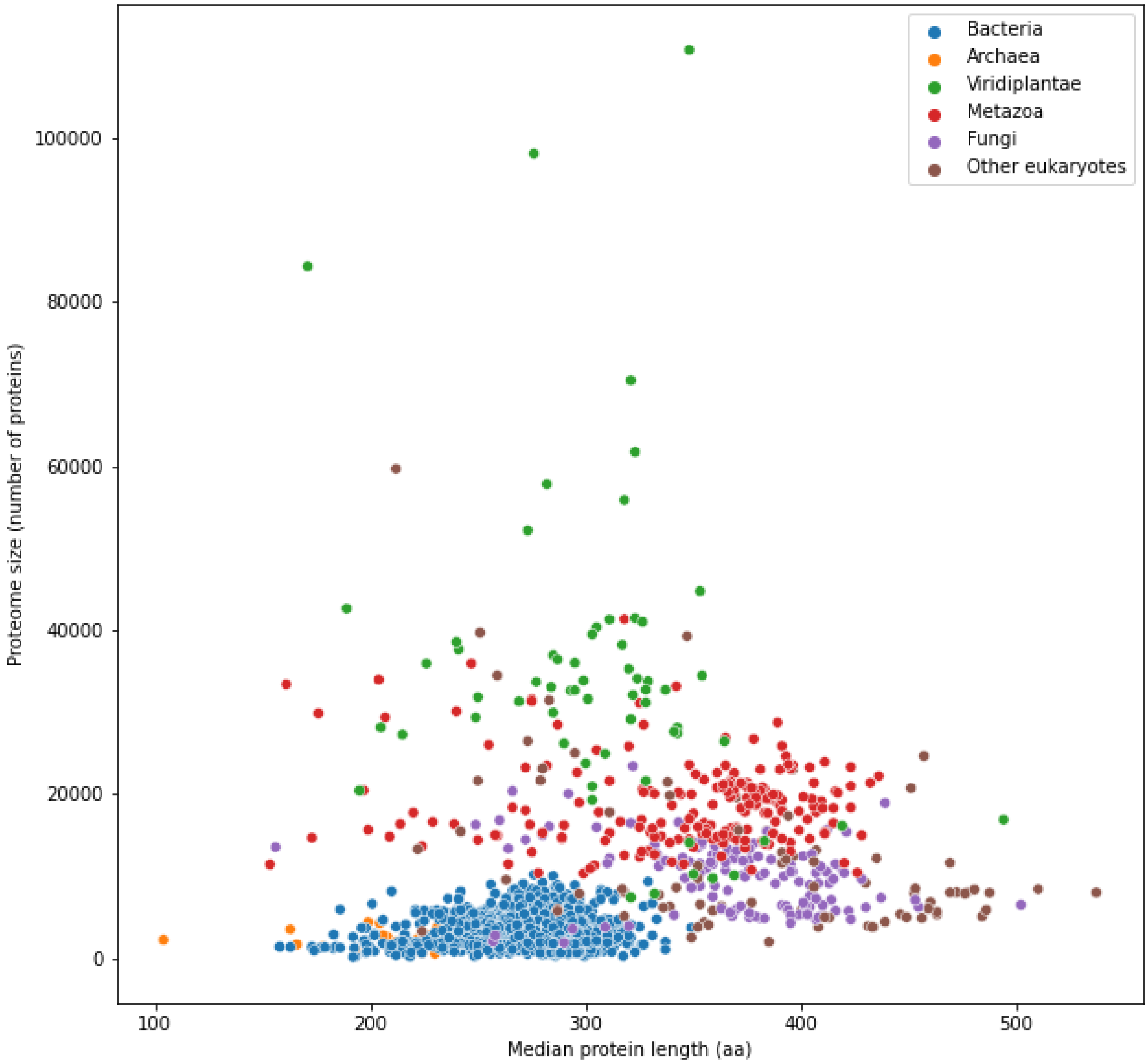

**Supplementary Figure 5**: Distribution of the Kolmogorov-Smirnov statistics in pairwise comparisons of protein length distributions (Heatmap in Figure 3b). a. Global b. By life domain.

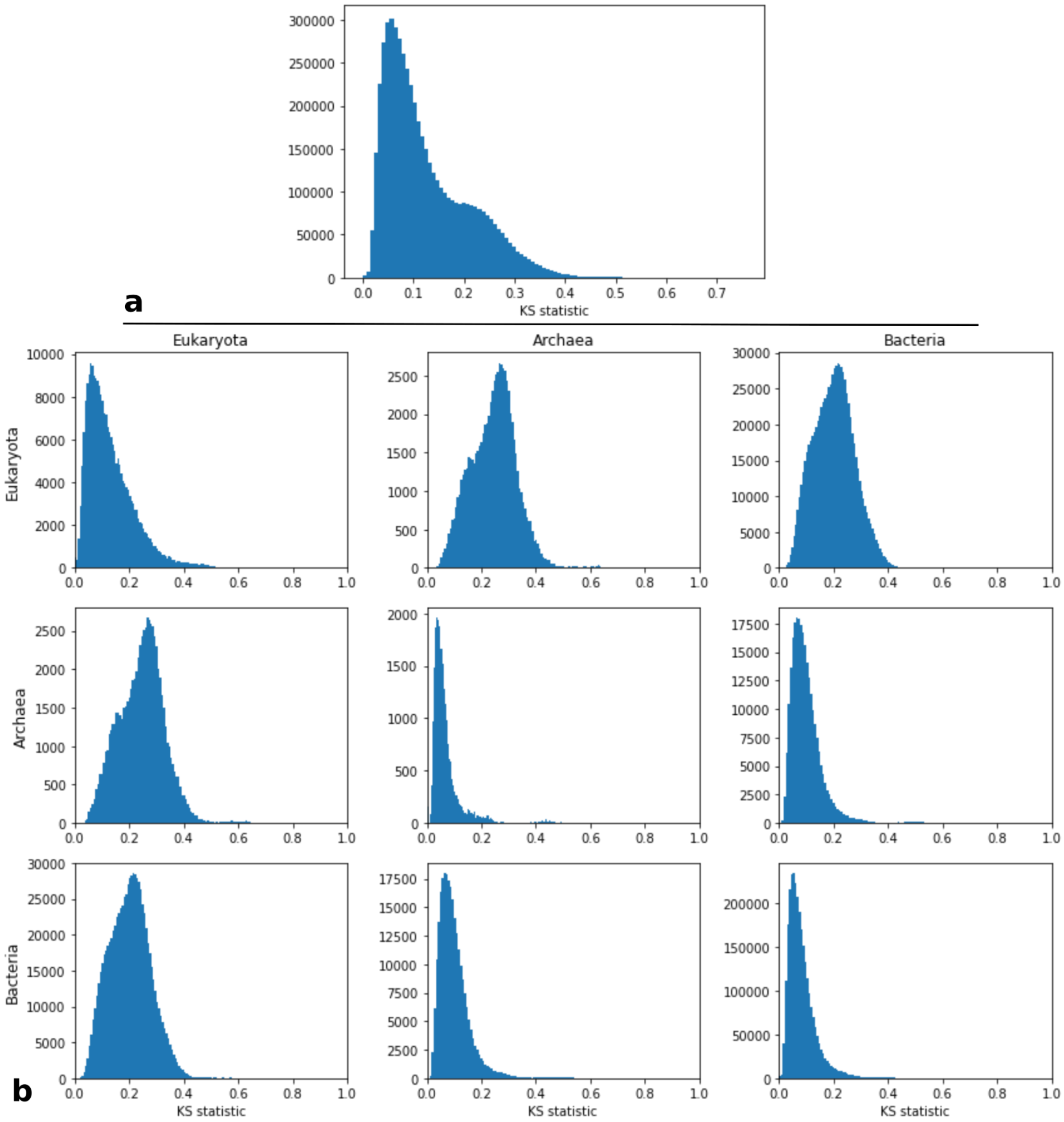

**Supplementary Figure 6**: Distribution of the Kolmogorov-Smirnov statistics in pairwise comparisons of gene length distributions (Heatmap in Figure 3b). a. Global b. By life domain.

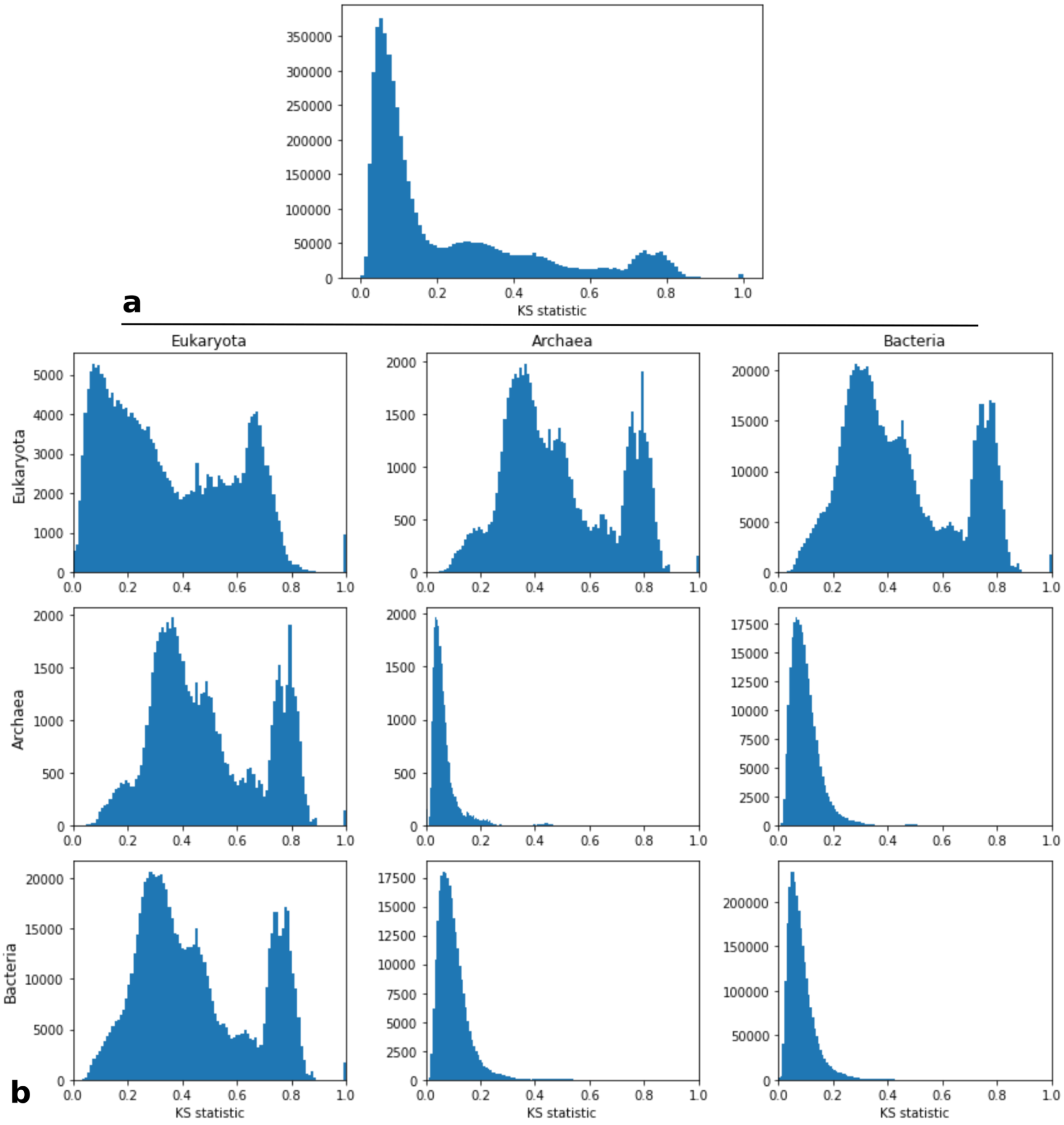

**Supplementary Figure 7**: Distribution of the Kolmogorov-Smirnov statistics in pairwise comparisons of number of protein domain distributions (Heatmap in Figure 3b). a. Global b. By life domain.

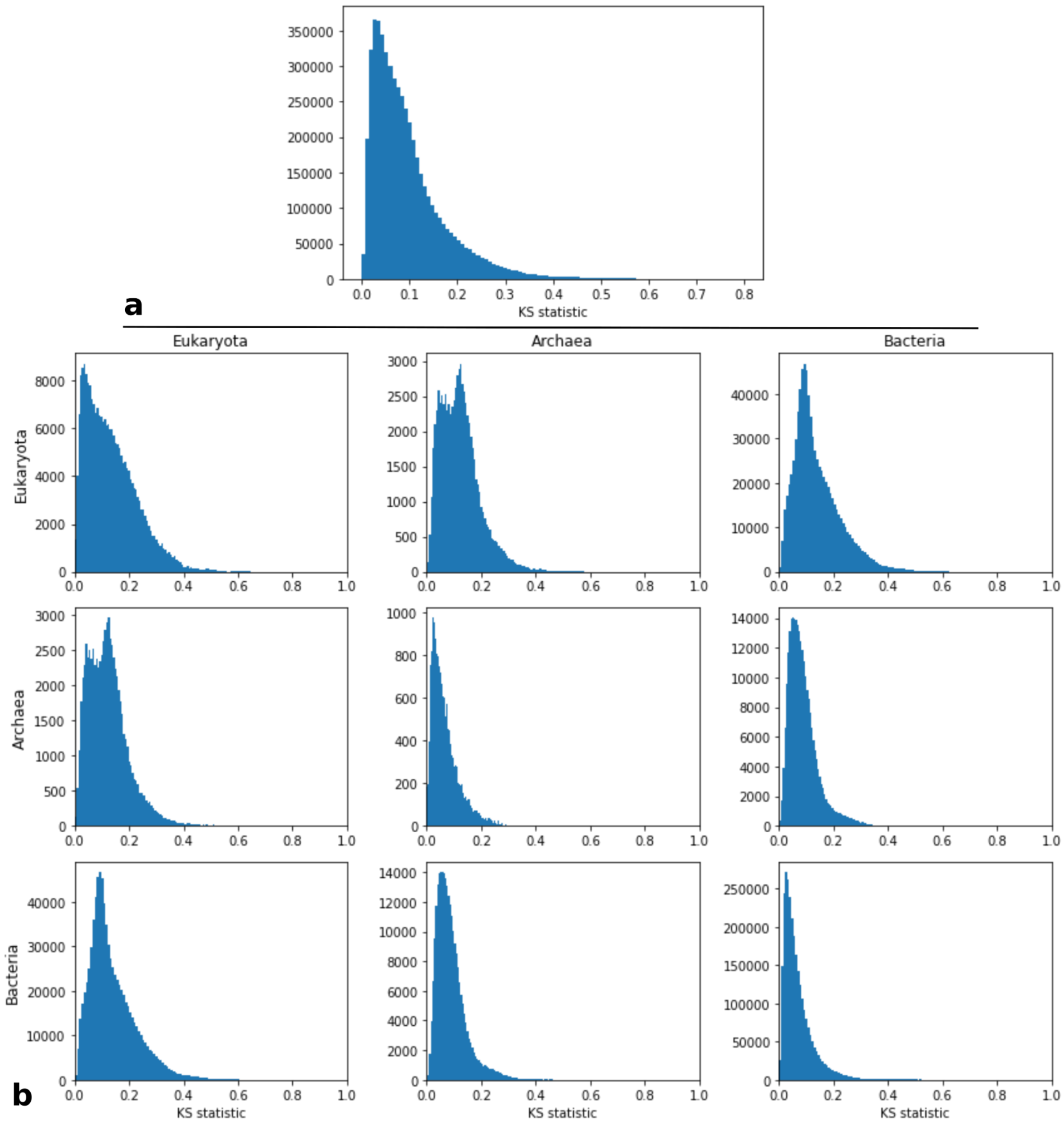

**Supplementary Figure 8**: Distribution of the Kolmogorov-Smirnov statistics in pairwise comparisons of number of gene GC content distribution (Heatmap in Figure 3b). a. Global b. By life domain.

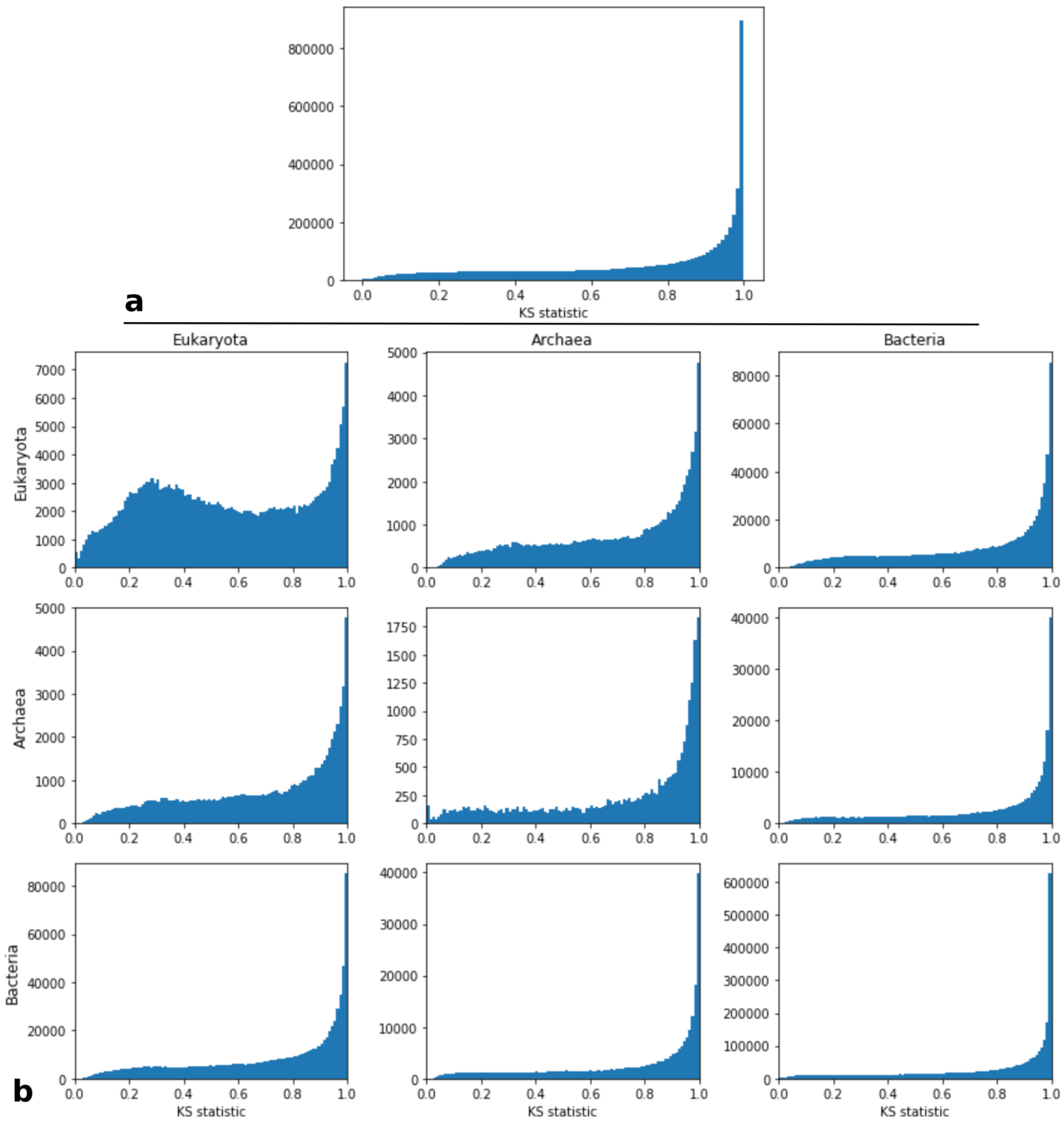

**Supplementary Figure 9**: Distribution of the Kolmogorov-Smirnov statistics in pairwise comparisons of number of protein isoelectric point distributions (Heatmap in Figure 3b). a. Global b. By life domain.

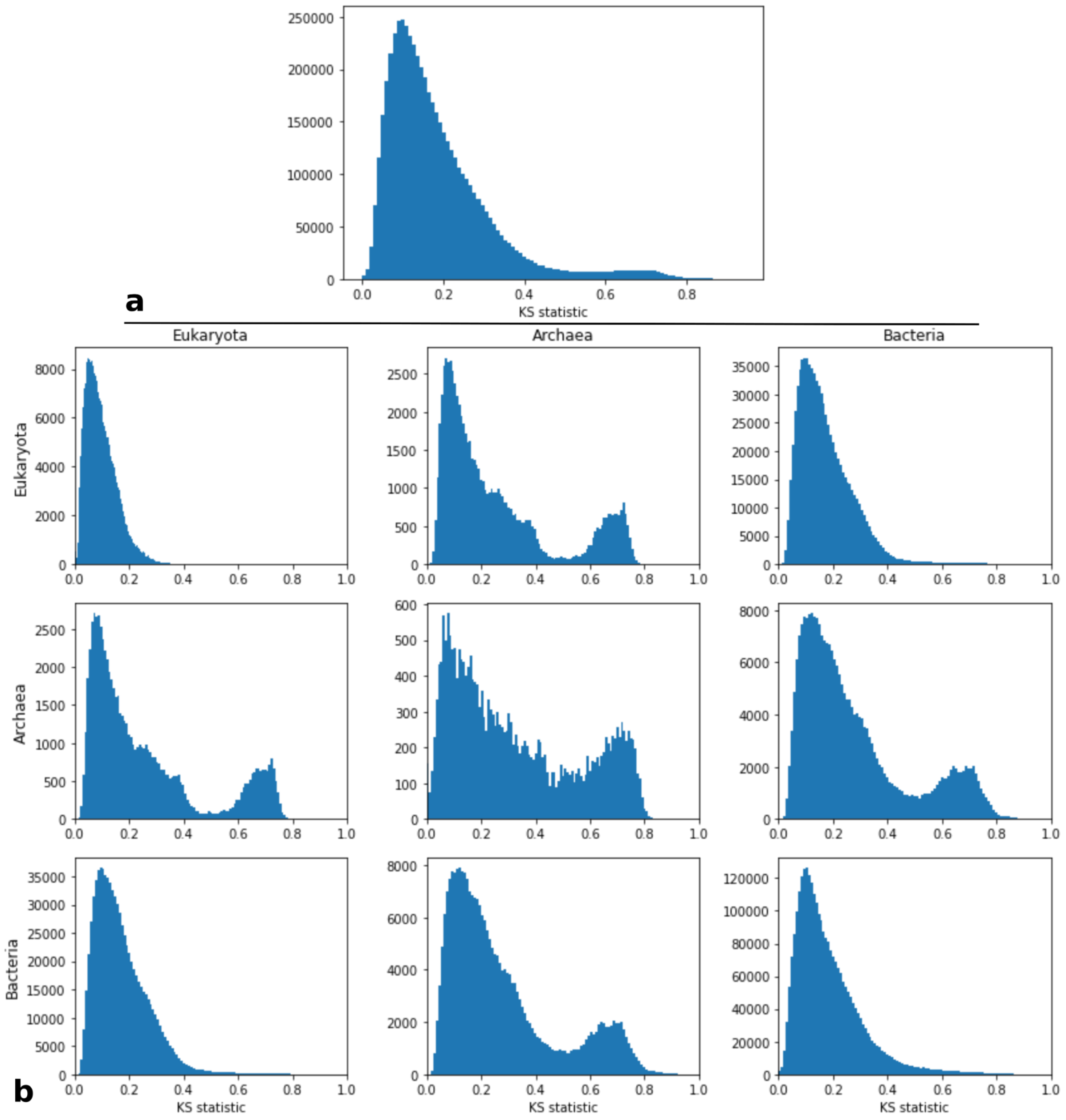


**Supplementary Figures 10-34:** Protein length distribution from different sources for proteomes with atypical distribution, one figure by species. For each outlier species: the protein length distribution of a close species with a canonical distribution (1st from the top), the OMA length distribution (2nd from the top), and comparison with annotation sets from RefSeq and Uniprot (respectively 3rd and 4th from the top). Missing plots mean no annotation set was found in the corresponding database.

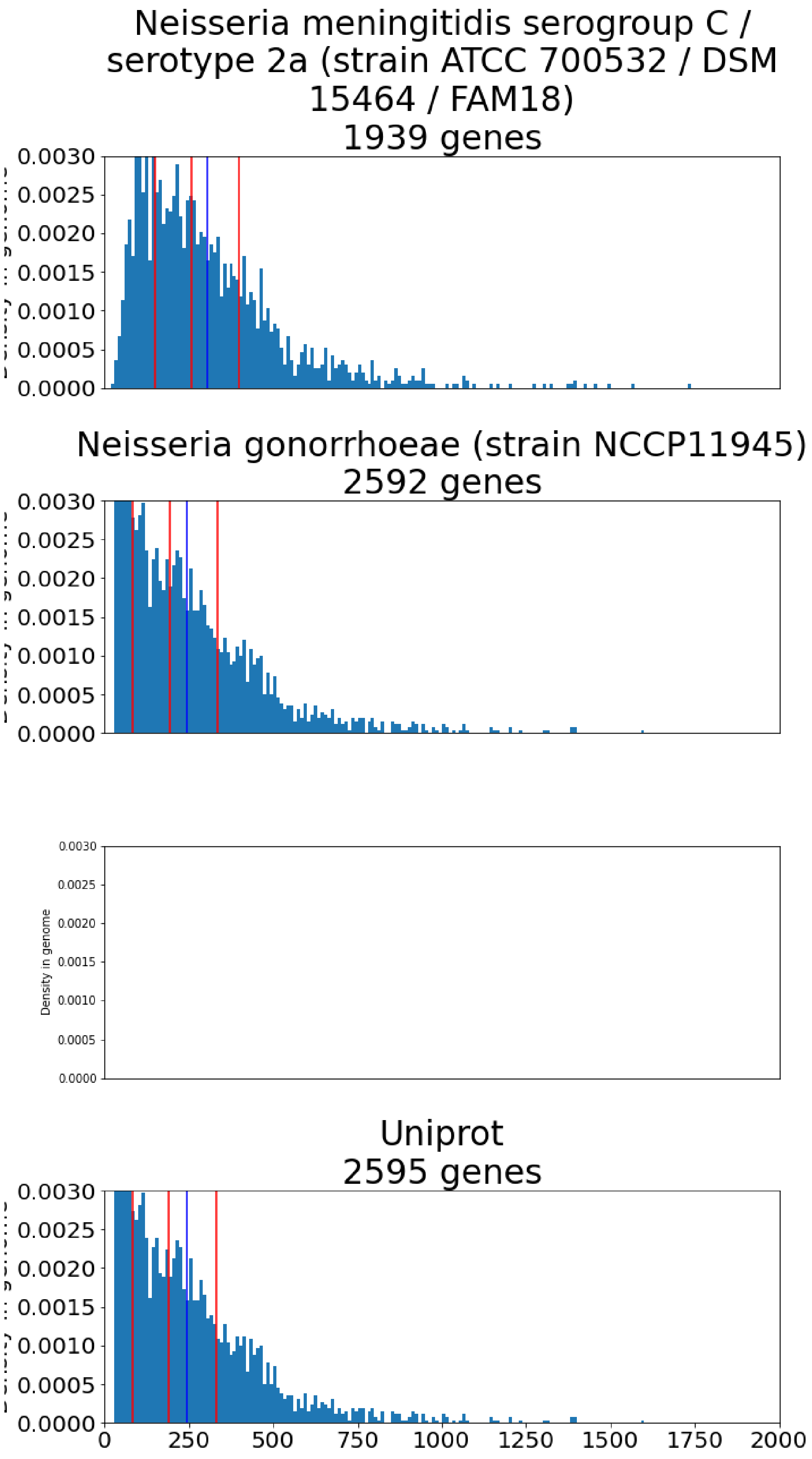
Neisseria meningitidis serogroup C / serotype 2a (strain ATCC 700532 / DSM 15464 / FAM18)
1939 genes
Neisseria gonorrhoeae (strain NCCP11945)
2592 genes
Density in genome
Uniprot
2595 genes
0.0000
0.0005
0.0010
0.0015
0.0020
0.0025
0.0030
0
250
500
750
1000
1250
1500
1750
2000

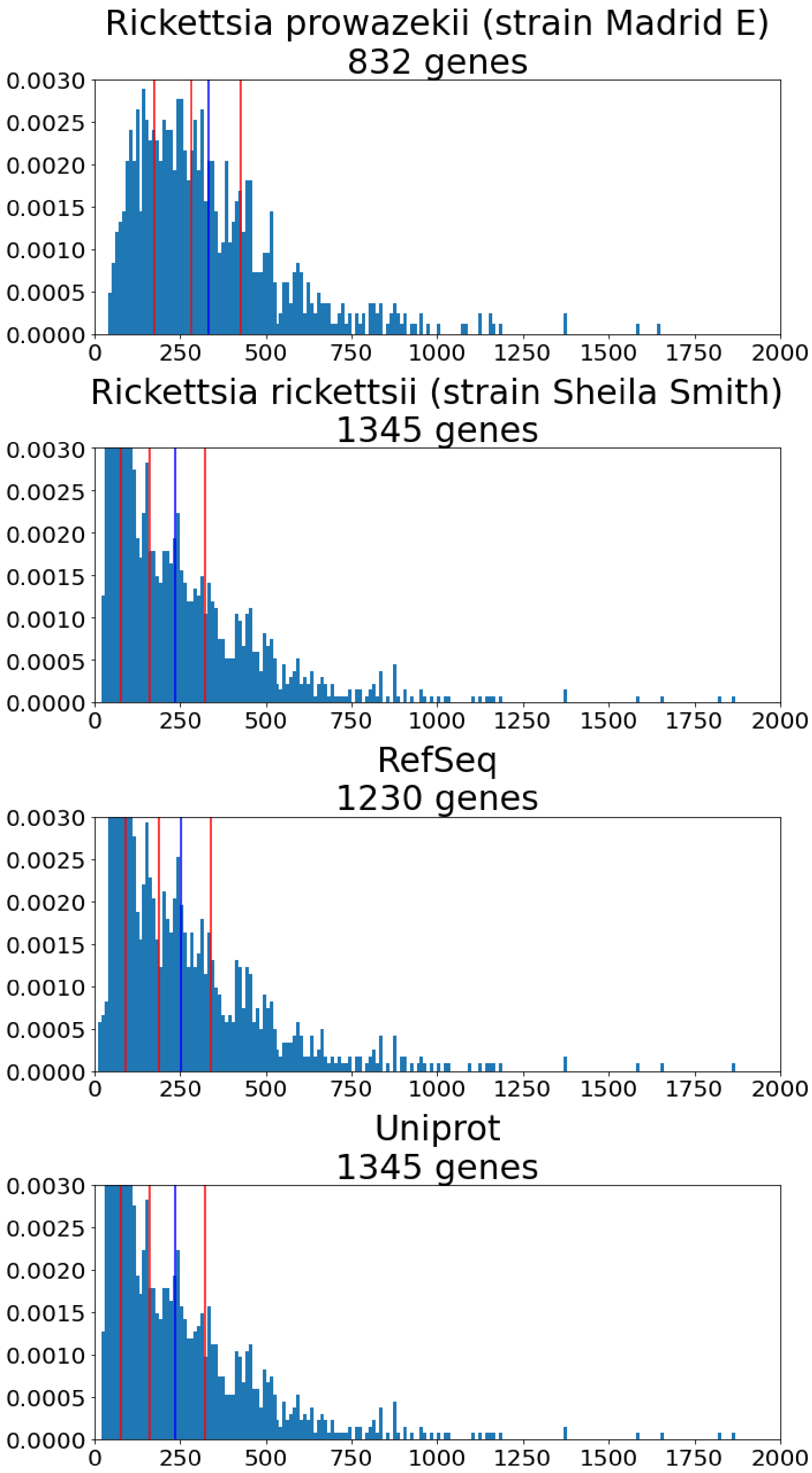
Rickettsia prowazekii (strain Madrid E)
832 genes
Rickettsia rickettsii (strain Sheila Smith)
1345 genes
RefSeq
1230 genes
Uniprot
1345 genes
0.0030
0.0025
0.0020
0.0015
0.0010
0.0005
0.0000
0
250
500
750
1000
1250
1500
1750
2000

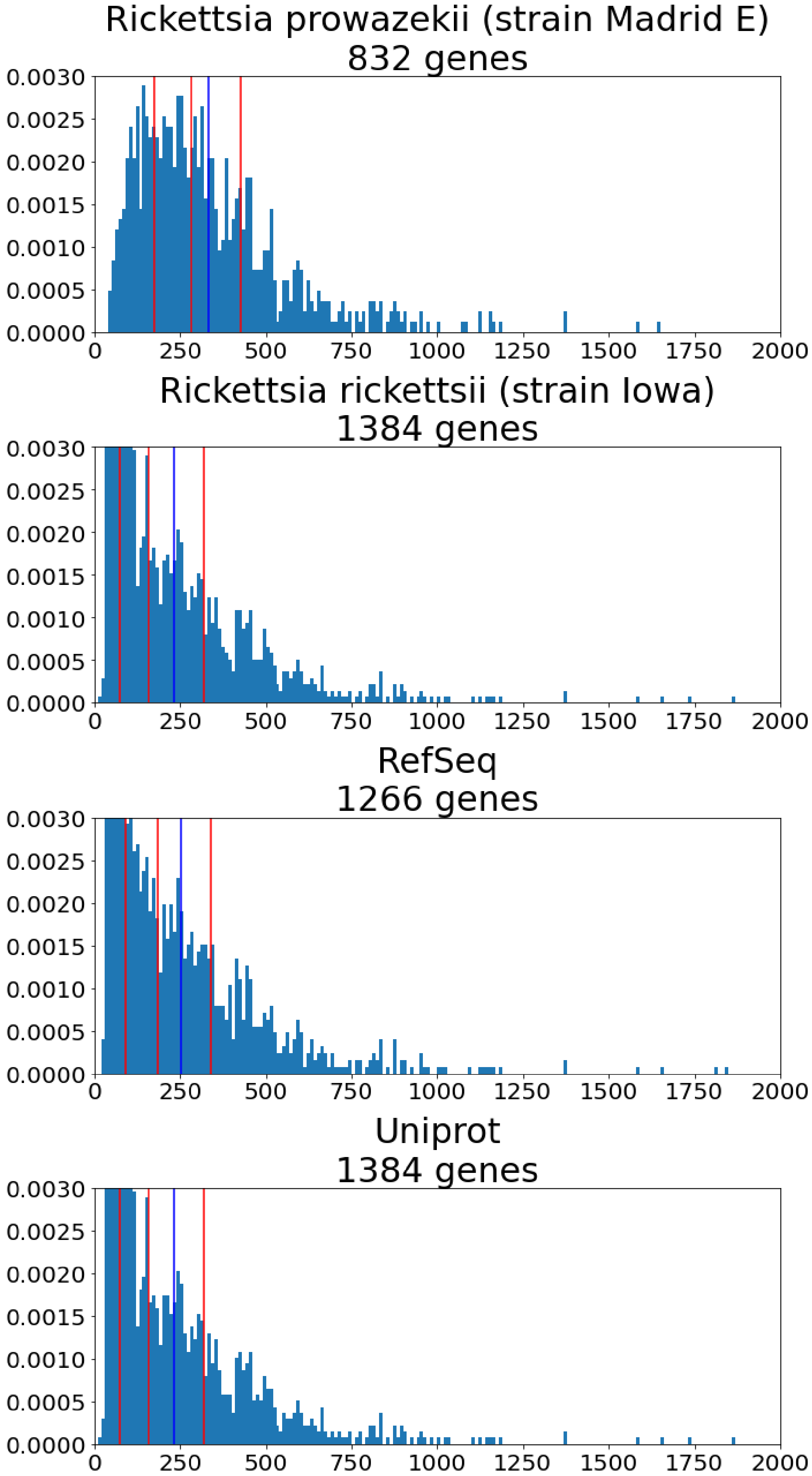

Rickettsia prowazekii (strain Madrid E)
832 genes
Rickettsia rickettsii (strain Iowa)
1384 genes
RefSeq
1266 genes
Uniprot
1384 genes
0.0030
0.0025
0.0020
0.0015
0.0010
0.0005
0.0000
0
250
500
750
1000
1250
1500
1750
2000

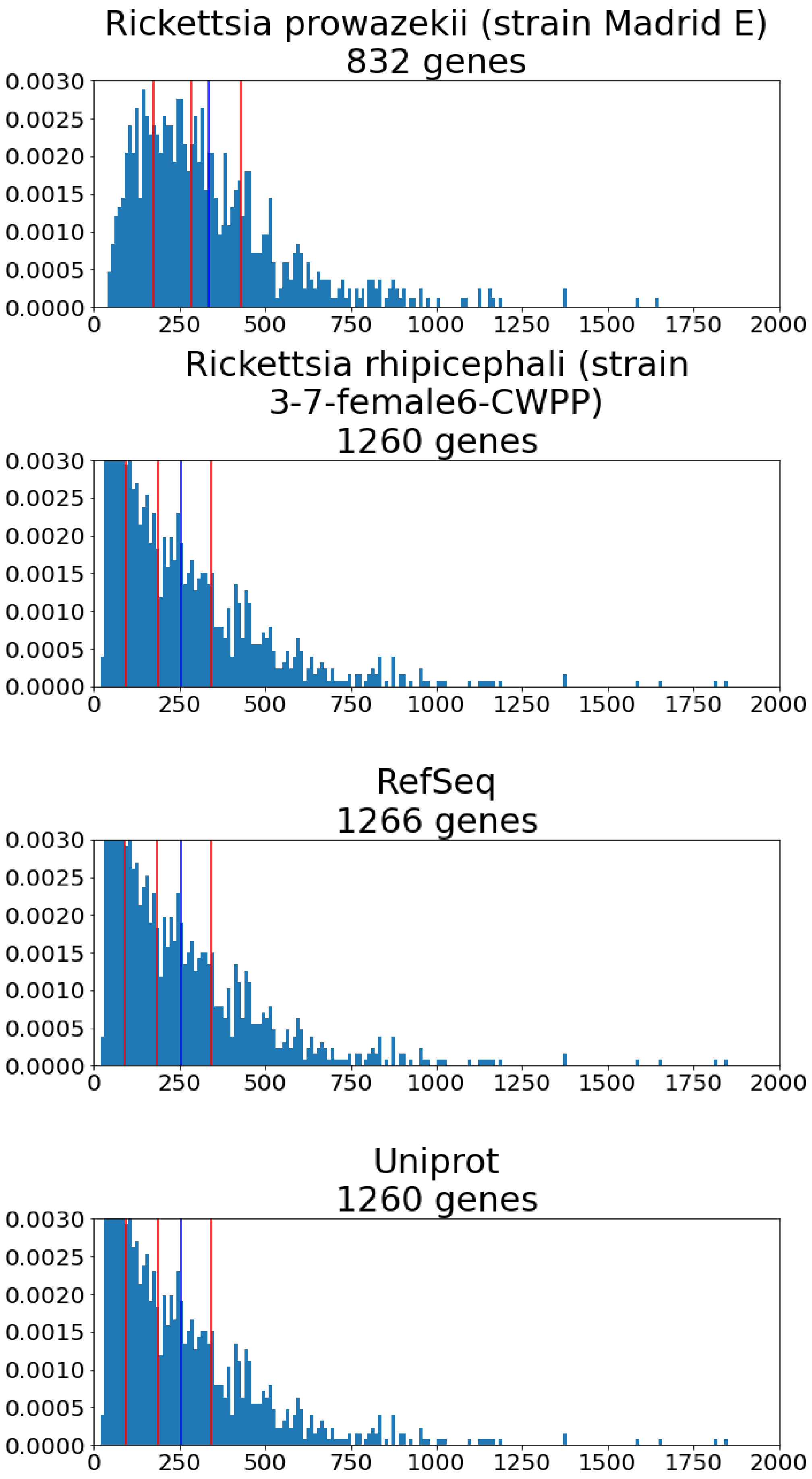
Rickettsia prowazekii (strain Madrid E)
832 genes
Rickettsia rhipicephali (strain
3-7-female6-CWPP)
1260 genes
RefSeq
1266 genes
Uniprot
1260 genes
0.0030
0.0025
0.0020
0.0015
0.0010
0.0005
0.0000
0
250
500
750
1000
1250
1500
1750
2000

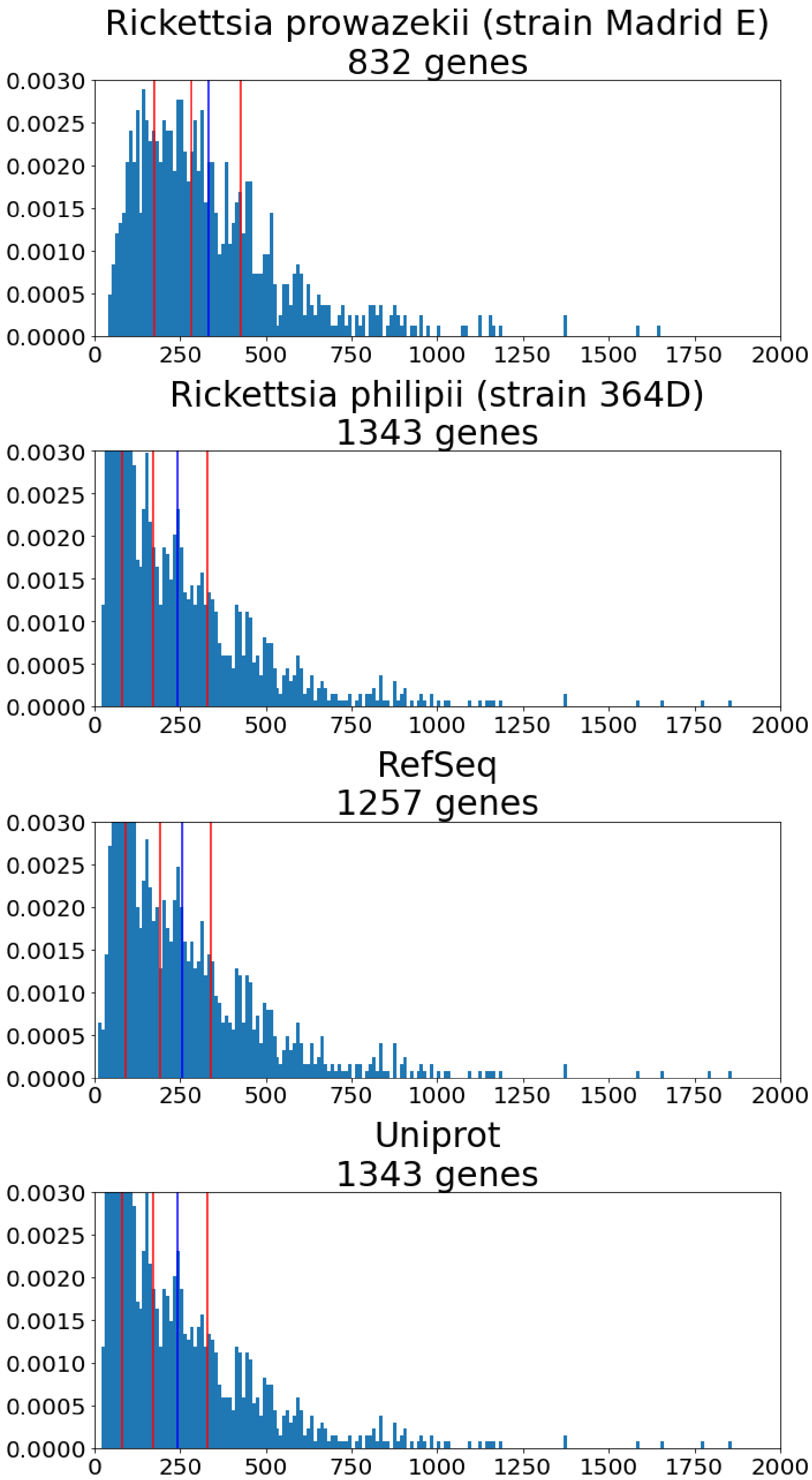
Rickettsia prowazekii (strain Madrid E)
832 genes
Rickettsia philipii (strain 364D)
1343 genes
RefSeq
1257 genes
Uniprot
1343 genes
0.0030
0.0025
0.0020
0.0015
0.0010
0.0005
0.0000
0
250
500
750
1000
1250
1500
1750
2000

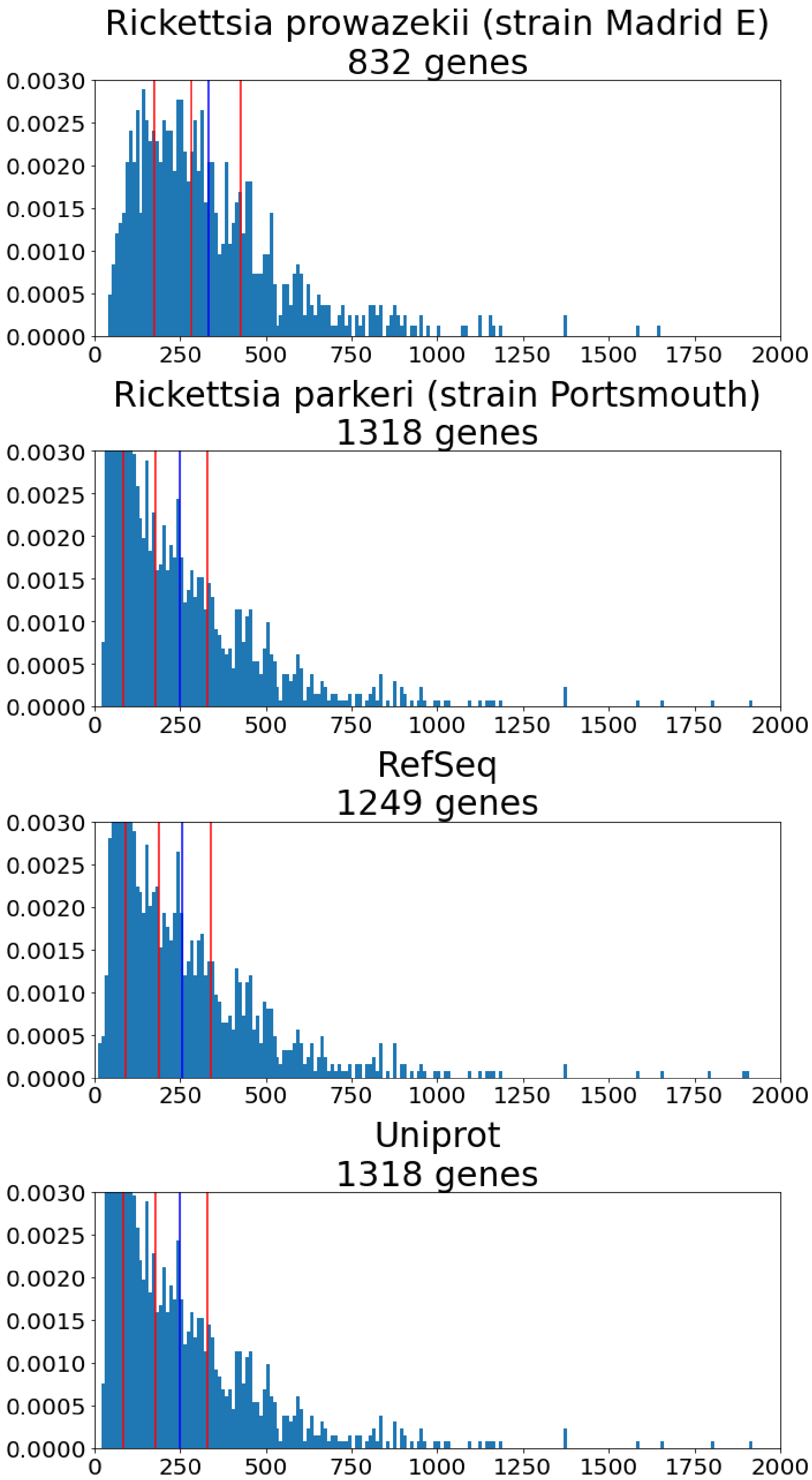
Rickettsia prowazekii (strain Madrid E)
832 genes
Rickettsia parkeri (strain Portsmouth)
1318 genes
RefSeq
1249 genes
Uniprot
1318 genes
0.0030
0.0025
0.0020
0.0015
0.0010
0.0005
0.0000
0
250
500
750
1000
1250
1500
1750
2000

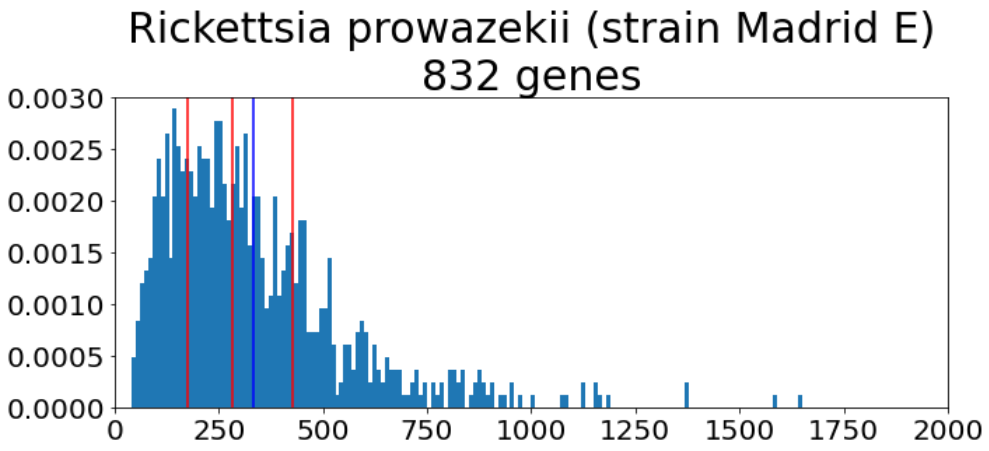
Rickettsia prowazekii (strain Madrid E)
832 genes

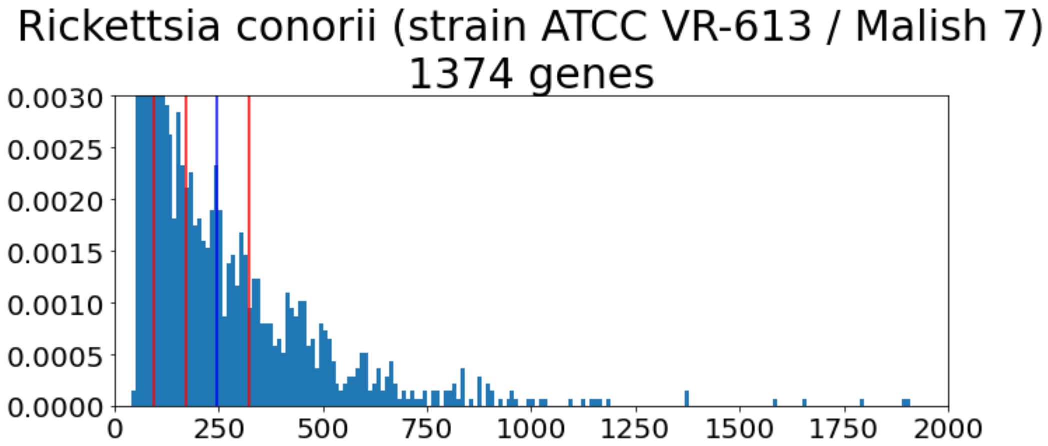
Rickettsia conorii (strain ATCC VR-613 / Malish 7)
1374 genes

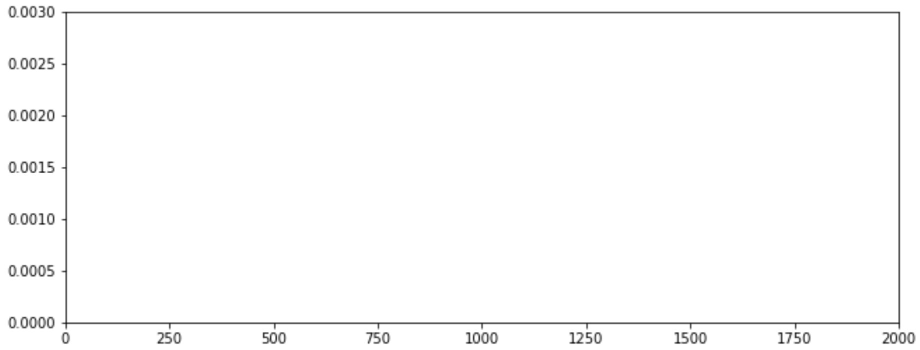

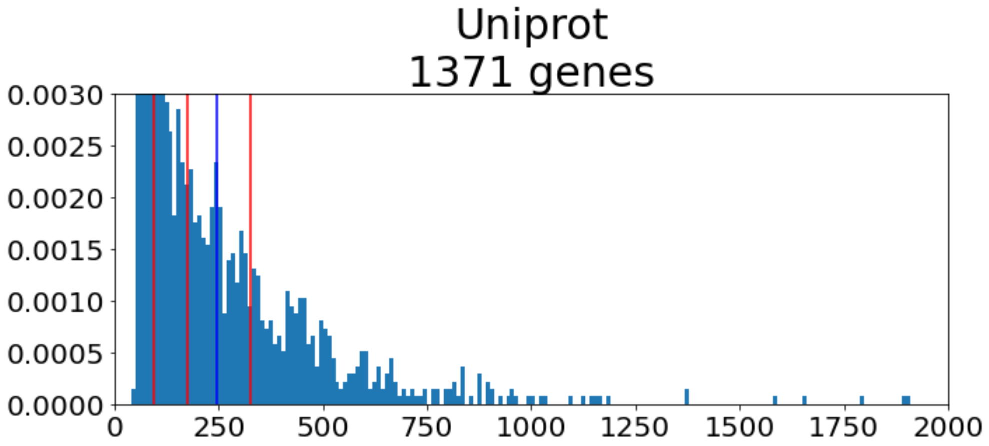
Uniprot
1371 genes

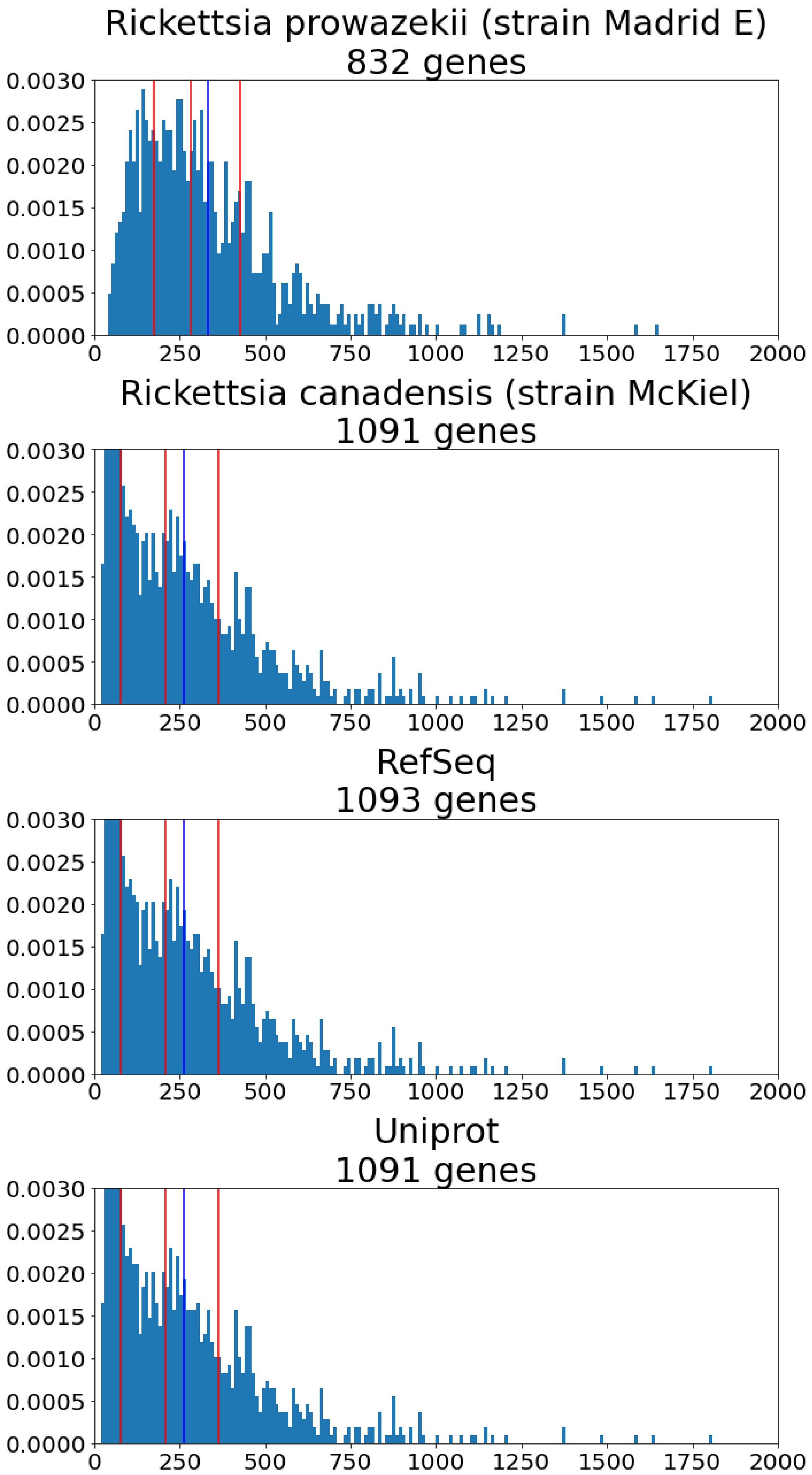
Rickettsia prowazekii (strain Madrid E)
832 genes
Rickettsia canadensis (strain McKiel)
1091 genes
RefSeq
1093 genes
Uniprot
1091 genes
0.0030
0.0025
0.0020
0.0015
0.0010
0.0005
0.0000
0
250
500
750
1000
1250
1500
1750
2000

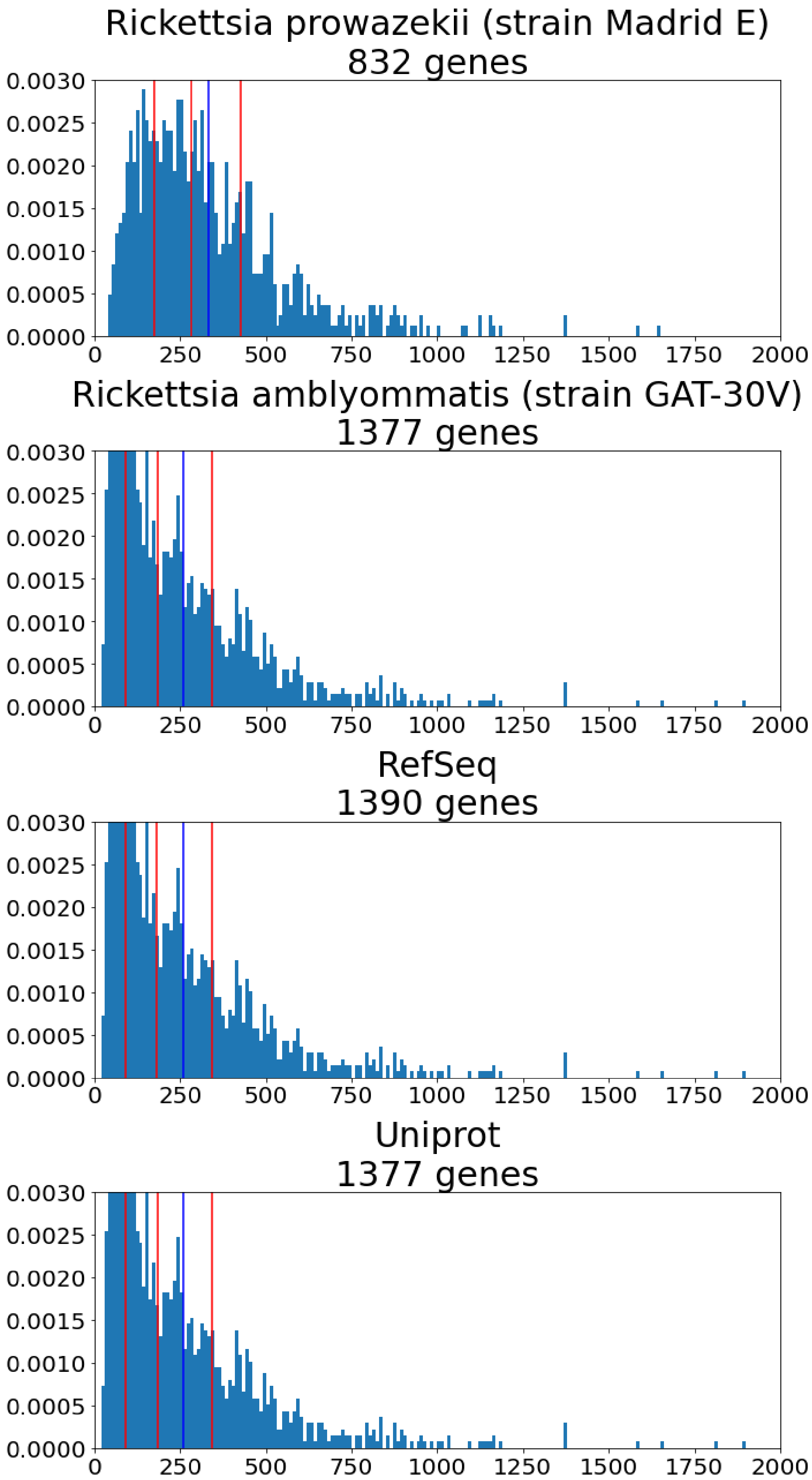
Rickettsia prowazekii (strain Madrid E)
832 genes
Rickettsia amblyommatis (strain GAT-30V)
1377 genes
RefSeq
1390 genes
Uniprot
1377 genes
0.0030
0.0025
0.0020
0.0015
0.0010
0.0005
0.0000
0
250
500
750
1000
1250
1500
1750
2000

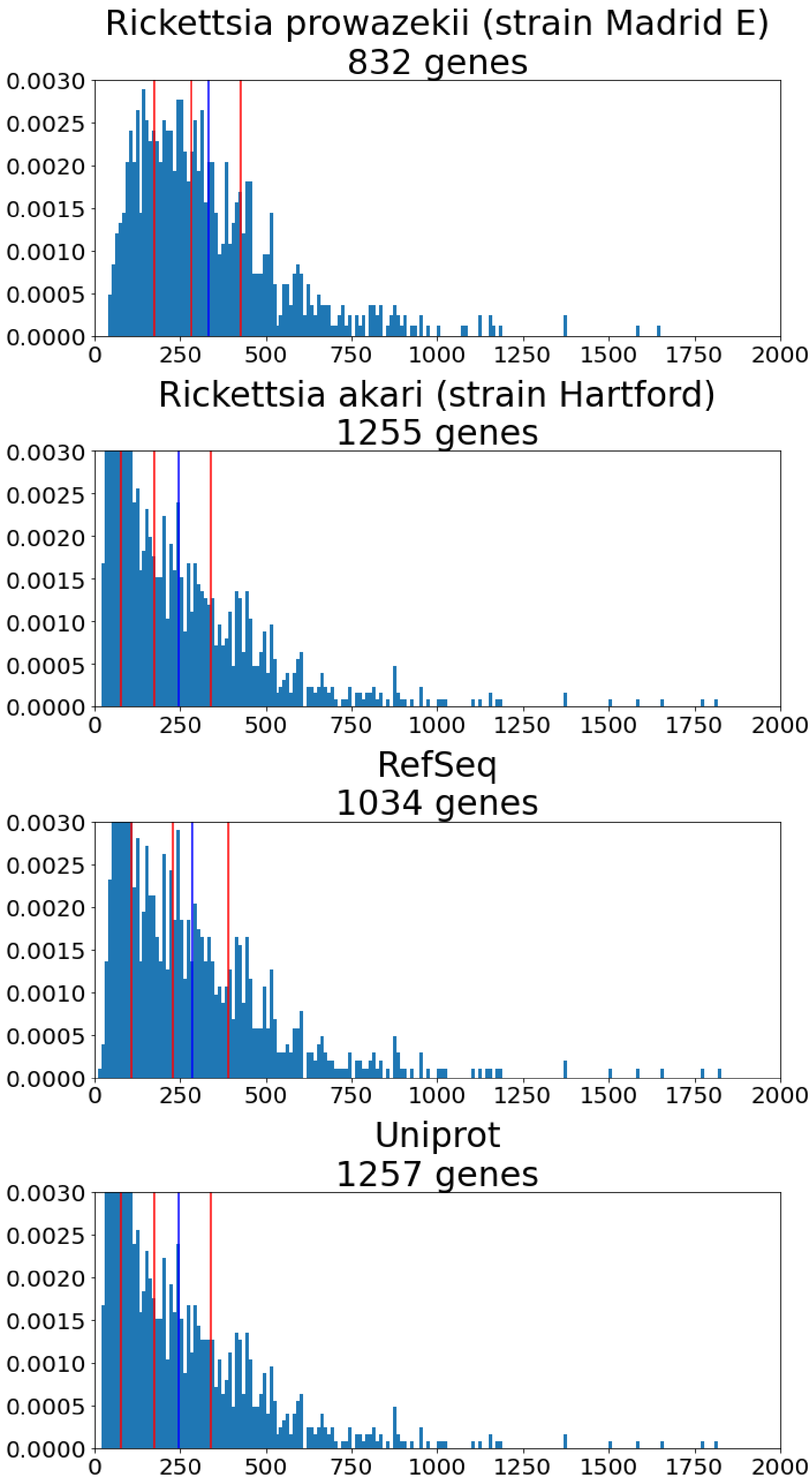

Rickettsia prowazekii (strain Madrid E)
832 genes
Rickettsia akari (strain Hartford)
1255 genes
RefSeq
1034 genes
Uniprot
1257 genes
0.0000
0.0005
0.0010
0.0015
0.0020
0.0025
0.0030
0
250
500
750
1000
1250
1500
1750
2000

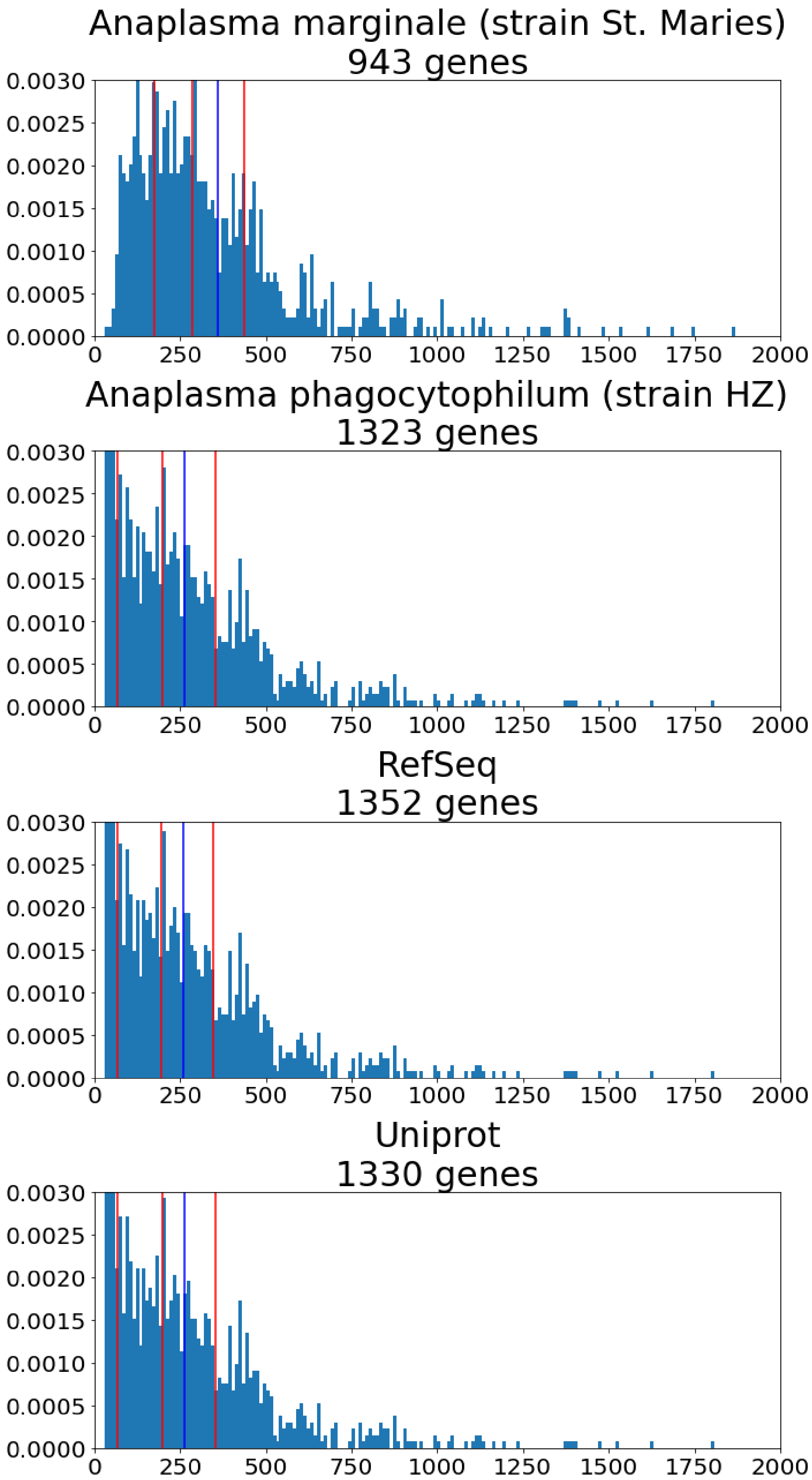
Anaplasma marginale (strain St. Maries)
943 genes
Anaplasma phagocytophilum (strain HZ)
1323 genes
RefSeq
1352 genes
Uniprot
1330 genes
0.0000
0.0005
0.0010
0.0015
0.0020
0.0025
0.0030
0
250
500
750
1000
1250
1500
1750
2000

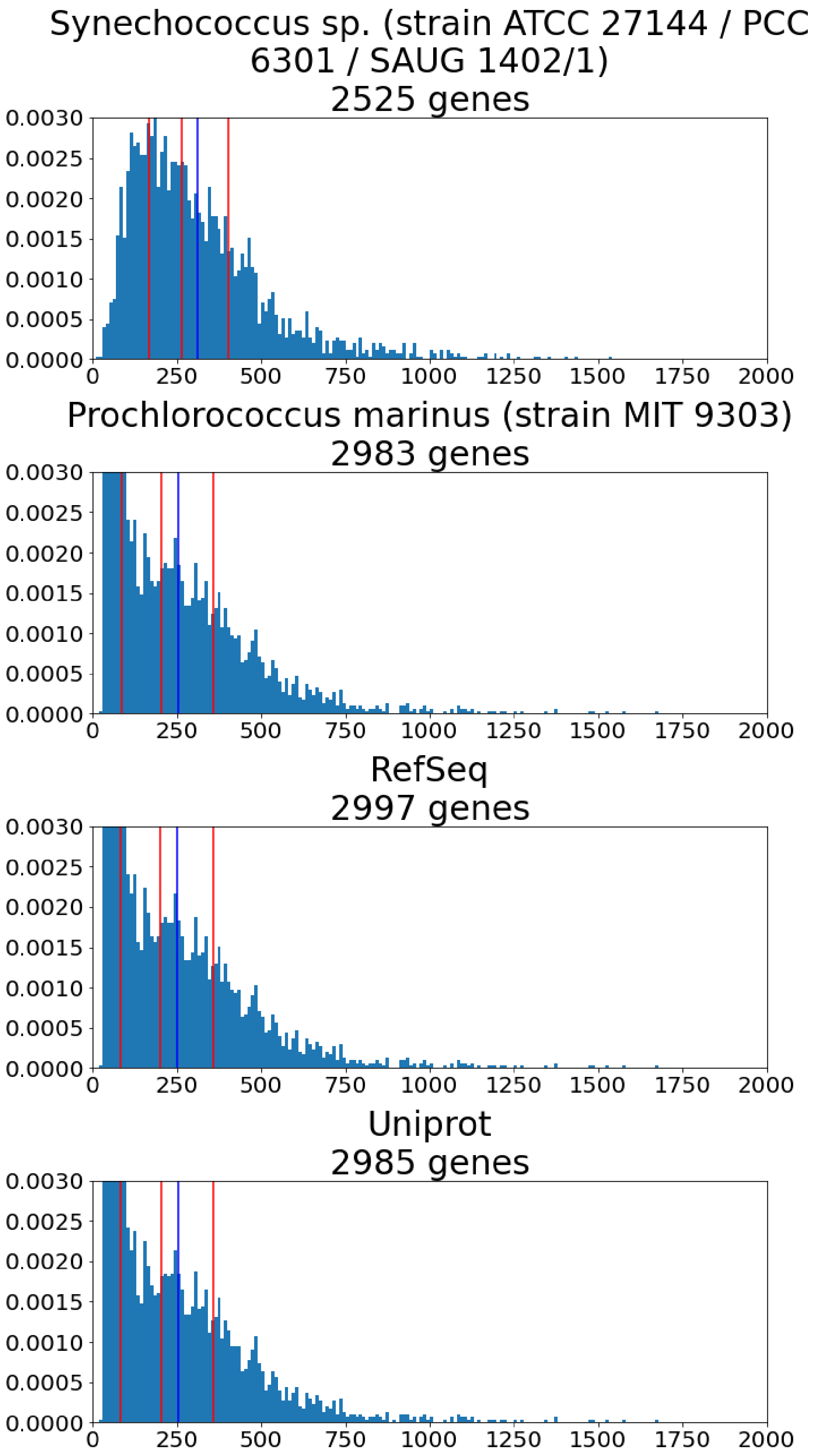
Synechococcus sp. (strain ATCC 27144 / PCC 6301 / SAUG 1402/1)
2525 genes
Prochlorococcus marinus (strain MIT 9303)
2983 genes
RefSeq
2997 genes
Uniprot
2985 genes
0.0000
0.0005
0.0010
0.0015
0.0020
0.0025
0.0030
0
250
500
750
1000
1250
1500
1750
2000

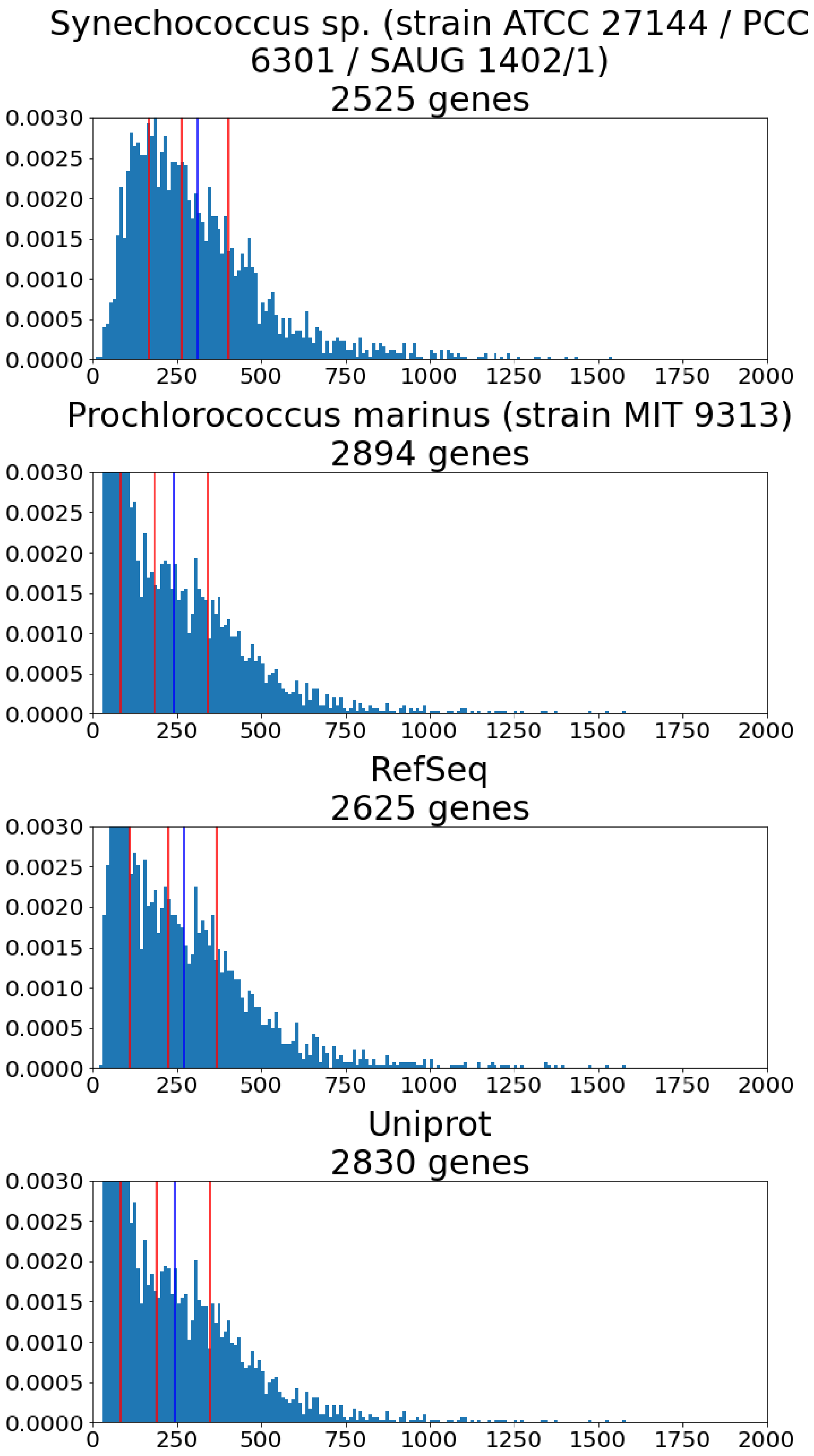

Synechococcus sp. (strain ATCC 27144 / PCC 6301 / SAUG 1402/1)
2525 genes
Prochlorococcus marinus (strain MIT 9313)
2894 genes
RefSeq
2625 genes
Uniprot
2830 genes
0.0000
0.0005
0.0010
0.0015
0.0020
0.0025
0.0030
0
250
500
750
1000
1250
1500
1750
2000

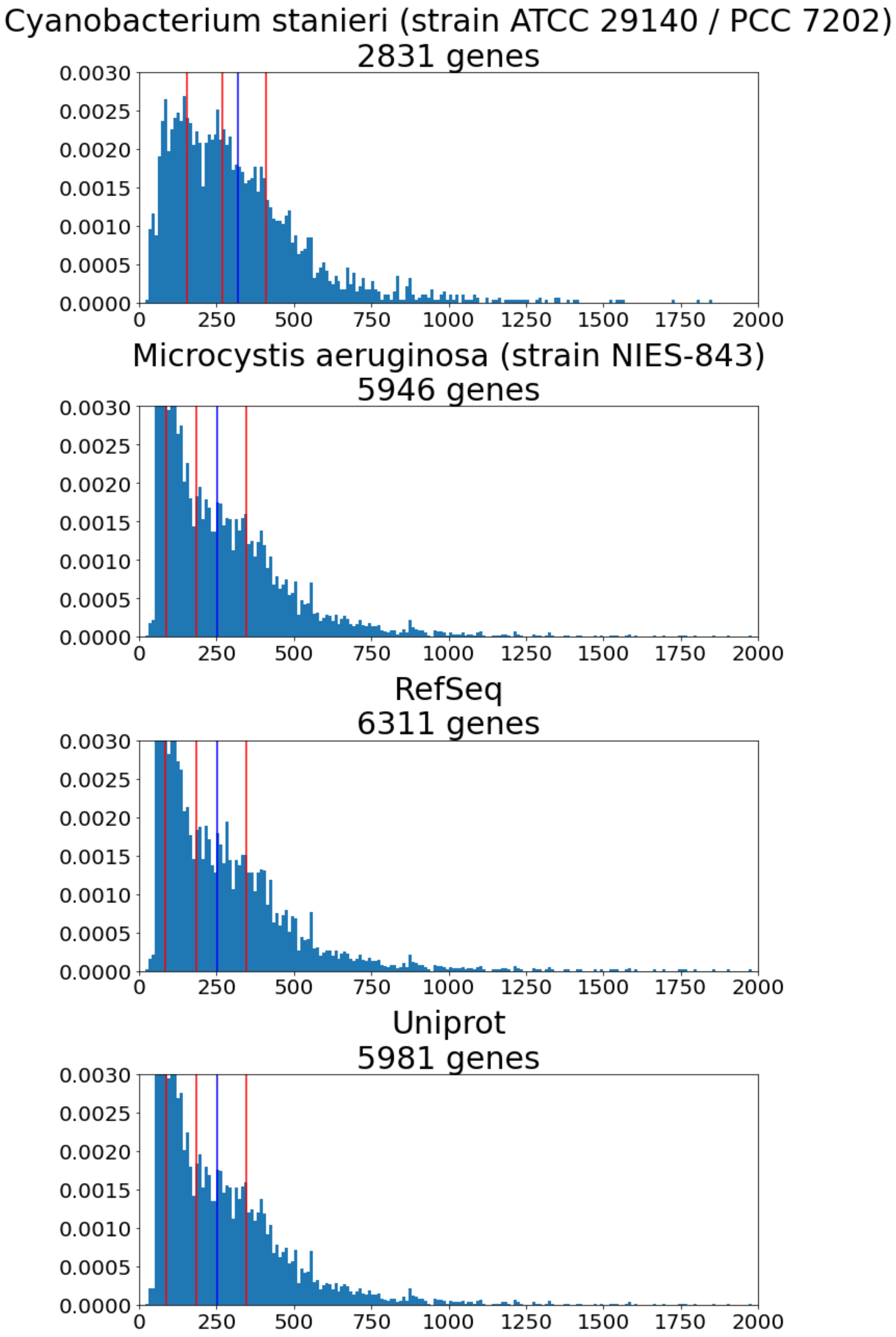
Cyanobacterium stanieri (strain ATCC 29140 / PCC 7202)
2831 genes
Microcystis aeruginosa (strain NIES-843)
5946 genes
RefSeq
6311 genes
Uniprot
5981 genes
0.0000
0.0005
0.0010
0.0015
0.0020
0.0025
0.0030
0
250
500
750
1000
1250
1500
1750
2000

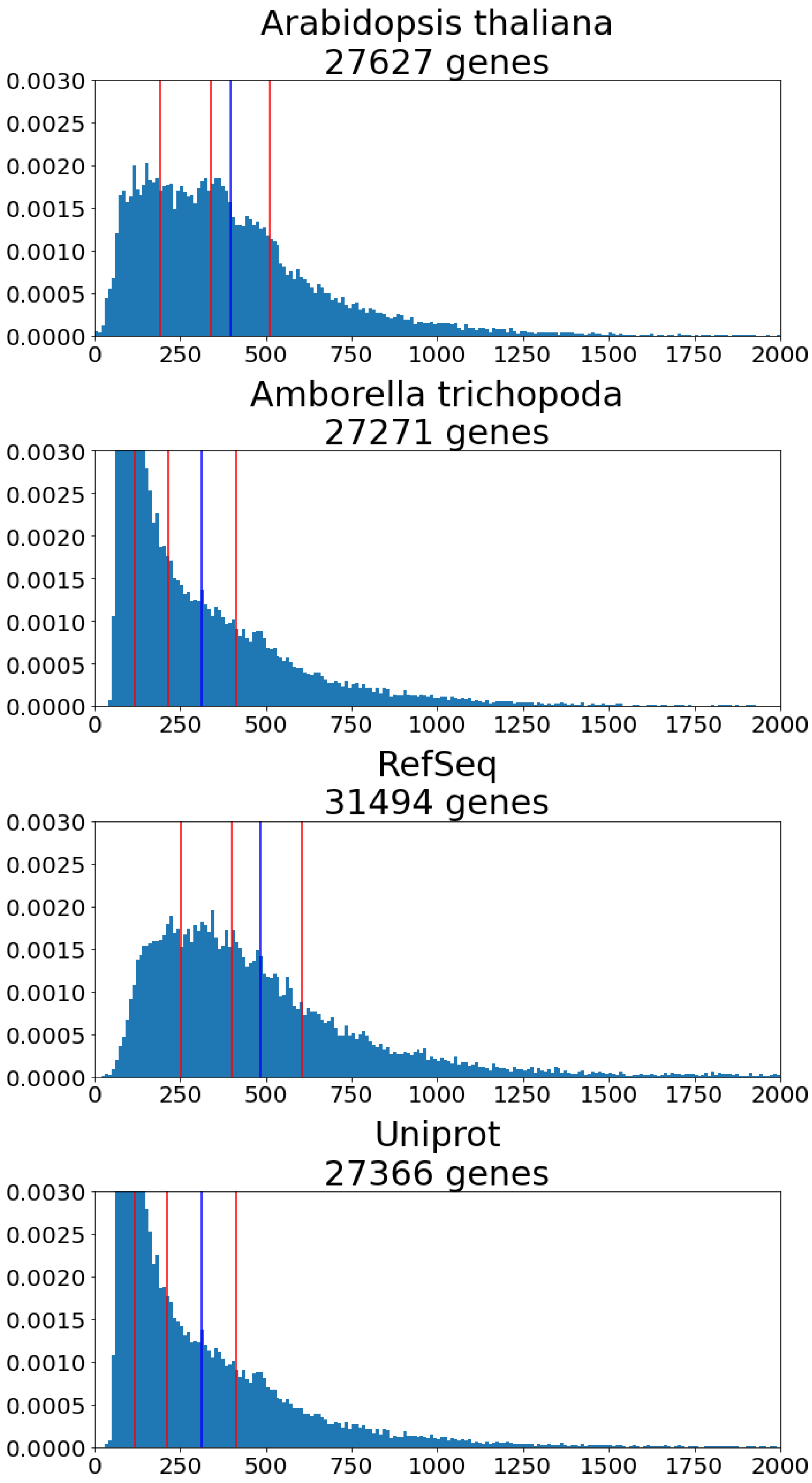
Arabidopsis thaliana
27627 genes
Amborella trichopoda
27271 genes
RefSeq
31494 genes
Uniprot
27366 genes
0.0030
0.0025
0.0020
0.0015
0.0010
0.0005
0.0000
0
250
500
750
1000
1250
1500
1750
2000

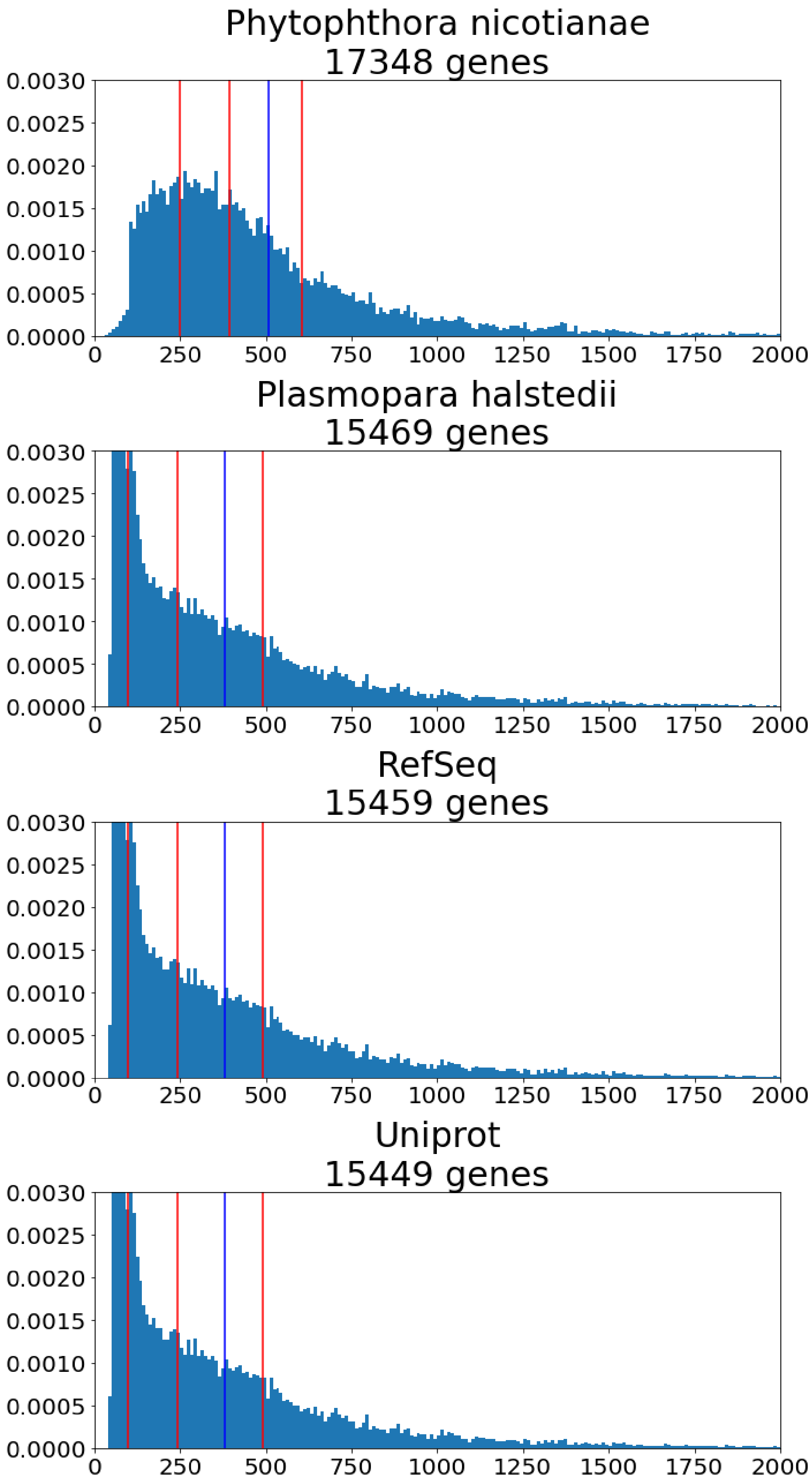
Phytophthora nicotianae
17348 genes
Plasmopara halstedii
15469 genes
RefSeq
15459 genes
Uniprot
15449 genes
0.0030
0.0025
0.0020
0.0015
0.0010
0.0005
0.0000
0
250
500
750
1000
1250
1500
1750
2000

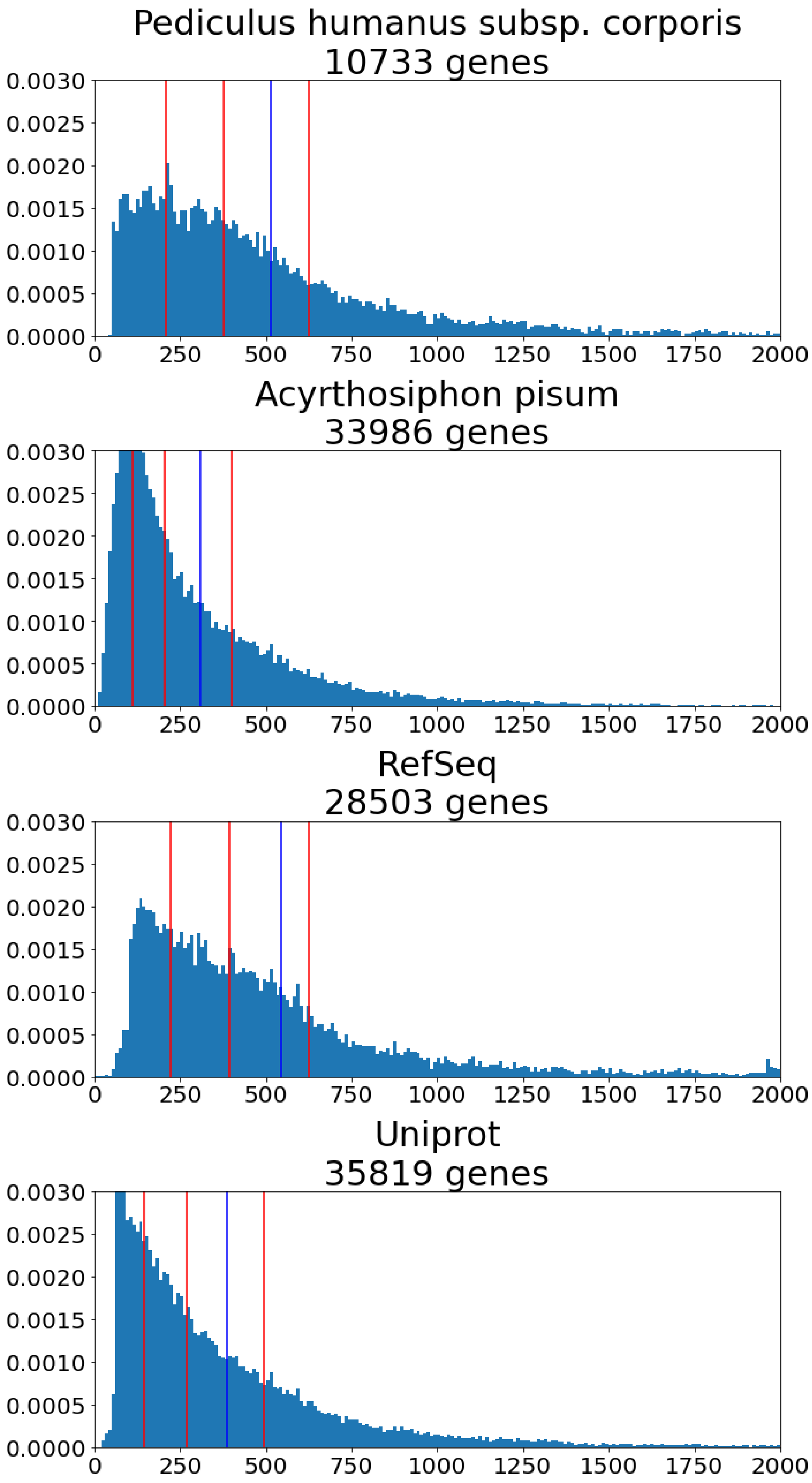
Pediculus humanus subsp. corporis
10733 genes
Acyrthosiphon pisum
33986 genes
RefSeq
28503 genes
Uniprot
35819 genes
0.0030
0.0025
0.0020
0.0015
0.0010
0.0005
0.0000
0
250
500
750
1000
1250
1500
1750
2000

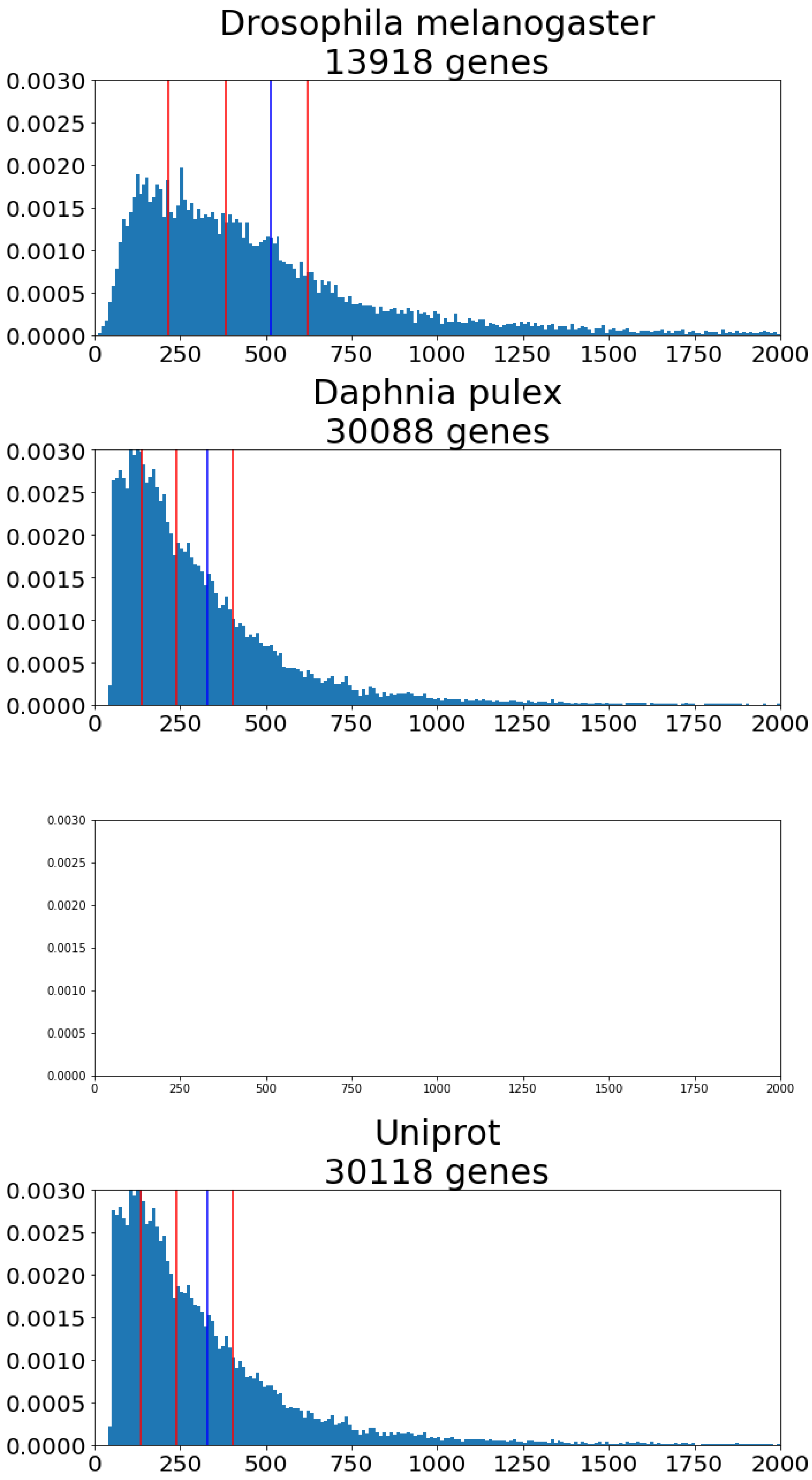
Drosophila melanogaster
13918 genes
0.0030
0.0025
0.0020
0.0015
0.0010
0.0005
0.0000
0 250 500 750 1000 1250 1500 1750 2000
Daphnia pulex
30088 genes
0.0030
0.0025
0.0020
0.0015
0.0010
0.0005
0.0000
0 250 500 750 1000 1250 1500 1750 2000
0.0030
0.0025
0.0020
0.0015
0.0010
0.0005
0.0000
0 250 500 750 1000 1250 1500 1750 2000
Uniprot
30118 genes
0.0030
0.0025
0.0020
0.0015
0.0010
0.0005
0.0000
0 250 500 750 1000 1250 1500 1750 2000

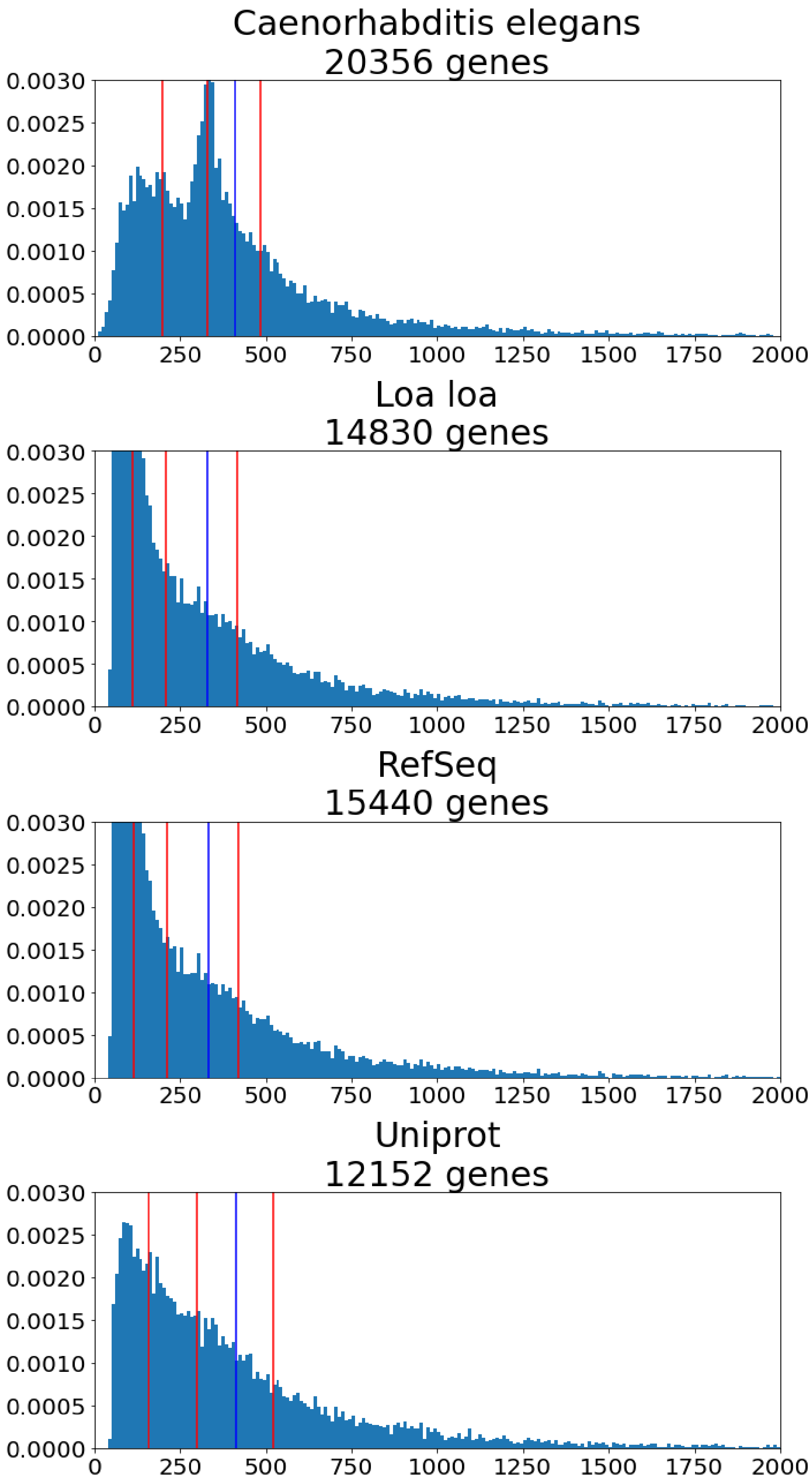
Caenorhabditis elegans
20356 genes
Loa loa
14830 genes
RefSeq
15440 genes
Uniprot
12152 genes
0.0030
0.0025
0.0020
0.0015
0.0010
0.0005
0.0000
0
250
500
750
1000
1250
1500
1750
2000

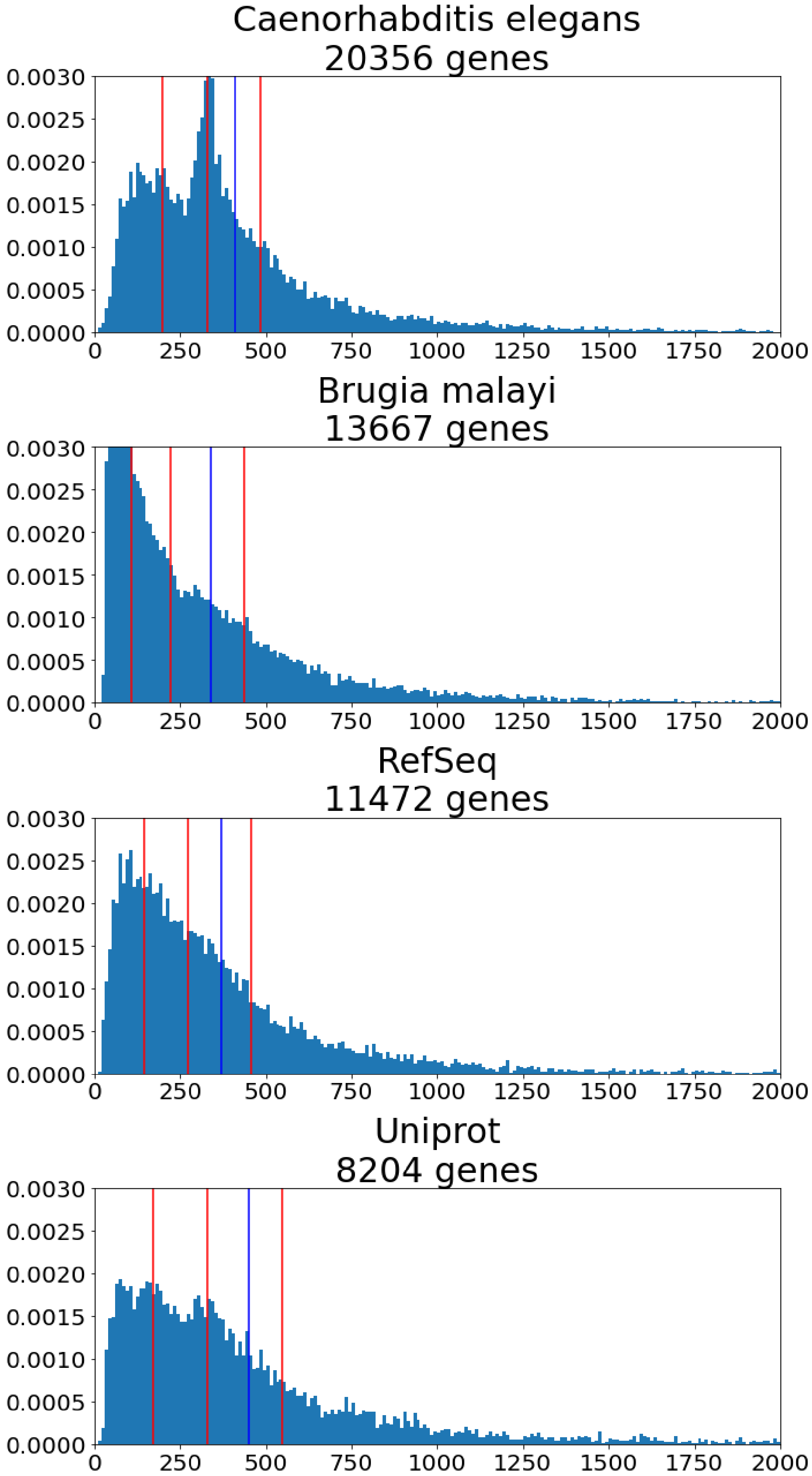
Caenorhabditis elegans
20356 genes
0.0030
0.0025
0.0020
0.0015
0.0010
0.0005
0.0000
0
250
500
750
1000
1250
1500
1750
2000
Brugia malayi
13667 genes
0.0030
0.0025
0.0020
0.0015
0.0010
0.0005
0.0000
0
250
500
750
1000
1250
1500
1750
2000
RefSeq
11472 genes
0.0030
0.0025
0.0020
0.0015
0.0010
0.0005
0.0000
0
250
500
750
1000
1250
1500
1750
2000
Uniprot
8204 genes
0.0030
0.0025
0.0020
0.0015
0.0010
0.0005
0.0000
0
250
500
750
1000
1250
1500
1750
2000

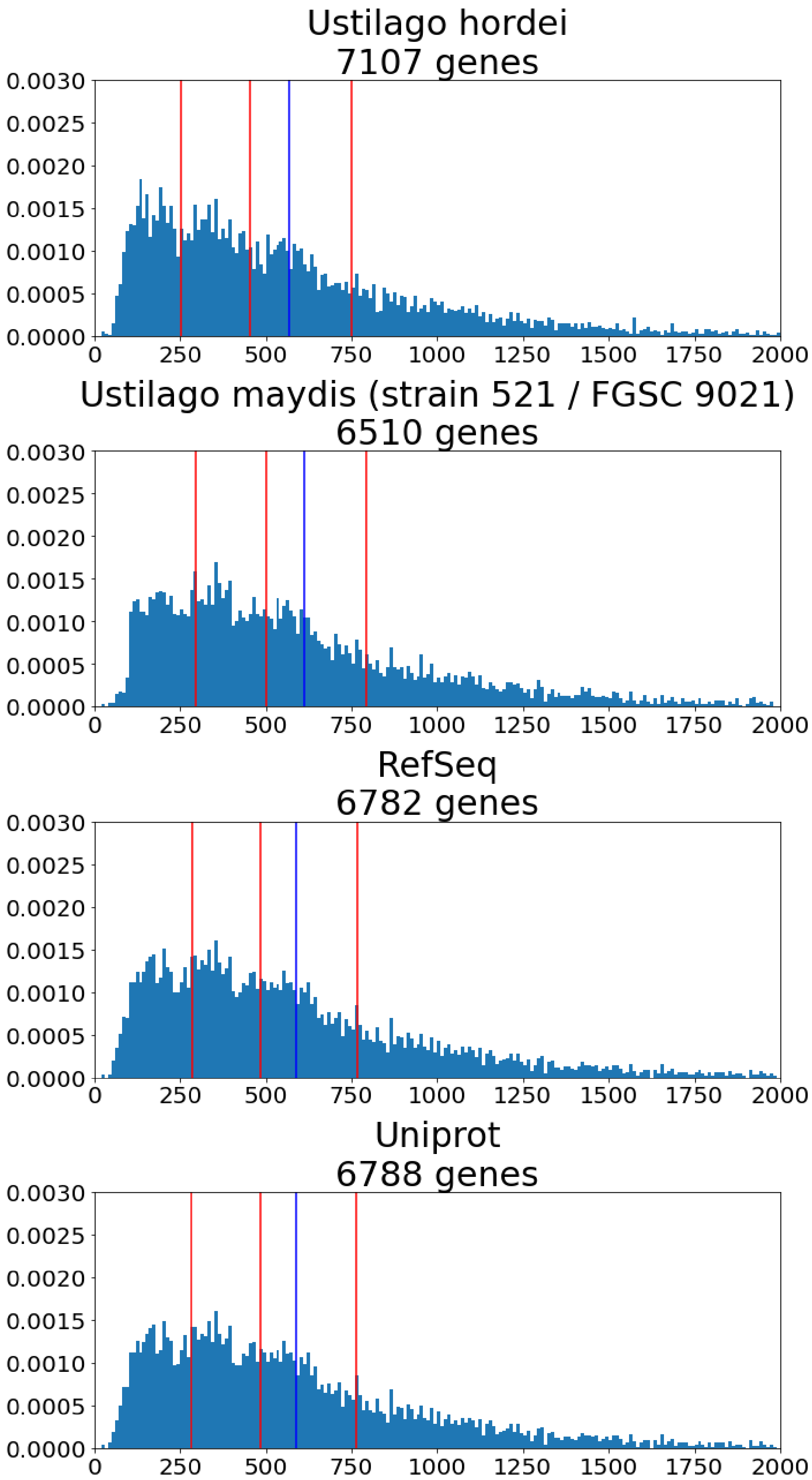
Ustilago hordei
7107 genes
Ustilago maydis (strain 521 / FGSC 9021)
6510 genes
RefSeq
6782 genes
Uniprot
6788 genes
0.0030
0.0025
0.0020
0.0015
0.0010
0.0005
0.0000
0
250
500
750
1000
1250
1500
1750
2000

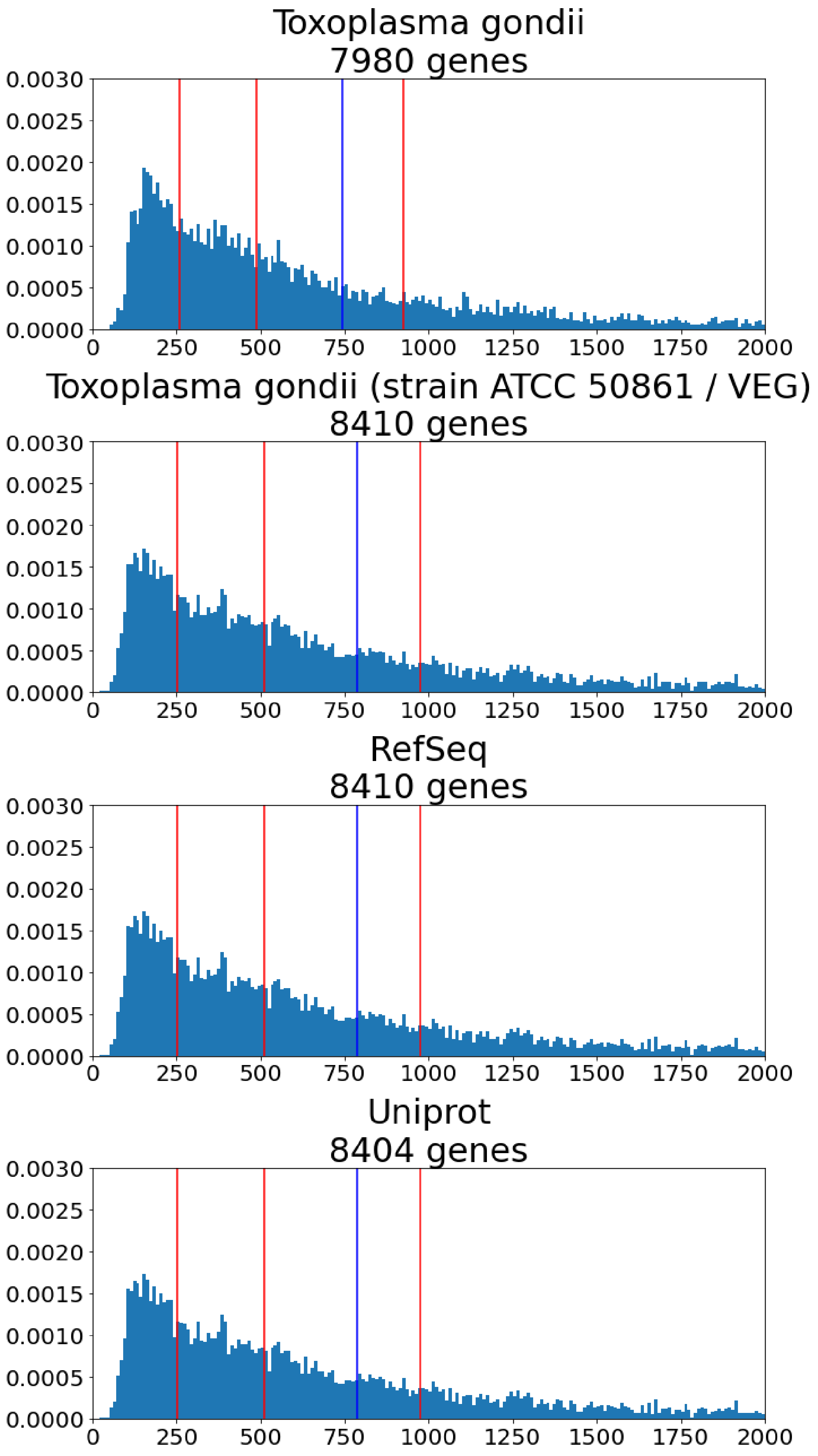
Toxoplasma gondii
7980 genes
Toxoplasma gondii (strain ATCC 50861 / VEG)
8410 genes
RefSeq
8410 genes
Uniprot
8404 genes
0.0030
0.0025
0.0020
0.0015
0.0010
0.0005
0.0000
0
250
500
750
1000
1250
1500
1750
2000

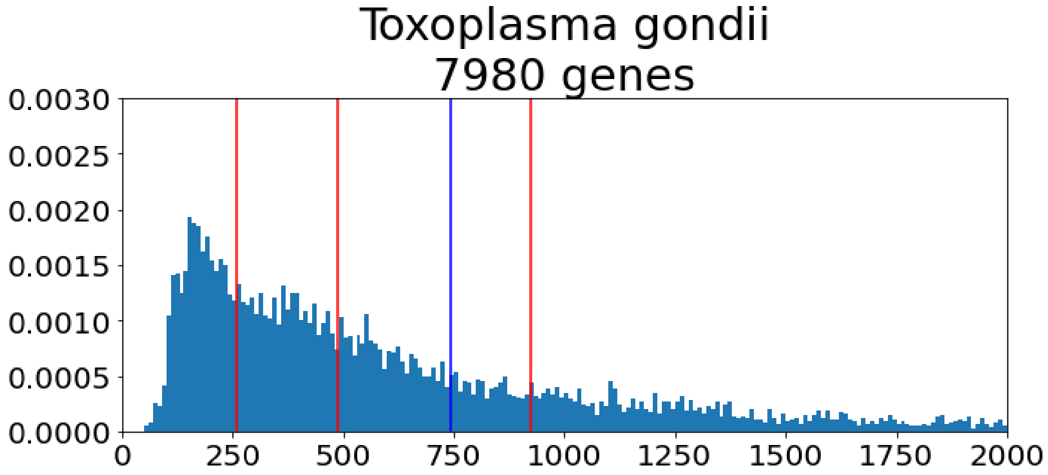
Toxoplasma gondii
7980 genes
0.0030
0.0025
0.0020
0.0015
0.0010
0.0005
0.0000
0
250
500
750
1000
1250
1500
1750
2000

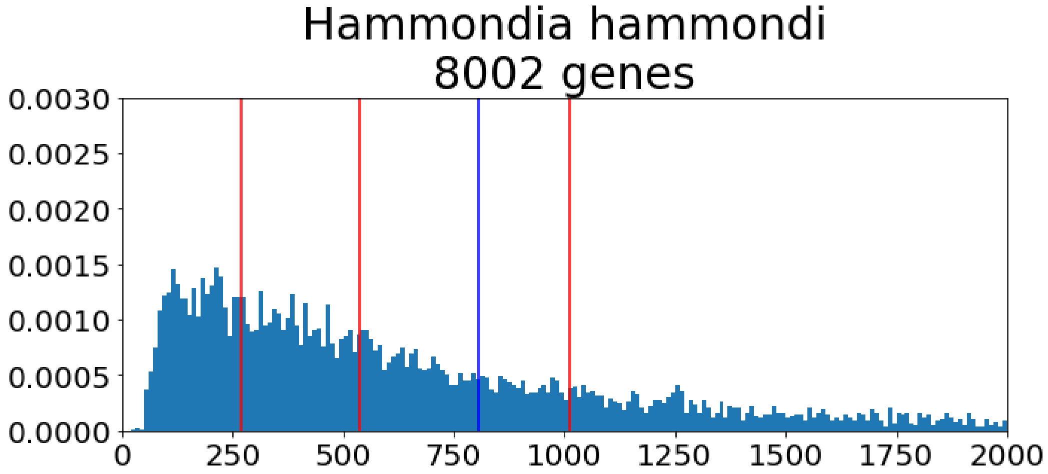
Hammondia hammondi
8002 genes
0.0030
0.0025
0.0020
0.0015
0.0010
0.0005
0.0000
0
250
500
750
1000
1250
1500
1750
2000

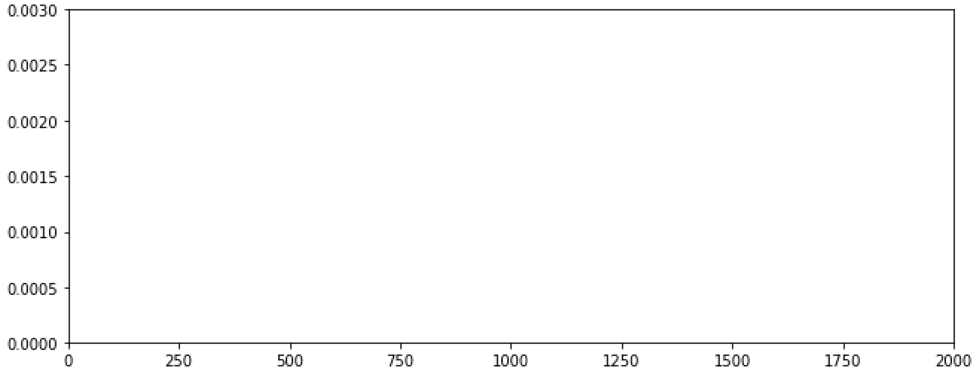
0.0030
0.0025
0.0020
0.0015
0.0010
0.0005
0.0000
0
250
500
750
1000
1250
1500
1750
2000

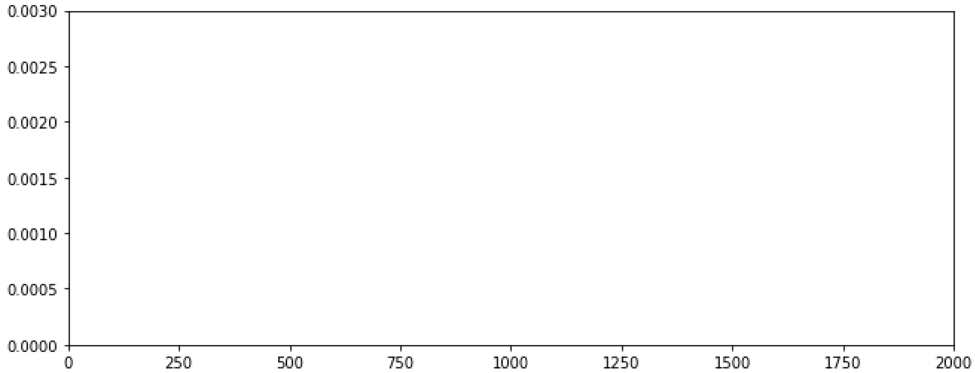
0.0030
0.0025
0.0020
0.0015
0.0010
0.0005
0.0000
0
250
500
750
1000
1250
1500
1750
2000

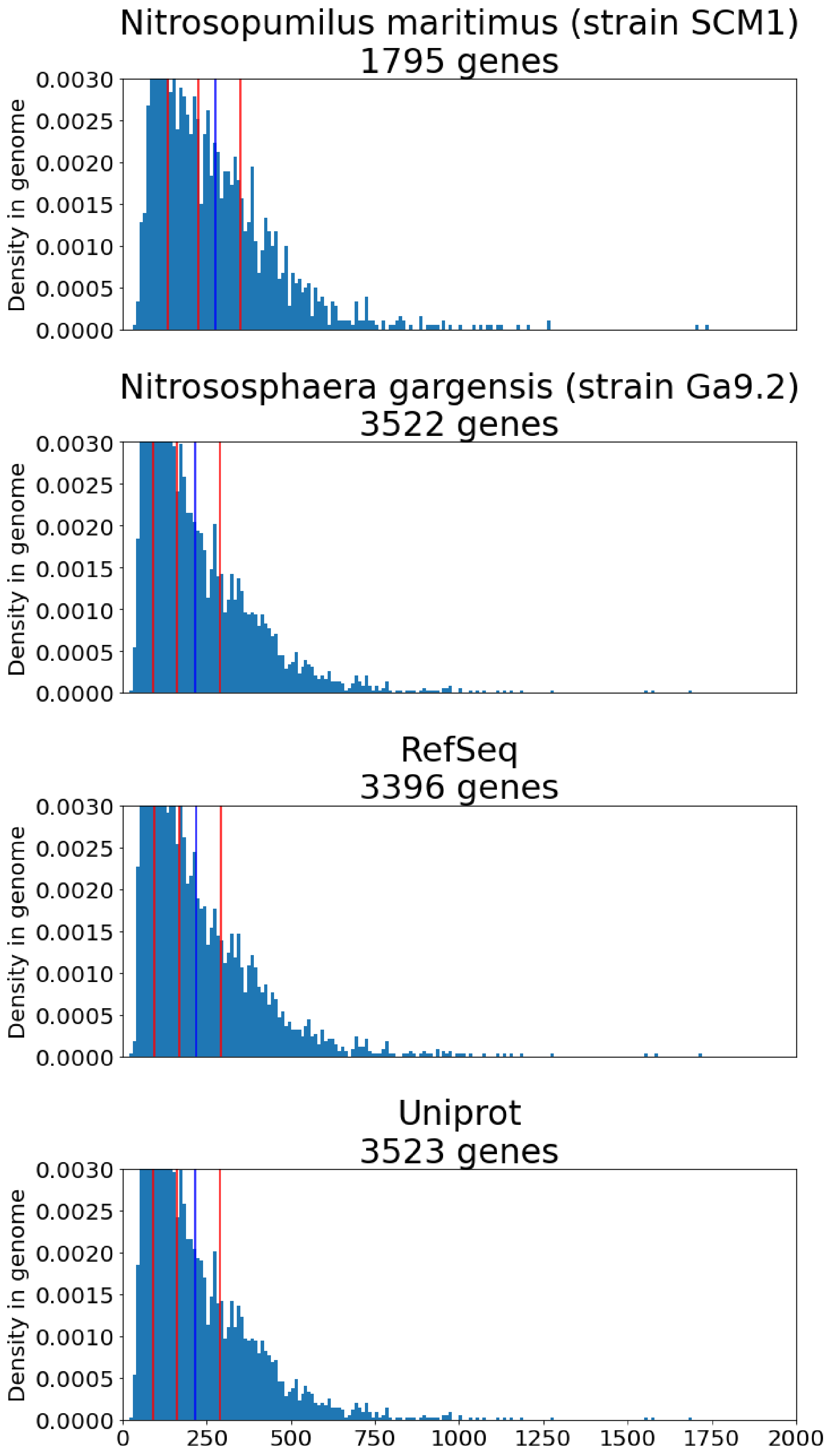

Nitrosopumilus maritimus (strain SCM1)
1795 genes
Nitrososphaera gargensis (strain Ga9.2)
3522 genes
RefSeq
3396 genes
Uniprot
3523 genes
Density in genome
0.0000
0.0005
0.0010
0.0015
0.0020
0.0025
0.0030
0
250
500
750
1000
1250
1500
1750
2000

Supplementary Figure 35: **Length distribution in the Rickettsia genus.** Left column represents length distribution in OMA (this study's dataset), the center column, the length distribution in RefSeq, and the right column the length distribution in Uniprot. The two rightmost columns are only filled for proteomes for which the distribution in OMA was labeled as outlier.

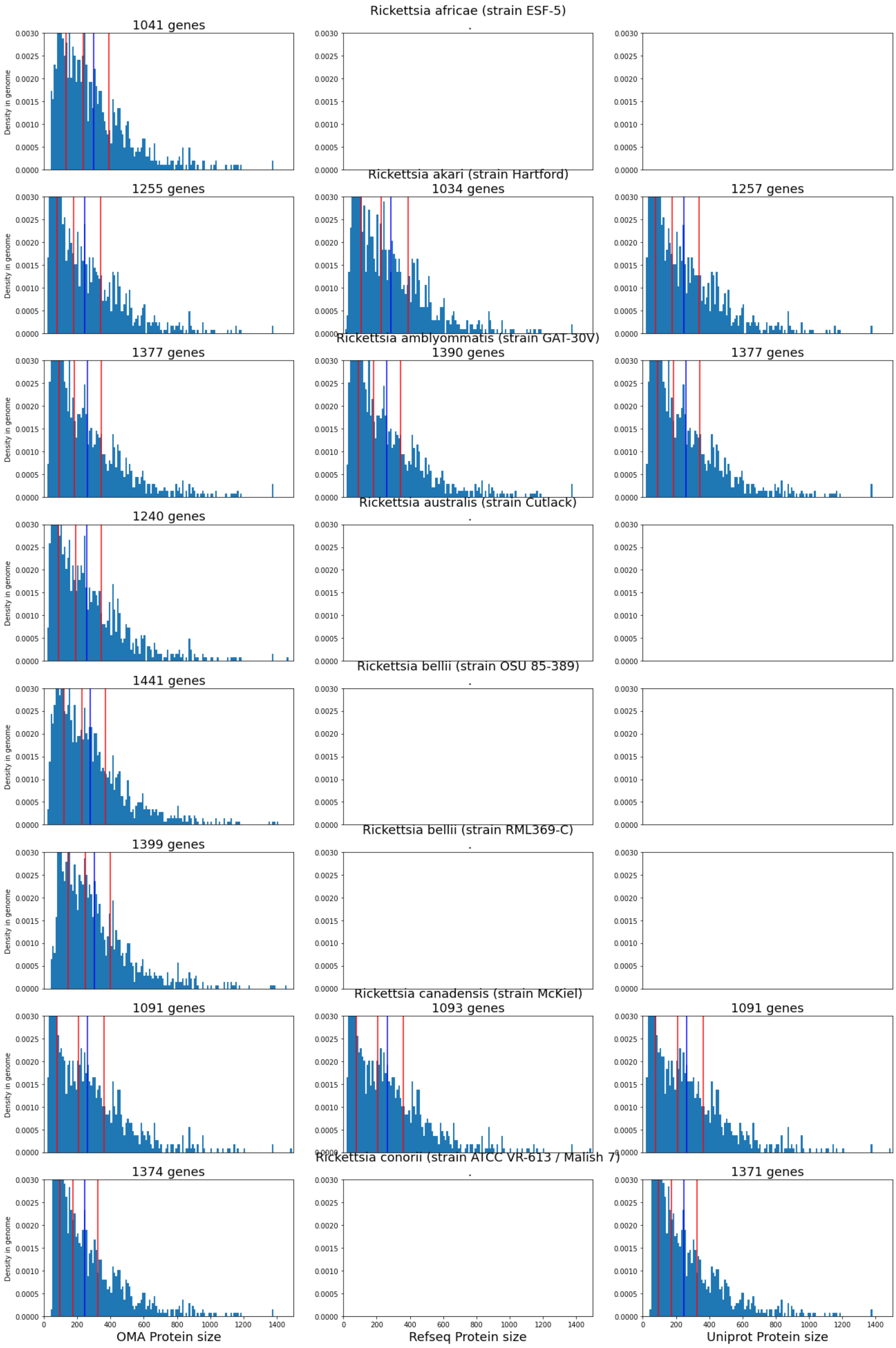
Rickettsia africae (strain ESF-5)
1041 genes
Rickettsia akari (strain Hartford)
1255 genes
1034 genes
1257 genes
Rickettsia amblyommatis (strain GAT-30V)
1377 genes
1390 genes
1377 genes
Rickettsia australis (strain Cutlack)
1240 genes
Rickettsia bellii (strain OSU 85-389)
1441 genes
Rickettsia bellii (strain RML369-C)
1399 genes
Rickettsia canadensis (strain McKiel)
1091 genes
1093 genes
1091 genes
Rickettsia conorii (strain ATCC VR-613 / Malish 7)
1374 genes
1371 genes
Density in genome
OMA Protein size
Refseq Protein size
Uniprot Protein size

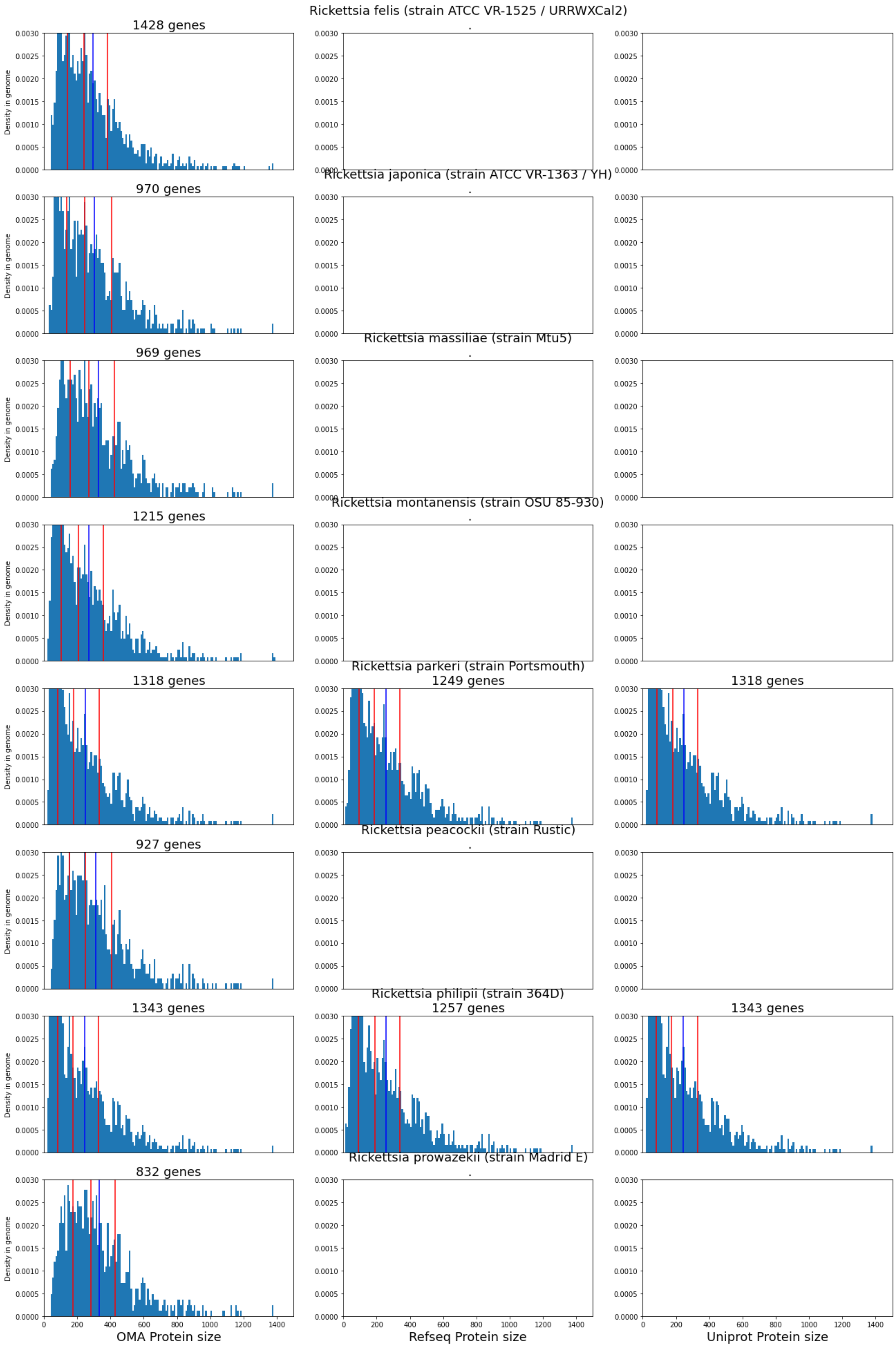

Rickettsia felis (strain ATCC VR-1525 / URRWXCal2)
1428 genes
Rickettsia japonica (strain ATCC VR-1363 / YH)
970 genes
Rickettsia massiliae (strain Mtu5)
969 genes
Rickettsia montanensis (strain OSU 85-930)
1215 genes
Rickettsia parkeri (strain Portsmouth)
1318 genes
1249 genes
1318 genes
Rickettsia peacockii (strain Rustic)
927 genes
Rickettsia philipii (strain 364D)
1343 genes
1257 genes
1343 genes
Rickettsia prowazekii (strain Madrid E)
832 genes
Density in genome
OMA Protein size
Refseq Protein size
Uniprot Protein size

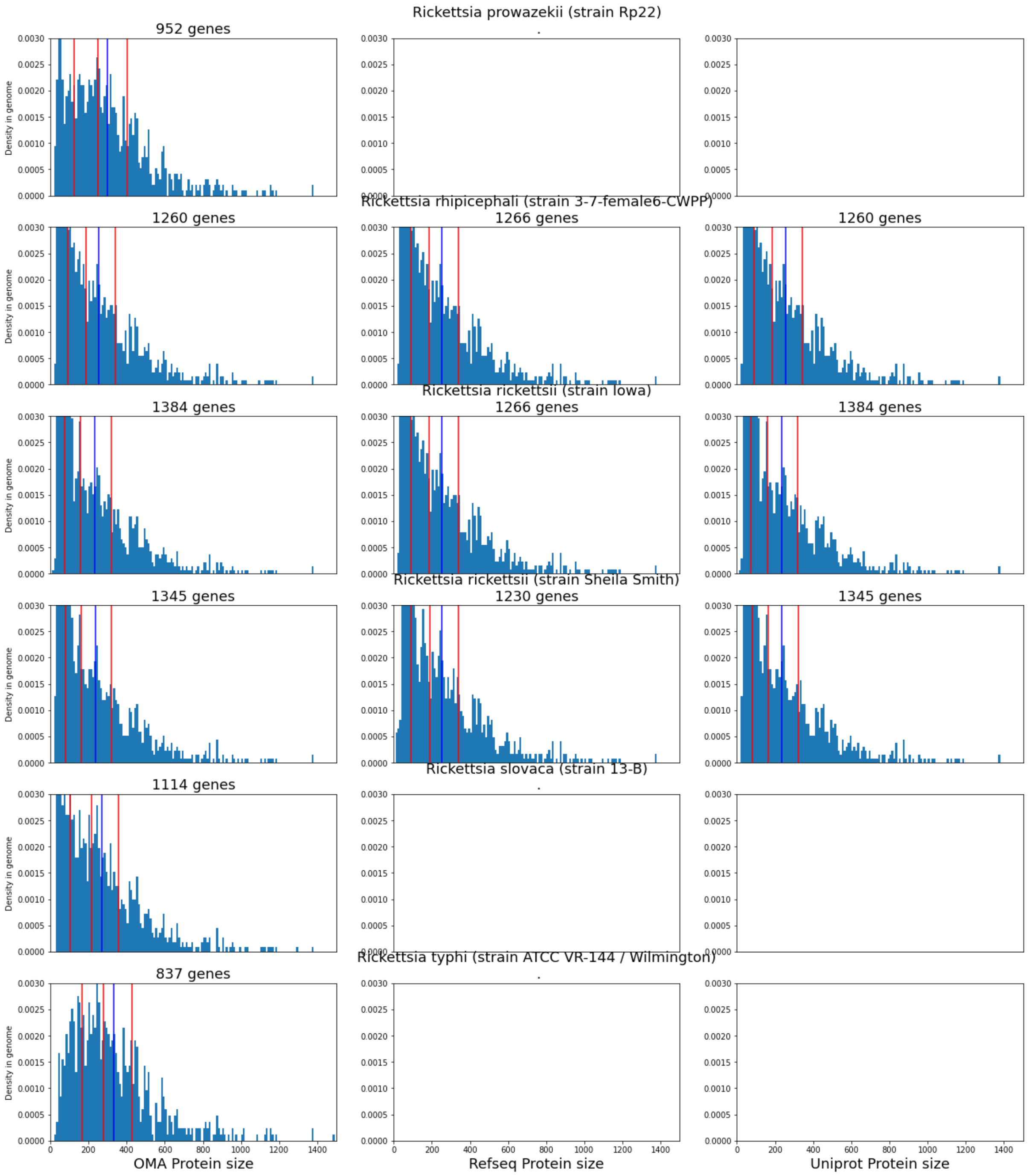
Rickettsia prowazekii (strain Rp22)
952 genes
Rickettsia rhipicephali (strain 3-7-female6-CWPP)
1260 genes
1266 genes
1260 genes
Rickettsia rickettsii (strain Iowa)
1384 genes
1266 genes
1384 genes
Rickettsia rickettsii (strain Sheila Smith)
1345 genes
1230 genes
1345 genes
Rickettsia slovaca (strain 13-B)
1114 genes
Rickettsia typhi (strain ATCC VR-144 / Wilmington)
837 genes
Density in genome
OMA Protein size
Refseq Protein size
Uniprot Protein size

Supplementary Figure 36 : **Protein length distributions in Apicomplexa proteomes**. Each plot represents the protein length distribution of the species proteome in OMA. Only length up to 2000 aa are represented. Species name is indicated as the top, with the number of proteins in the proteome. Red lines indicate, from left to right: 1st quartile, median and 3rd quartile of protein length. The blue line represents the mean.

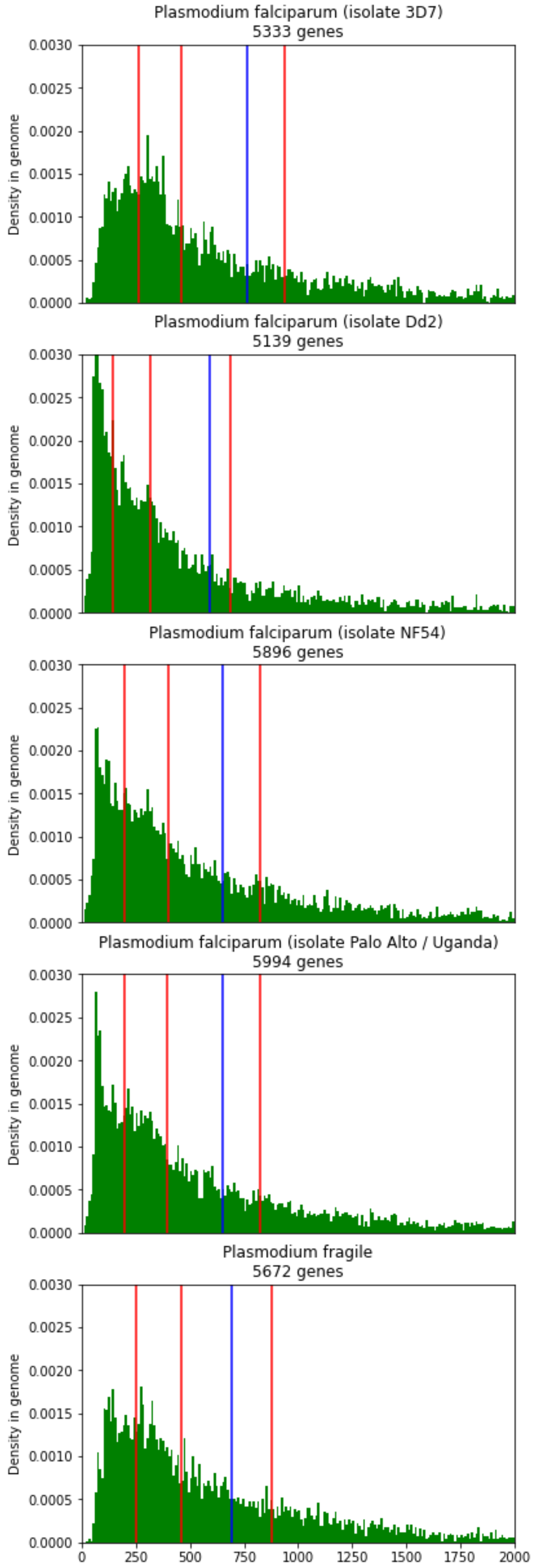

Plasmodium falciparum (isolate 3D7)
5333 genes
Plasmodium falciparum (isolate Dd2)
5139 genes
Plasmodium falciparum (isolate NF54)
5896 genes
Plasmodium falciparum (isolate Palo Alto / Uganda)
5994 genes
Plasmodium fragile
5672 genes
Density in genome
0.0000
0.0005
0.0010
0.0015
0.0020
0.0025
0.0030
0
250
500
750
1000
1250
1500
1750
2000

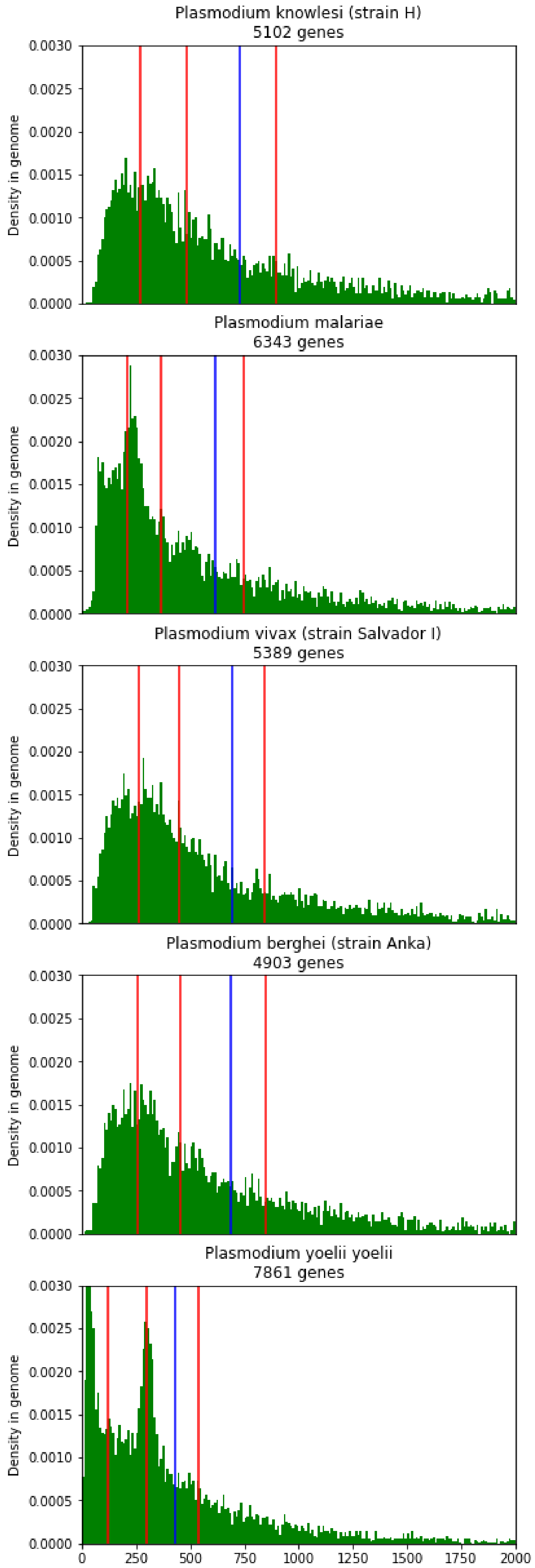
Plasmodium knowlesi (strain H)
5102 genes
Plasmodium malariae
6343 genes
Plasmodium vivax (strain Salvador I)
5389 genes
Plasmodium berghei (strain Anka)
4903 genes
Plasmodium yoelii yoelii
7861 genes
Density in genome
0.0000
0.0005
0.0010
0.0015
0.0020
0.0025
0.0030
0
250
500
750
1000
1250
1500
1750
2000

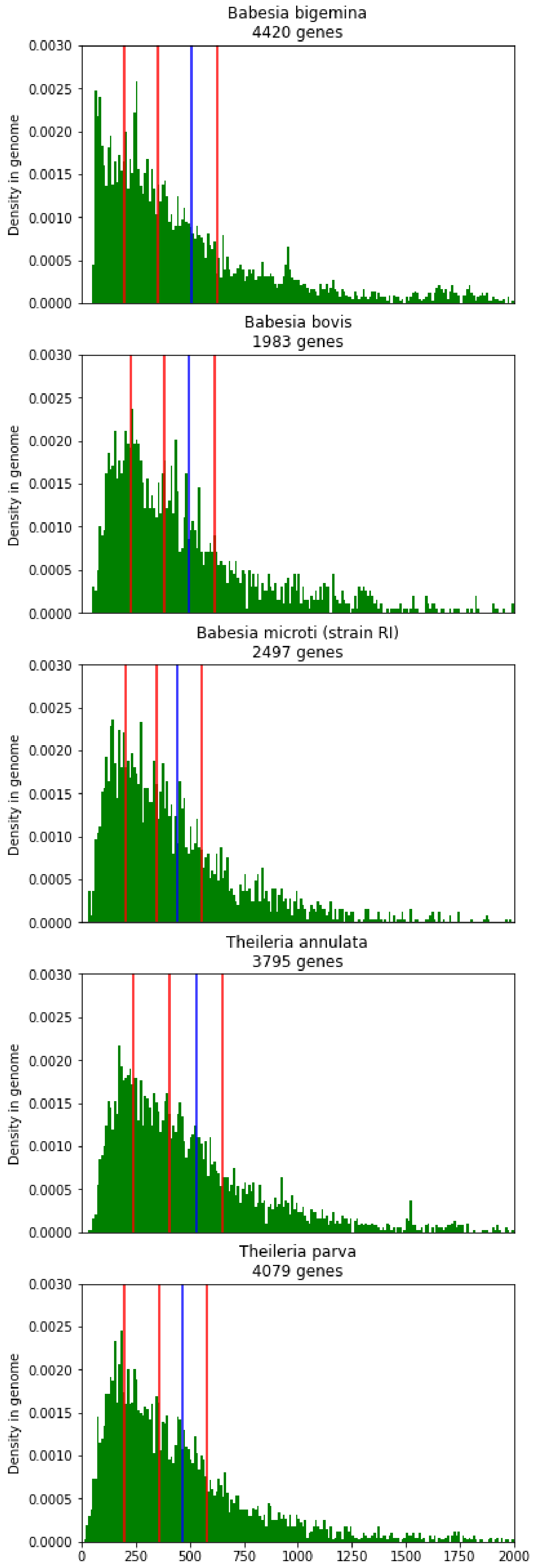
Babesia bigemina
4420 genes
Babesia bovis
1983 genes
Babesia microti (strain RI)
2497 genes
Theileria annulata
3795 genes
Theileria parva
4079 genes
Density in genome
0.0000
0.0005
0.0010
0.0015
0.0020
0.0025
0.0030
0
250
500
750
1000
1250
1500
1750
2000

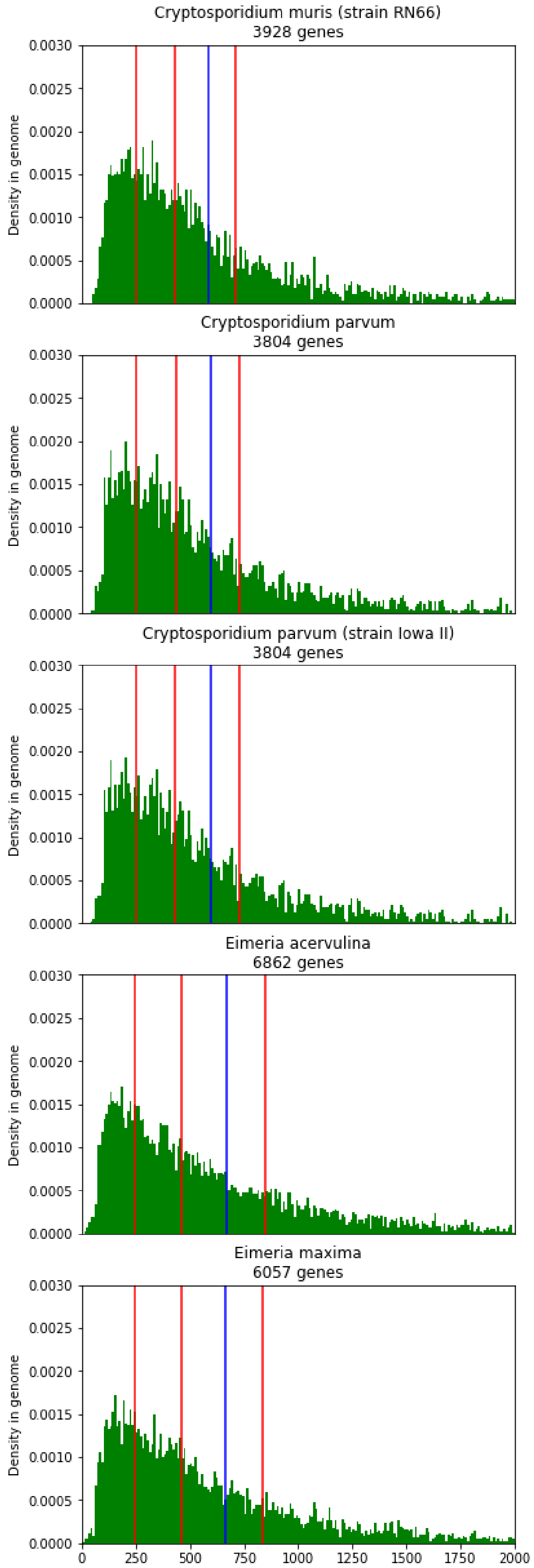

Cryptosporidium muris (strain RN66)
3928 genes
Cryptosporidium parvum
3804 genes
Cryptosporidium parvum (strain Iowa II)
3804 genes
Eimeria acervulina
6862 genes
Eimeria maxima
6057 genes
Density in genome
0.0000
0.0005
0.0010
0.0015
0.0020
0.0025
0.0030
0
250
500
750
1000
1250
1500
1750
2000

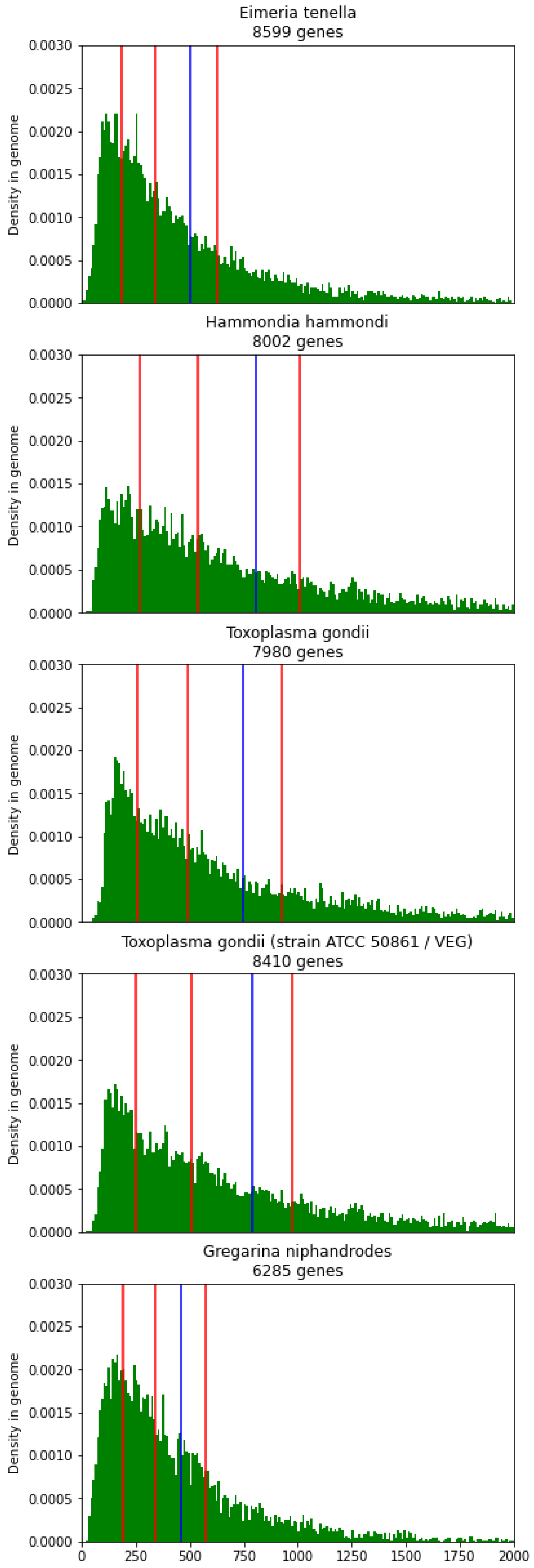

Eimeria tenella
8599 genes
Hammondia hammondi
8002 genes
Toxoplasma gondii
7980 genes
Toxoplasma gondii (strain ATCC 50861 / VEG)
8410 genes
Gregarina niphandrodes
6285 genes
Density in genome
0.0000
0.0005
0.0010
0.0015
0.0020
0.0025
0.0030
0
250
500
750
1000
1250
1500
1750
2000

Supplementary Figure 36: **Protein length distributions in proteomes from the *Ustilago* genus**. Each plot represents the protein length distribution of the species proteome in OMA. Only length up to 2000 aa are represented. Species name is indicated as the top, with the number of proteins in the proteome. Red lines indicate, from left to right : 1st quartile, median and 3rd quartile of protein length. The blue line represents the mean.

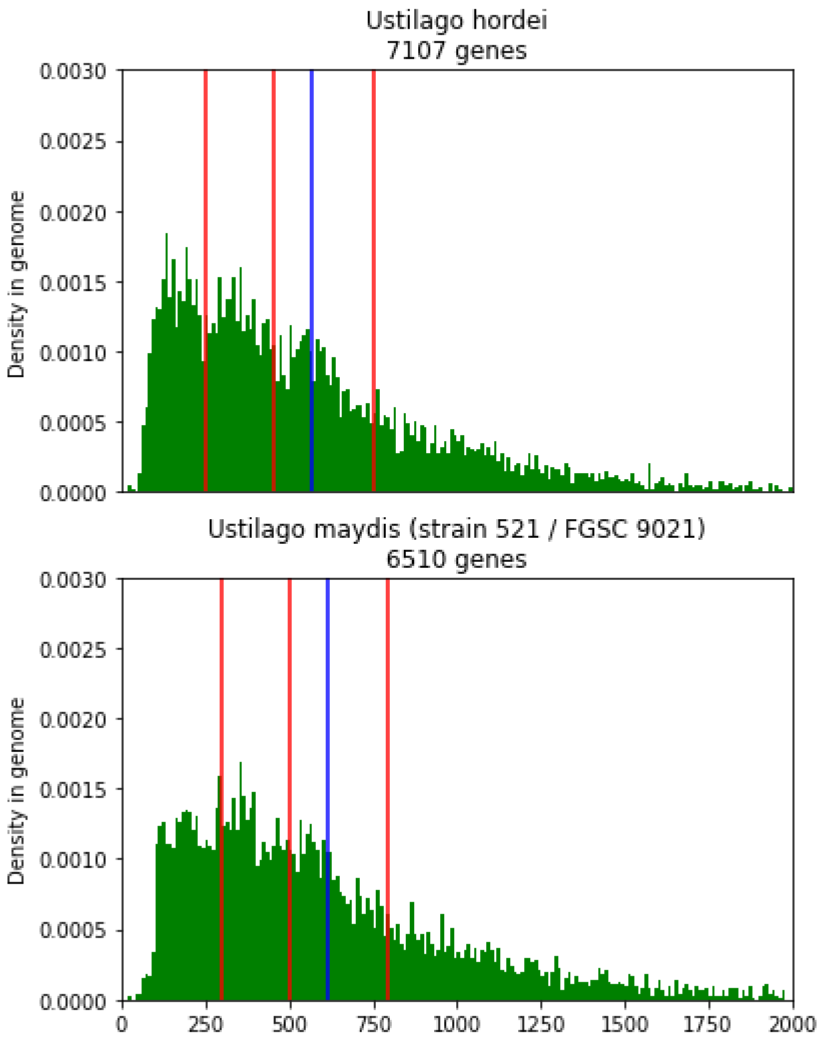

Supplementary Figure 37 : **Gene ontology enrichment of long genes for *Ustilago maydis***. Enrichment of genes larger than 1,000 aa for the different GO categories (top: BP- biological process, middle: MF - Molecular Function, bottom : CC - Cellular component), with the background set being either the whole gene repertoire of the species (left) or all the proteins in the dataset larger than 1,000 aa (right). Results are shown in semantic similarity scatterplots, which summarize the enriched GO terms by removing term redundancy. The p-values for the enriched terms are shown by color, and the number of terms each circle represents is shown by its size.

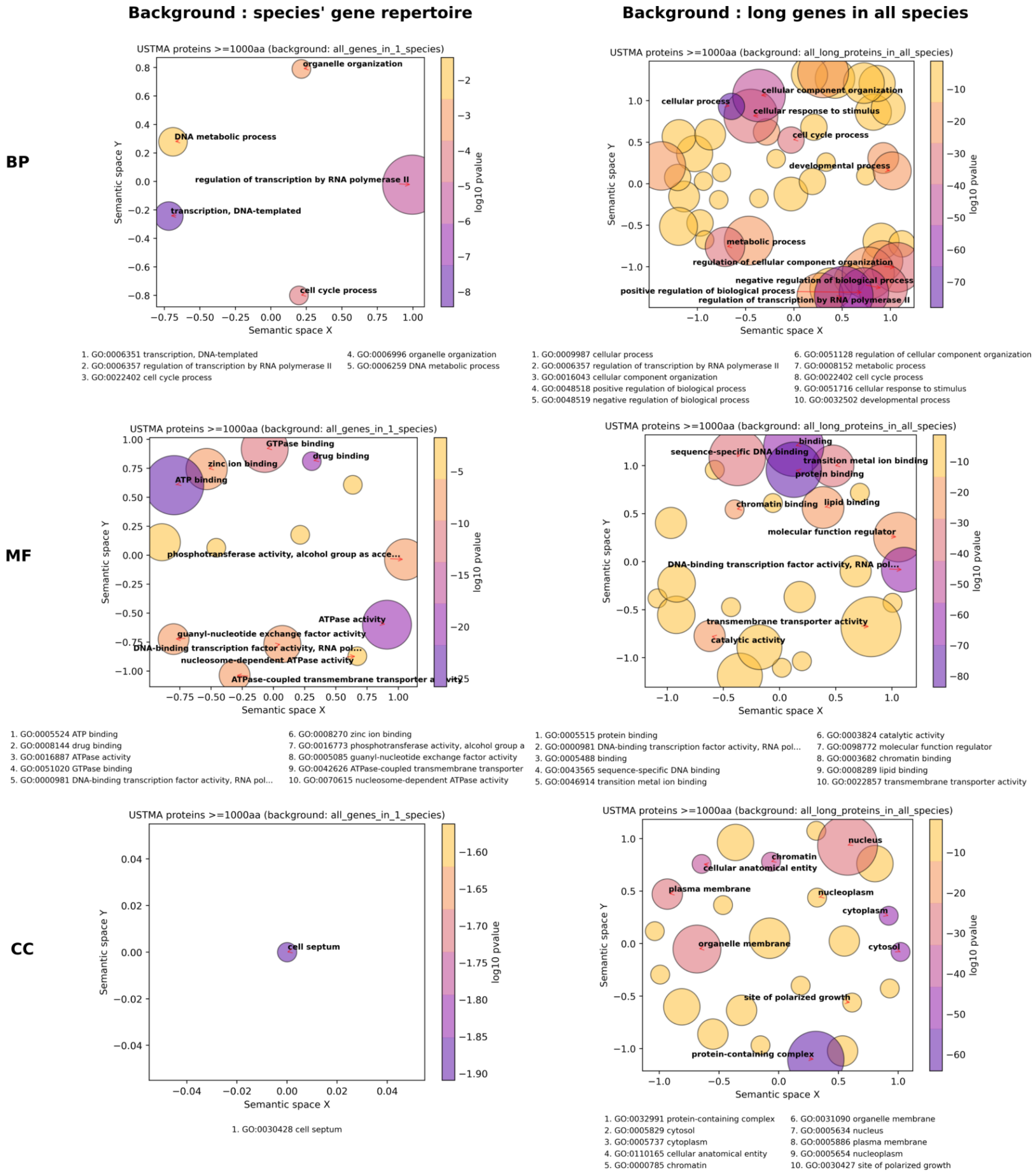

Supplementary Figure 38: **Gene ontology enrichment of long genes for *Toxoplasma gondii* (strain VEG).** Enrichment of genes larger than 1,000 aa for the different GO categories (top: BP- biological process, middle: MF - Molecular Function, bottom : CC - Cellular component), with the background set being either the whole gene repertoire of the species (left) or all the proteins in the dataset larger than 1,000 aa (right). Results are shown in semantic similarity scatterplots, which summarize the enriched GO terms by removing term redundancy. The p-values for the enriched terms are shown by color, and the number of terms each circle represents is shown by its size.

**Background : species' gene repertoire** | **Background : long genes in all species**

**BP**

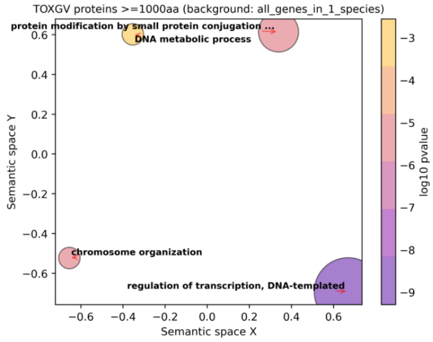


1. GO:0006355 regulation of transcription, DNA-templated
2. GO:0051276 chromosome organization
3. GO:0070647 protein modification by small protein conjugation ...
4. GO:0006259 DNA metabolic process

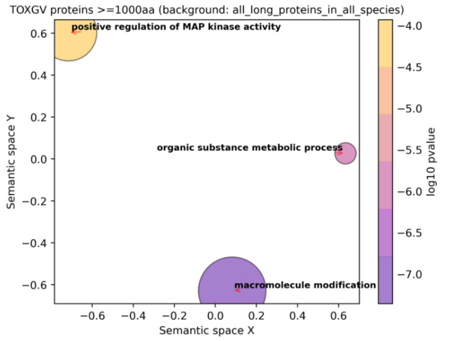


1. GO:0043412 macromolecule modification
2. GO:0071704 organic substance metabolic process
3. GO:0043406 positive regulation of MAP kinase activity

**MF**

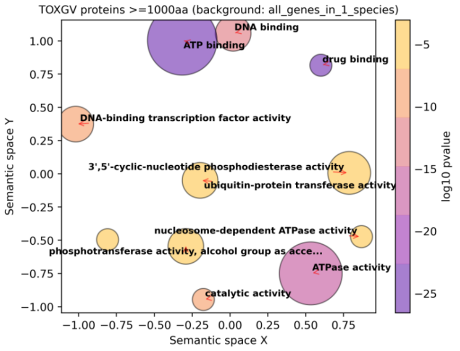


1. GO:0005524 ATP binding
2. GO:0008144 drug binding
3. GO:0016887 ATPase activity
4. GO:0003677 DNA binding
5. GO:0003700 DNA-binding transcription factor activity
6. GO:0003824 catalytic activity
7. GO:0070615 nucleosome-dependent ATPase activity
8. GO:0004842 ubiquitin-protein transferase activity
9. GO:0016773 phosphotransferase activity, alcohol group as acce..
10. GO:0004114 3',5'-cyclic-nucleotide phosphodiesterase activity

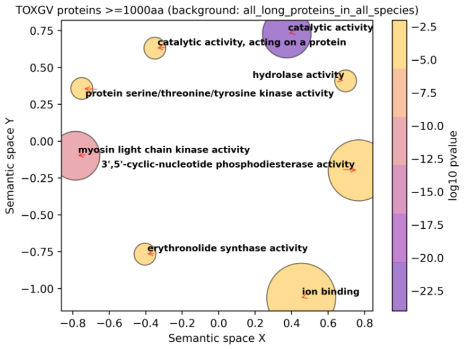


1. GO:0003824 catalytic activity
2. GO:0004687 myosin light chain kinase activity
3. GO:0016787 hydrolase activity
4. GO:0140096 catalytic activity, acting on a protein
5. GO:0043167 ion binding
6. GO:0004114 3',5'-cyclic-nucleotide phosphodiesterase activity
7. GO:0004712 protein serine/threonine/tyrosine kinase activity
8. GO:0047879 erythronolide synthase activity

**CC**

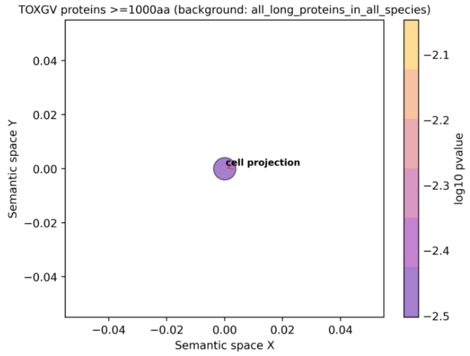


1. GO:0042995 cell projection

Supplementary Figure 39: **Gene ontology enrichment of long genes for *Plasmodium falciparum*** (isolate 3D7)**.** Enrichment of genes larger than 1,000 aa for the different GO categories (top: BP- biological process, middle: MF - Molecular Function, bottom : CC - Cellular component), with the background set being either the whole gene repertoire of the species (left) or all the proteins in the dataset larger than 1,000 aa (right). Results are shown in semantic similarity scatterplots, which summarize the enriched GO terms by removing term redundancy. The p-values for the enriched terms are shown by color, and the number of terms each circle represents is shown by its size.

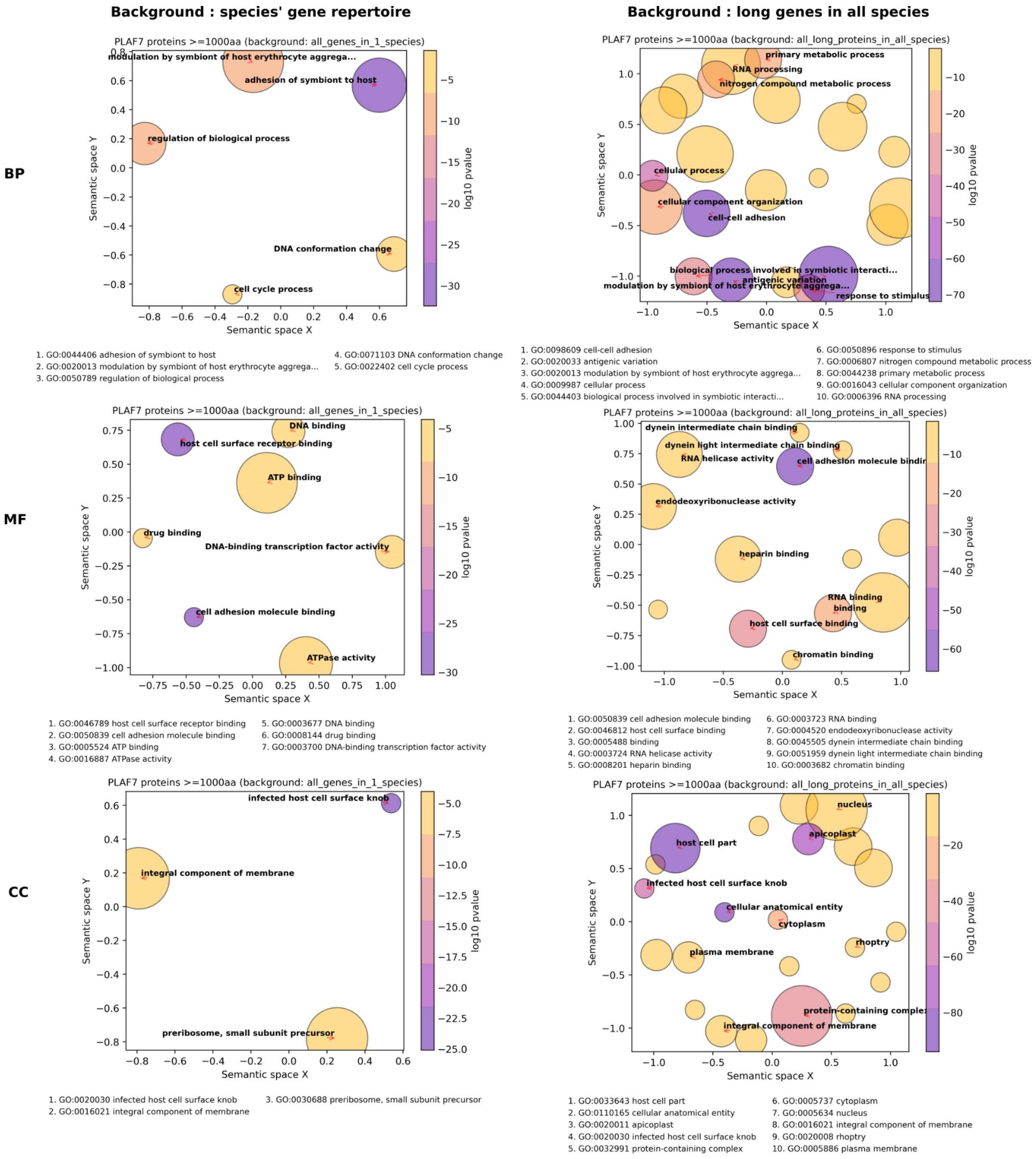
